# Species Distribution Modeling of *Aedes aegypti* in Maricopa County, Arizona from 2014-2020

**DOI:** 10.1101/2022.03.30.486464

**Authors:** Whitney M. Holeva-Eklund, Steven J. Young, James Will, Nicole Busser, John Townsend, Crystal M. Hepp

## Abstract

*Aedes aegypti* mosquitoes, which are of great importance to public health, have recently and rapidly expanded their global range; however, little is understood about the distribution of these mosquitoes in desert environments. Trapping records for *Ae. aegypti* in Maricopa County, Arizona from 2014-2020 were used to construct species distribution models using Maxent. Satellite imagery and socioeconomic factors were used as predictors. Maps of predicted habitat suitability were converted to binary presence/absence maps, and consensus maps were created that represent “core” habitat for the mosquito over six years of time. Generally, population density was the most important predictor in the models while median income was the least important. This study is the first step toward understanding the distribution of the vector in a desert environment.

**One-Sentence Summary:** This study uses satellite imagery and socioeconomic factors to identify core habitat over six years in a desert environment.

## Main Text

*Aedes aegypti* mosquitoes are vectors of great public health importance as they are capable of transmitting dengue, Zika, chikungunya, and yellow fever viruses (*1–5*). Importantly, the range of this vector has recently expanded rapidly throughout the world (*3*), putting an even greater number of people at risk of contracting these diseases. This vector lives in close proximity to human settlements, is active during daytime hours, and generally relies almost exclusively on humans for blood meals, making the mosquito an efficient vector of diseases (*6*). The mosquito has a very small range, typically considered to be less than 100 meters over the course of their lifetime (*7*), so their distribution is generally not homogeneous over an entire urban area but rather depends on habitat availability and microenvironment characteristics.

The goal of this study is to create quarterly species distribution models based on satellite imagery and socioeconomic indicators for *Ae. aegypti* in Maricopa County, Arizona from 2014-2020. Maricopa County is located in the Sonoran Desert, habitat that may not naturally be suitable for the mosquito vector. However, anthropogenic modifications to the environment have created an oasis in the desert that has allowed this mosquito vector to thrive in this area. The *Ae. aegypti* population in Maricopa County, Arizona has been increasing over the last decade, and the mosquito is now routinely trapped year-round in this region. This project will be a step toward the long-term objective of developing an understanding of the underlying geographic distribution and movement of the mosquito vector *Ae. aegypti* in Maricopa County, Arizona and, more broadly, in arid desert environments. In addition to contributing to the growing body of research that is attempting to understand the distribution of this vector, especially in territories into which the mosquito has recently expanded, the findings of this study will be beneficial to vector control and public health efforts as the mosquito population increases and if outbreaks of disease occur.

## Results

Trapping observations from 2014-2020 were split into 28 quarterly time periods (*8*), with full data available for only 25 quarters (Table 1). The number of observations varied for each quarter from a low of 73 locations (2/1/2014-4/30/2014) to a high of 778 locations (8/1/2014- 10/31/2014) (Table 1). Overall, for all 25 quarters for which data was available, there were 11,399 locations at which *Ae. aegypti* was observed, although the locations were not unique for each quarter. The mean number of locations at which *Ae. aegypti* was observed per quarter was 456 (standard deviation: 203).

**Table 1.** Quarters with descriptive statistics and results. This table shows the quarter number, dates, number of observations used in model building, average test AUC from Maxent output, the threshold used to convert the suitability output to binary presence/absence data using a 10 percent omission threshold, the percentage of are calculated as suitable after converting to binary data, and the most important predictor with the permutation importance.

| Quarter | Dates | Number of Obs. Used in Model | Avg. Test AUC | Threshold to Convert to Binary Map | Area Predicted as Suitable (%) | Most Important Predictor / Permutation Importance |
| --- | --- | --- | --- | --- | --- | --- |
| 1 | 2/1/14—4/30/14 | 73 | 0.942 | 0.52 | 9.99 | Pop. Dens. / 89.4 |
| 2 | 5/1/14—7/31/14 | 233 | 0.950 | 0.50 | 7.83 | Pop. Dens. / 82.4 |
| 3 | 8/1/14—10/31/14 | 778 | 0.900 | 0.68 | 12.09 | Pop. Dens. / 66.4 |
| 4 | 11/1/14—1/31/15 | 382 | 0.900 | 0.60 | 14.31 | Pop. Dens. / 62.1 |
| 5 | 2/1/15—4/30/15 | 326 | 0.928 | 0.58 | 10.81 | Pop. Dens. / 67.8 |
| 6 | 5/1/15—7/31/15 | 571 | 0.917 | 0.56 | 13.02 | Pop. Dens. / 76.8 |
| 7 | 8/1/15—10/31/15 | 587 | 0.913 | 0.54 | 11.17 | Pop. Dens. / 61.0 |
| 8 | 11/1/15—1/31/16 | 337 | 0.924 | 0.60 | 10.14 | Pop. Dens. / 70.5 |
| 9 | 2/1/16—4/30/16 | 247 | 0.933 | 0.64 | 8.97 | Pop. Dens. / 80.9 |
| 10 | 5/1/16—7/31/16 | 484 | 0.921 | 0.62 | 9.88 | NDMI / 59.5 |
| 11 | 8/1/16—10/31/16 | 702 | 0.915 | 0.56 | 10.75 | Pop. Dens. / 67.8 |
| 12 | 11/1/16—1/31/17 | 511 | 0.921 | 0.56 | 11.00 | Pop. Dens. / 68.9 |
| 13 | 2/1/17—4/30/17 | 299 | 0.933 | 0.54 | 11.50 | Pop. Dens. / 90.8 |
| 14 | 5/1/17—7/31/17 | 613 | 0.908 | 0.60 | 12.08 | Pop. Dens. / 55.0 |
| 15 | 8/1/17—10/31/17 | 763 | 0.908 | 0.62 | 11.17 | Pop. Dens. / 53.1 |
| 16 | 11/1/17—1/31/18 | 388 | 0.924 | 0.60 | 11.75 | Pop. Dens. / 57.8 |
| 17 | 2/1/18—4/30/18 | 121 | 0.942 | 0.60 | 10.51 | Pop. Dens. / 85.4 |
| 18 | 5/1/18—7/31/18 | 580 | 0.921 | 0.58 | 10.77 | Pop. Dens. / 71.0 |
| 19 | 8/1/18—10/31/18 | n/a | n/a | n/a | n/a | n/a |
| 20 | 11/1/18—1/31/19 | n/a | n/a | n/a | n/a | n/a |
| 21 | 2/1/19—4/30/19 | 304 | 0.920 | 0.66 | 11.23 | Pop. Dens. / 84.7 |
| 22 | 5/1/19—7/31/19 | 711 | 0.911 | 0.62 | 11.33 | NDMI / 57.1 |
| 23 | 8/1/19—10/31/19 | 705 | 0.900 | 0.58 | 13.17 | NDMI / 49.4 |
| 24 | 11/1/19—1/31/20 | 383 | 0.930 | 0.58 | 9.33 | Pop. Dens. / 77.8 |
| 25 | 2/1/20—4/30/20 | 210 | 0.919 | 0.52 | 11.57 | Pop. Dens. / 83.8 |
| 26 | 5/1/20—7/31/20 | 612 | 0.915 | 0.62 | 10.77 | NDMI / 47.5 |
| 27 | 8/1/20—10/31/20 | 479 | 0.925 | 0.66 | 9.43 | NDMI / 56.7 |
| 28 | 11/1/20—1/31/21 | n/a | n/a | n/a | n/a | n/a |

The following five predictors were used: normalized difference vegetation index (NDVI), normalized difference moisture index (NDMI), elevation, population density, and median income (*8*). The importance for each predictor varied from quarter to quarter, but population density was almost always the most important predictor (20/25 models) while median income (16/25 models) or elevation (9/25 models) was the least important predictor for all quarters. NDMI was the most important predictor for five quarters: quarters 10, 22, 23, 26, and 27.

For each quarter, Maxent output was produced using 10-fold cross validation (*8*). Output included an image showing the pixel-by-pixel habitat suitability value on a scale of 0-1 (with 1 being highest suitability) (Fig. S1-S25), response curves for each predictor (Fig. S37-S61), the value for the average permutation importance for each predictor across all folds (*9, 10*), a receiver operating characteristic (ROC) curve, and test area under curve (AUC) value (*11*) which provides a threshold-independent approach to evaluating model performance (Table 1). All quarterly models had high (≥ 0.90) test AUC, indicating excellent model fit (*11*) (Table 1). Suitability was generally high in the Phoenix metropolitan area and lower in the rural areas of the county; however, there did exist quite a bit of variation in suitability throughout the Phoenix area, with the eastern and southeastern portions of the metropolitan area generally having the highest predicted suitability.

Consensus maps were created for each year (e.g., year 1 was an aggregate of quarter 1, quarter 2, quarter 3, and quarter 4) and for each season (e.g., all February-April quarters; all May-July quarters; all August-October quarters; and all November-January quarters) by averaging the predicted suitability by pixel (Fig. S26-S36; *8*). Additionally, an overall consensus map was created by averaging the predicted suitability values for all 25 quarterly maps (Fig. 1; *8*).

**Fig 1.**
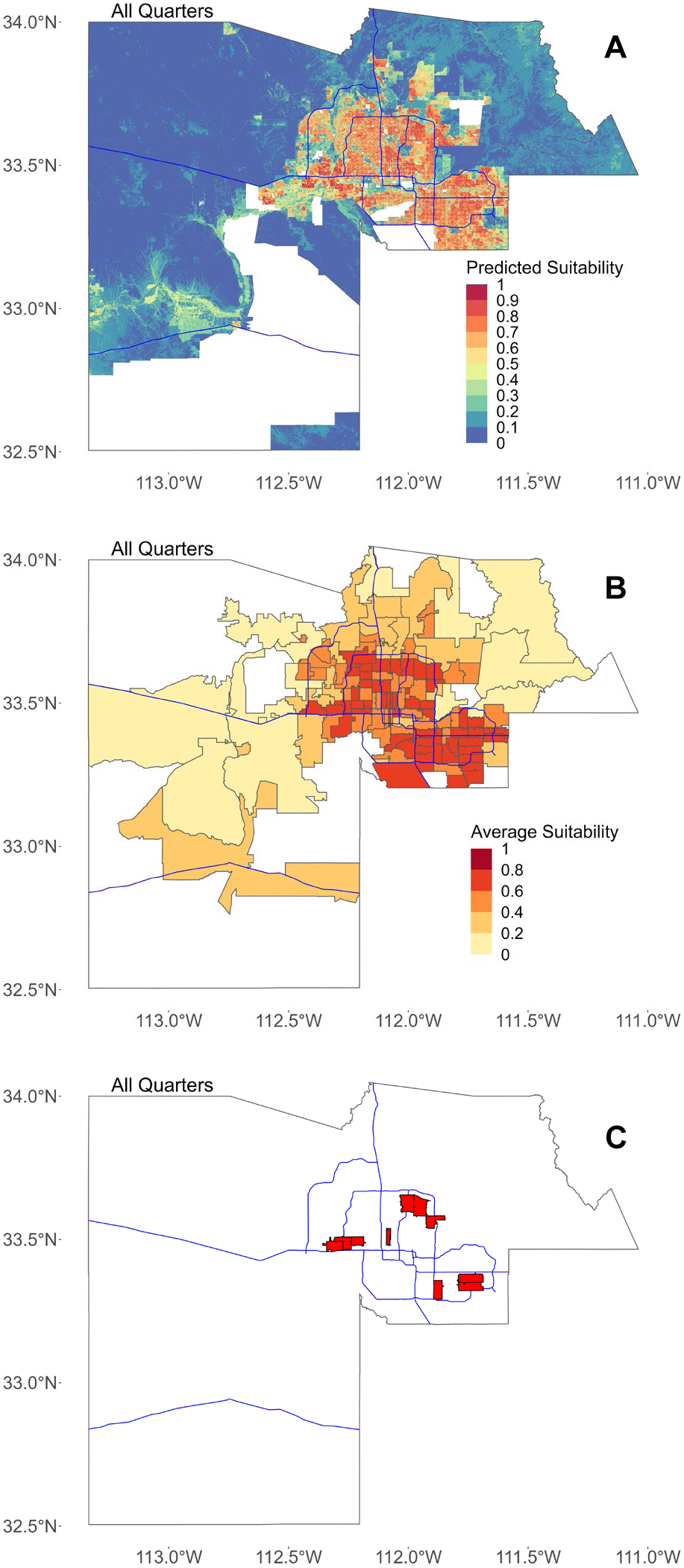
Average predicted suitability for all quarters. In all figures, blue lines represent major freeways in Maricopa County as a spatial reference. (**A**) Map showing the mean predicted suitability for each 30-meter by 30-meter pixel, calculated as the mean of the predicted suitability for all 25 quarters. Areas with missing data where suitability was not predicted are shown in white. (**B**) Predicted suitability for all quarters averaged by ZCTA. (**C**) The location of ten ZCTAs with the highest average suitability for all quarters.

The suitability maps were converted to binary presence/absence maps using the 10 percent omission threshold for each quarter (Table 1; Fig. S1-S25; *8*). After the suitability maps were converted to binary values, we calculated the percentage of area in the county that was deemed suitable for each quarter (Table 1; *8*). Further, consensus maps were created by overlaying the binary maps for each year, each season, and for all maps as described above (Fig. 2; Fig. S26-36; *8*). Areas that were predicted as consistently suitable in each of the quarterly maps were considered suitable for the consensus maps (*12*). For the consensus maps, the percent of area predicted as suitable and the omission rate was calculated (Table 2; *8*).

**Fig 2.**
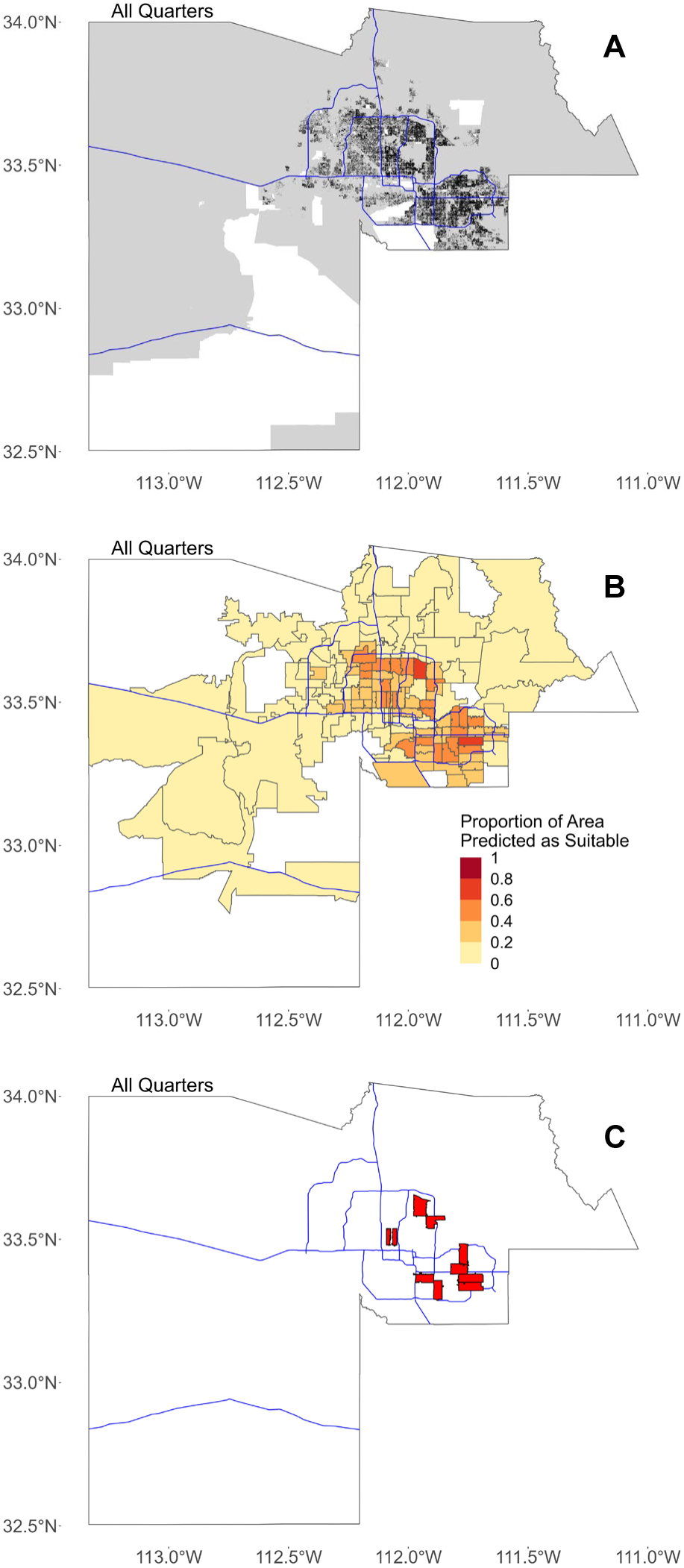
Binary maps for all quarters. In all figures, blue lines represent major freeways in Maricopa County as a spatial reference. (**A**) The consensus binary map for all quarters created by overlaying the binary map for all 25 quarters. Areas with missing data where suitability was not predicted are shown in white. (**B**) Proportion of area in each ZCTA predicted as suitable based on binary map in (A). (**C**) The location of ten ZCTAs with the highest proportion of area predicted as suitable for all quarters.

**Table 2.**
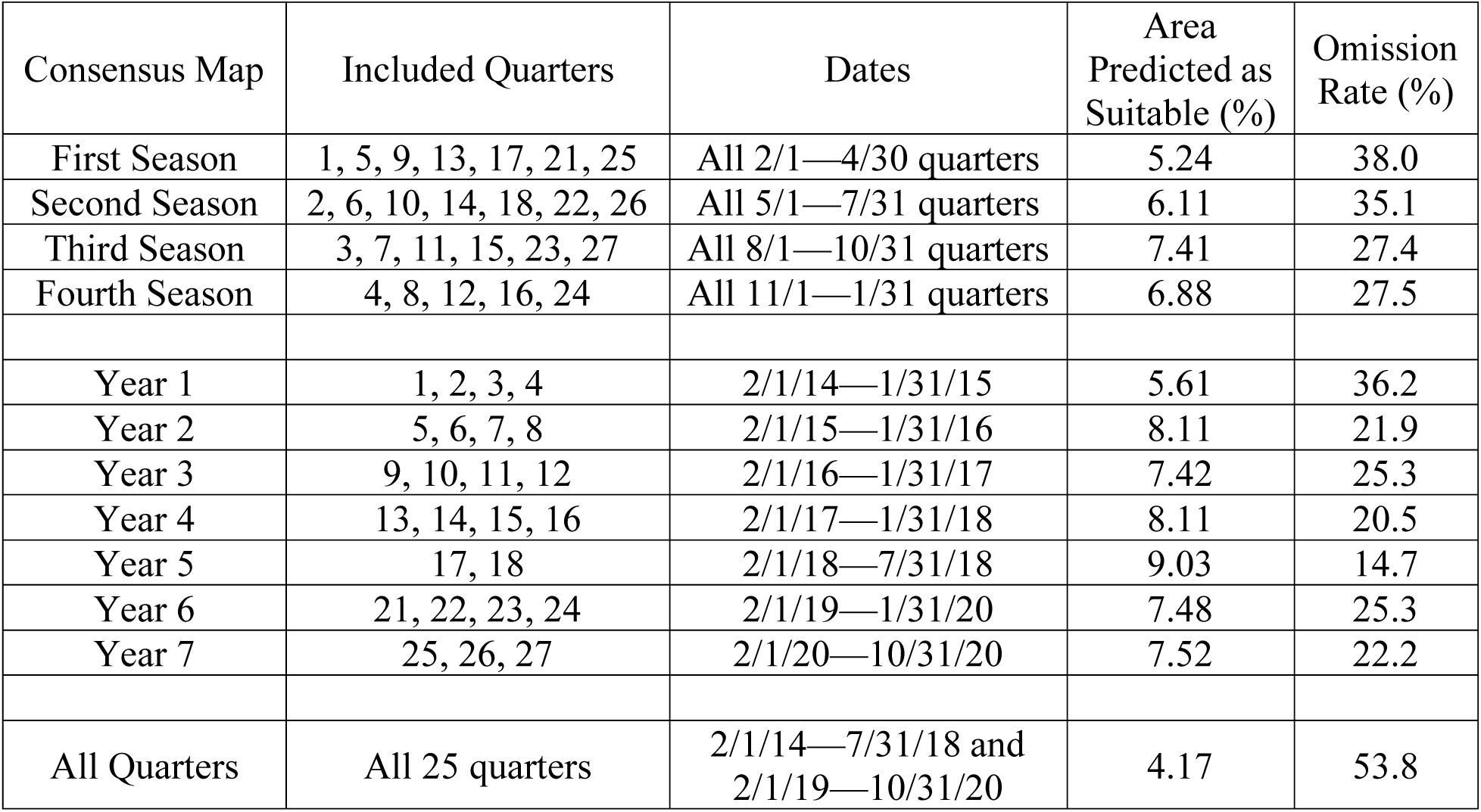
Consensus map aggregations with descriptive statistics and results. This table shows the consensus aggregation with the quarters used for each, the dates for each consensus map, the percentage of area predicted as suitable after converting to a binary map, and the omission rate of the binary map.

We also aggregated the output for census block groups, census tracts, and census zip code tabulation areas (ZCTA) in two different ways in order to produce more actionable maps for vector control (*8*), since individual 30-meter by 30-meter pixels would be difficult to target for mosquito control. First, we used the habitat suitability (on a scale of 0-1) to determine the average suitability for the block groups, tracts, and ZCTAs. We produced these average maps for each quarter and again for all consensus average predicted suitability maps (Fig. 1; Fig. S1-S36; *8*). The top 10 block groups, tracts, and ZCTAs in terms of overall average suitability were spread throughout the Phoenix metropolitan area in the southeastern, northeastern, central, and southwestern areas (Fig. 1). Second, using the quarterly and consensus binary maps, we determined the proportion of area that was predicted to be suitable within the block groups, tracts, and ZCTAs for each quarter and for all consensus binary maps (Fig. 2; Fig. S1-S36; *8*). All of the top 10 block groups, tracts, and ZCTAs in terms of overall average predicted suitability are located in the central Phoenix or the eastern/southeastern Phoenix metropolitan area (Fig. 2).

## Discussion

Through the creation of species distribution models for 25 quarterly time periods over six years, we were able to identify potential long term “core” breeding habitat for the mosquito vector, as well as some “hotspot” locations with greater than average suitability that could potentially be targeted for vector control activity. One of the greatest strengths of this study is the ability to look at seasonal trends over the course of six years, providing further insight into the long-term breeding habits of the vector in this region.

Binary maps, especially the consensus binary maps, are potentially great tools for vector control planning purposes. Distilling six years of trapping data down into one map that shows locations with consistent habitat suitability for *Ae. aegypti* populations can help to provide an overall summary of the activity of *Ae. aegypti* since it has become strongly established in this region. It also helps to “fill in the gaps” between trapping locations and to identify additional areas that may have high suitability for the vector but are not currently being as intensively trapped or observed. We found that the omission rate was much higher for the consensus maps than it was for quarterly maps, indicating that there is a consistent area that is suitable for the vector, but that a lot of mosquito activity is also observed outside of this core area. It is unclear if the mosquito is breeding in the core areas and then migrating to additional areas or if breeding habitat varies from year to year. Future studies should examine the movement of the mosquito in this region over time which would provide another clue as to how the mosquito is surviving and behaving in a desert region.

Tract and ZCTA aggregations should be interpreted with caution since the census data for population density and median income were both collected at the block group level. Some tracts and ZCTAs have missing data due to 50 census block groups having missing median income information. Further, the census tracts and ZCTAs may not have the same composition as the block groups that made up the predictors, but instead are an aggregate of all characteristics in that area. In particular, the ZCTA results may be skewed because ZCTAs are not meant to represent a homogenous population. However, the results are still useful for vector control purposes because they provide an easier tool for planning instead of individual 30-meter by 30-meter pixels, and in the event of an outbreak suitability can quickly be linked to human case data which is generally reported at the ZCTA level. Importantly, the overall characteristics of the tract or ZCTA may not reflect the smaller areas that are actually suitable for the vector.

One limitation of this study was that trapping data was represented only as presence data. This means that outputs that represent suitability do not account for differences in abundance throughout the county (*13*). Some areas that are predicted to be suitable may only have small or transient populations of *Ae. aegypti* whereas others may have large, consistent populations of the vector and therefore a large increase in the potential risk of disease transmission by the mosquito. Presence only data also may not account for fluctuations in populations due to interventions such as insecticide application, which were not accounted for in this study.

Future studies that explore species distribution models that account for abundance could be helpful to more specifically identify areas that have large populations of *Ae. aegypti* where people may be more likely to be exposed to disease. Additionally, models that explicitly account for interventions could help to evaluate the effectiveness of these interventions.

Another limitation of this study was the lack of weather or climate predictors. We made the decision to exclude these predictors due to the relatively small study area that should have fairly consistent weather and climate and also due to the lack of availability of this type of data at the scale of our other predictors (30 meters). If this study is expanded to larger regions with variable climate, it will be important to incorporate predictors such as temperature and rainfall.

Although we can identify areas that have higher average suitability for *Ae. aegypti* populations than other areas in Maricopa County, it remains unclear whether these areas have higher suitability because they represent good breeding habitat or because adult females are attracted to these areas. In other words, it is unclear if the mosquito is spending its entire life cycle in these areas or if the adult female mosquitoes are attracted to these areas from the site of their origination. It has generally been thought that *Ae. aegypti* has a very small flight range, generally less than 150 meters over the course of its lifetime (*14*). However, other studies demonstrate that the mosquito can fly much further (*15*). As of yet, it is unclear how the mosquito is behaving in a desert habitat. Again, future studies that examine the movement of the vector in this region will be helpful to help answer these questions.

As of now, we cannot determine whether the findings presented in this study for highly suitable areas would be correlated with risk of disease in the event of an outbreak, although findings by Arboleda et al. indicate that areas predicted as suitable by a species distribution model are correlated with human cases of dengue (*12*). In the event of a future outbreak, it will be important to evaluate the relationship between our suitability maps and the incidence of human cases of disease; however, past studies suggest that the maps produced by this study could be helpful for a public health response during an outbreak of disease vectored by *Ae. aegypti*. It is our hope that preparing this information ahead of time could help to reduce the burden and cost of disease in Maricopa County if an outbreak does occur.

## Acknowledgments

Research reported in this publication was supported by the National Institute On Minority Health And Health Disparities of the National Institutes of Health under Award Number F31MD015674. The content is solely the responsibility of the authors and does not necessarily represent the official views of the National Institutes of Health.

## Author contributions

Conceptualization: WHE, CH

Methodology: WHE

Investigation: WHE, CH

Visualization: WHE

Funding acquisition: WHE, CH

Project administration: WHE, CH

Supervision: CH

Writing – original draft: WHE

Writing – review & editing: WHE, CH, SY, JW, NB, JT

## Data and materials availability

All data and code used in this analysis is available in a public Google Drive at https://drive.google.com/drive/folders/1Hrqs-Vpv-9w64rtBBXuKq9Po51D3A3-G?usp=sharing. View the File Descriptions, File Structure, and File Workflow in the README folder.

## Supplementary Materials

**Other Supplementary Materials for this manuscript include the following:**

All data and code used in this analysis is available in a public Google Drive at https://drive.google.com/drive/folders/1Hrqs-Vpv-9w64rtBBXuKq9Po51D3A3-G?usp=sharing.

### Materials and Methods

#### Maricopa County, Arizona: Study Location and Climate

Maricopa County is located in south central Arizona in the southwestern United States. The county comprises approximately 24,000 square kilometers of land and contains Arizona’s capital and most populous city, Phoenix, along with over 20 additional recognized cities and towns (*16*). The population of Maricopa County is approximately 4.5 million residents (*16*). The county is located in a desert climate with generally low humidity. Phoenix has an average annual rainfall of 20.4 cm, an average maximum temperature of 41.2°C for the hottest moth, July, and an average minimum temperature of 7.1°C for the coldest month, December (*17*). Rainfall is scarce and sporadic for most of the year, while monsoon season in July and August can bring torrential rains (*17*). Like most desert climates, the temperature can fluctuate drastically between the daytime high and the nighttime low with variations of 15°C or more in one day being common (*17*).

#### Trapping Data and Data Organization

Maricopa County Environmental Services Vector Control Division performs regular CO2 trapping throughout Maricopa County, but with traps concentrated in the Phoenix metropolitan area, by setting up more than 800 overnight traps for a 12-hour period each week. The trapping program in Maricopa County is one of the most comprehensive programs in the country, both in the amount of area covered and in the regularity with which traps are placed. Through an established partnership with Maricopa County Vector Control, we gained access to this trapping data for the years 2014-2020. The provided data includes the trapping date, number of adult mosquitoes trapped, species and sex of the mosquitoes, identification number of the trap, and latitude and longitude values.

To create the multitemporal models for this study, we divided the study period into quarters that roughly reflect four seasons of the year with varying levels of mosquito activity. The quarters were defined as follows: February 1-April 30, May 1-July 31, August 1-October 31, and November 1-January 31 (Table 1). To suit the requirements of Maxent, the species distribution model used for this study, trapping data were converted to presence only records for each quarter by creating a list of observations at unique locations defined by latitude and longitude and then removing duplicate observations for each location in each quarter. Due to missing data for 3 quarters (8/1/2018-10/31/2018, 11/1/2018-1/31/2019, and 11/1/2020- 1/31/2021), presence only records were available to create 25 quarterly models from 2014-2020.

#### Remote Sensing Predictors

Google Earth Engine (*18*) was used to download Landsat 8 OLI satellite imagery (*19*) clipped to Maricopa County, Arizona. For each quarter, all available Landsat 8 images with top of atmosphere reflectance were downloaded. Then, any pixels with a cloud score of 1 or cloud shadow confidence score of 3 were masked. For each Landsat 8 image available, the Normalized Difference Vegetation Index (NDVI) and the Normalized Difference Moisture Index (NDMI) were calculated using bands 5 and 4 or bands 5 and 6, respectively. Finally, the mean NDVI and mean NDMI values were calculated for each pixel using all Landsat 8 images in that quarter.

Landsat 8 imagery has a resolution of 30 meters, so the resulting NDVI and NDMI products for each quarter in turn also had a 30-meter resolution. Elevation data from the NASA Shuttle Radar Topography Mission (SRTM) was also clipped to Maricopa County, Arizona in Google Earth Engine (*20*). The images were exported from Google Earth Engine to be used with Maxent. The resample function from the raster package (Version 3.5-11) was used to ensure that the elevation raster matched the extent and cell size of the NDVI and NDMI rasters exactly (*21*).

#### Socioeconomic Predictors

Socioeconomic predictors were also included due to the close relationship that *Ae. aegypti* mosquitoes have with humans. Predictors were chosen based on a priori assumptions about potential influences on the suitability of habitat and the availability of blood meals for the vector. Socioeconomic predictors were downloaded from the United States Census TIGER/Line Shapefiles with Selected Demographic and Economic Data from the American Community Survey 5-Year Estimates from 2019 at the block group resolution (*22*). Data were downloaded for all of Arizona, so only those block groups that have their centroid in Maricopa County were selected for further use using ArcMap (Version 10.7.1).

Socioeconomic predictors of interest were population density and median income. Population density was calculated by dividing the population of each block group by the area of each block group in square kilometers. The polygon layers for these two variables of interest were converted to rasters using the Polygon to Raster tool in ArcMap (Version 10.7.1) using the setting MAXIMUM_AREA; extent and cell size was set to match that of the NDVI rasters downloaded from Google Earth Engine. The rasters for population density and median income were exported from ArcMap (Version 10.7.1) in ASCII format. Before continuing with further analysis, the socioeconomic predictor rasters were imported to RStudio (Version 1.4.1717) along with the NDVI, NDMI, and elevation rasters (*23*). The resample function from the raster package (Version 3.5-11) was used to ensure that the socioeconomic rasters matched the extent and cell size of the NDVI, NDMI, and elevation rasters exactly (*21*).

#### Modeling/Maxent

The first step in the modeling process involved evaluating the correlation of the five rasters (NDVI, NDMI, elevation, population density, median income) from each quarter to ensure that no rasters had a correlation of > 0.7. No rasters needed to be removed in this step, as the correlation between all four rasters was < 0.7 for every quarter. Next, we determined the best settings with which to run Maxent for each quarter individually using the ENMeval package (Version 2.0.2) in RStudio (Version 1.4.1717) using 10-fold cross validation (*23, 24*). The possible settings from which we chose the best model included the features L, LQ, H, LQH, LQHP, LQHPT (L = Linear; Q = Quadratic; P = Product; H = Hinge; T = Threshold) with a regularization multiplier between 1-5 (*10, 25*). The best settings for each quarter were chosen based on the model with the lowest delta AICc. Using the ENMeval package, a bias file was also created based on the observations for each quarter. Bias files help address the issues that arise when sampling is not random throughout the study area by biasing the background points selected to be in the same area where the majority of observations occurred, thereby reducing potential bias towards predicting higher suitability in areas that are simply more accessible and therefore more sampled. This approach is preferred over filtering observations when the species is expected to be spatially clumped, as is the case for *Ae. aegypti*, which would be expected to have greater populations around densely populated areas (*26*).

Next, we ran Maxent (version 3.4.0) for each quarter using the bias file created for each quarter and the best settings possible as described above. Each Maxent model was created using 10-fold cross validation, and the average output across all 10 folds was further evaluated. See Table S1 for the feature settings and regularization multiplier used for each quarter.

#### Output Manipulation

The Maxent output included habitat suitability maps for Maricopa County with values ranging from 0 to 1 representing the estimated habitat suitability for *Ae. aegypti* mosquitoes for each 30-meter by 30-meter square in the county, based on the observations for that quarter (*10*). These maps were then converted to binary presence/absence maps based on the 10 percent omission threshold determined using the package SDMtools (Version 1.1-221.2) in RStudio (Version 1.4.1717) (*23, 27*). After the suitability maps were converted to binary values, we calculated the percentage of area in the county that was deemed suitable for each quarter by dividing the number of pixels deemed suitable by the total number of pixels in the study area.

Further, consensus binary maps were created by overlaying the binary maps for each year and season as well as overall for all quarters. Maps of major freeways were downloaded from the AZGeo Data Hub to use as location references for the maps produced in this study. The data for the freeways maps was provided by the Maricopa Association of Governments and is publicly available and free to download as a shapefile (https://azgeo-data-hub-agic.hub.arcgis.com/).

Using the binary maps, we were able to calculate the percentage of area in Maricopa County that was determined to be suitable for Ae. aegypti for each quarter as well as for all of the consensus maps. We were also able to aggregate the results for block groups, tracts, and ZCTAs in two different ways. First, using the original habitat suitability output (with values ranging 0-1), we were also able to determine the average suitability for the aggregate areas for each quarter and for the consensus maps. Second, using the quarterly and consensus binary maps, we are able to determine the proportion of area that was predicted to be suitable within block groups, tracts, and ZCTAs. The 2019 shapefiles for the various census areas were downloaded from the United States Census Bureau (https://www.census.gov/geographies/mapping-files/time-series/geo/tiger-line-file.2019.html), then imported to RStudio (Version 1.4.1717) where we used the function ‘zonal’ from the raster (Version 3.5-11) package to determine the mean suitability value and the proportion of area predicted as suitable for each aggregate area. Maps were then created using ggplot2 (Version 3.3.5) and the sf (Version 1.0-5) packages or using the raster (Version 3.5-11) package (*21, 23*, *28, 29*).

**Figure S1.**
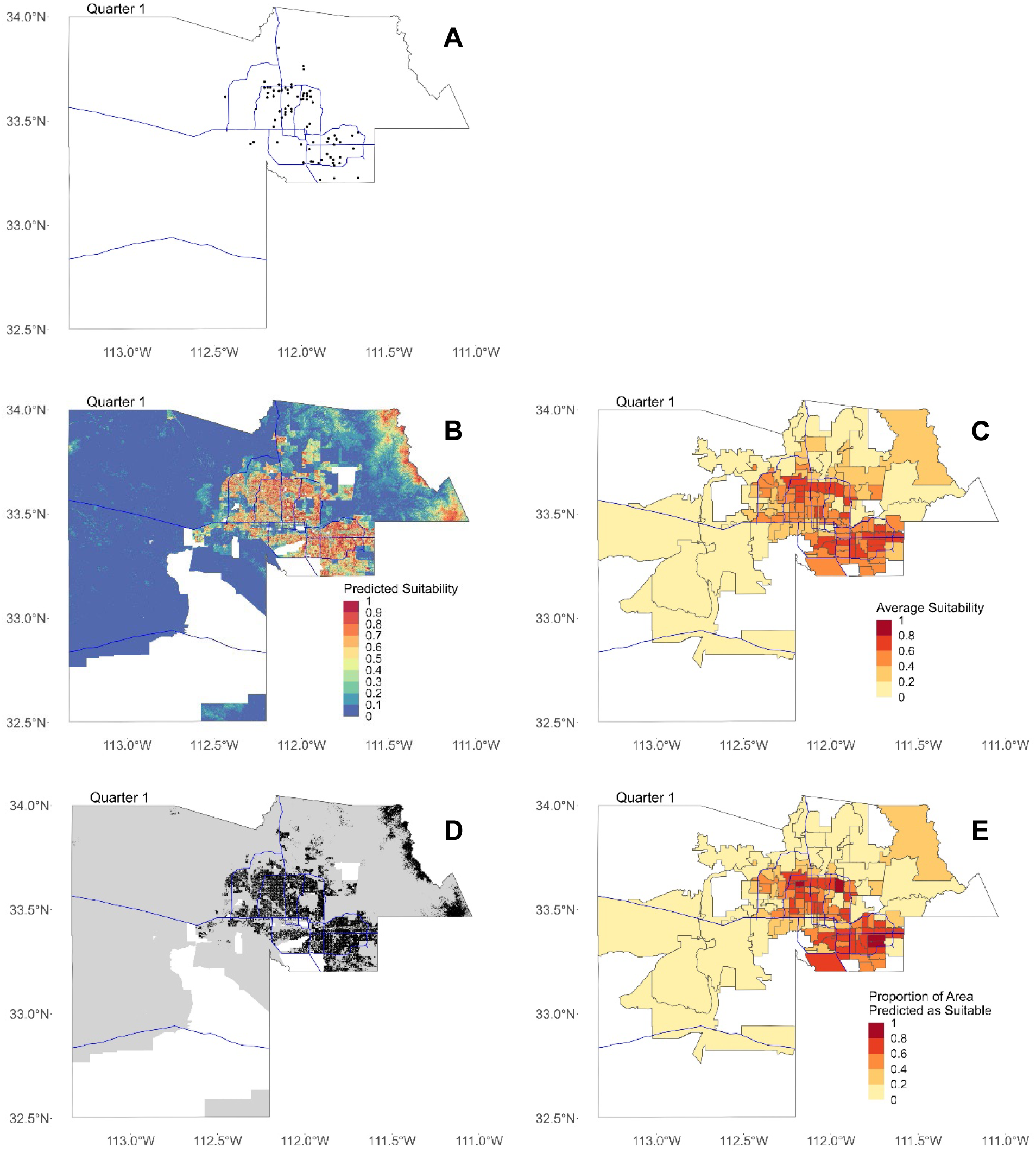
Maps for Quarter 1 (2/1/14—4/30/14). In all figures, blue lines represent major freeways in Maricopa County as a spatial reference. (**A**) Map of all locations where Ae. aegypti females were trapped in Maricopa County in Quarter 1. (**B**) Maxent output showing predicted suitability for 30-meter by 30-meter pixels. (**C**) Predicted suitability averaged by ZCTA. (**D**) Binary map showing areas predicted as suitable in black and not suitable in gray. In (B) and (C), areas with missing data where suitability was not predicted are shown in white. (**E**) Proportion of area in each ZCTA predicted as suitable after converting to binary map.

**Figure S2.**
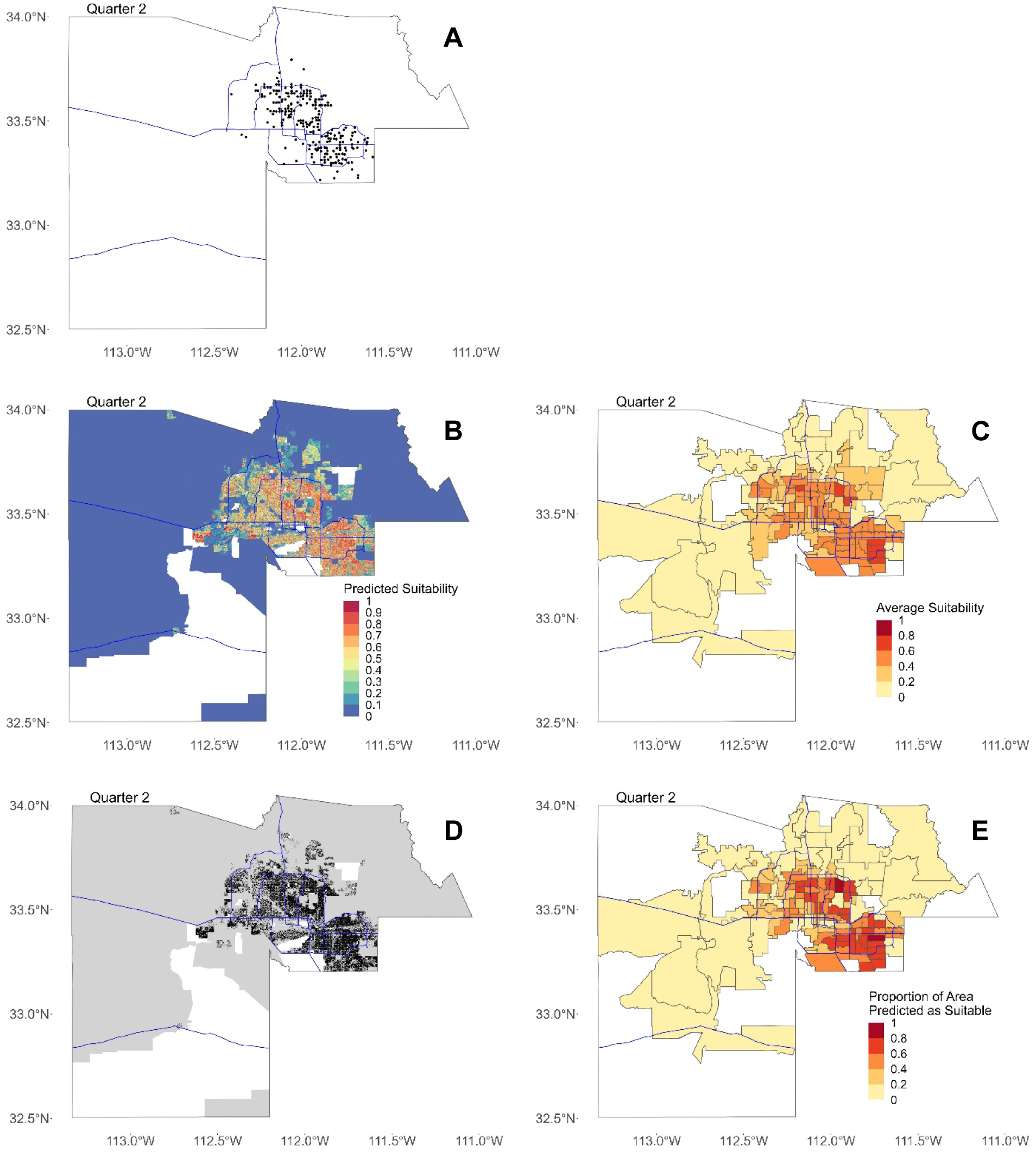
Maps for Quarter 2 (5/1/14—7/31/14). In all figures, blue lines represent major freeways in Maricopa County as a spatial reference. (**A**) Map of all locations where Ae. aegypti females were trapped in Maricopa County in Quarter 2. (**B**) Maxent output showing predicted suitability for 30-meter by 30-meter pixels. (**C**) Predicted suitability averaged by ZCTA. (**D**) Binary map showing areas predicted as suitable in black and not suitable in gray. In (B) and (C), areas with missing data where suitability was not predicted are shown in white. (**E**) Proportion of area in each ZCTA predicted as suitable after converting to binary map.

**Figure S3.**
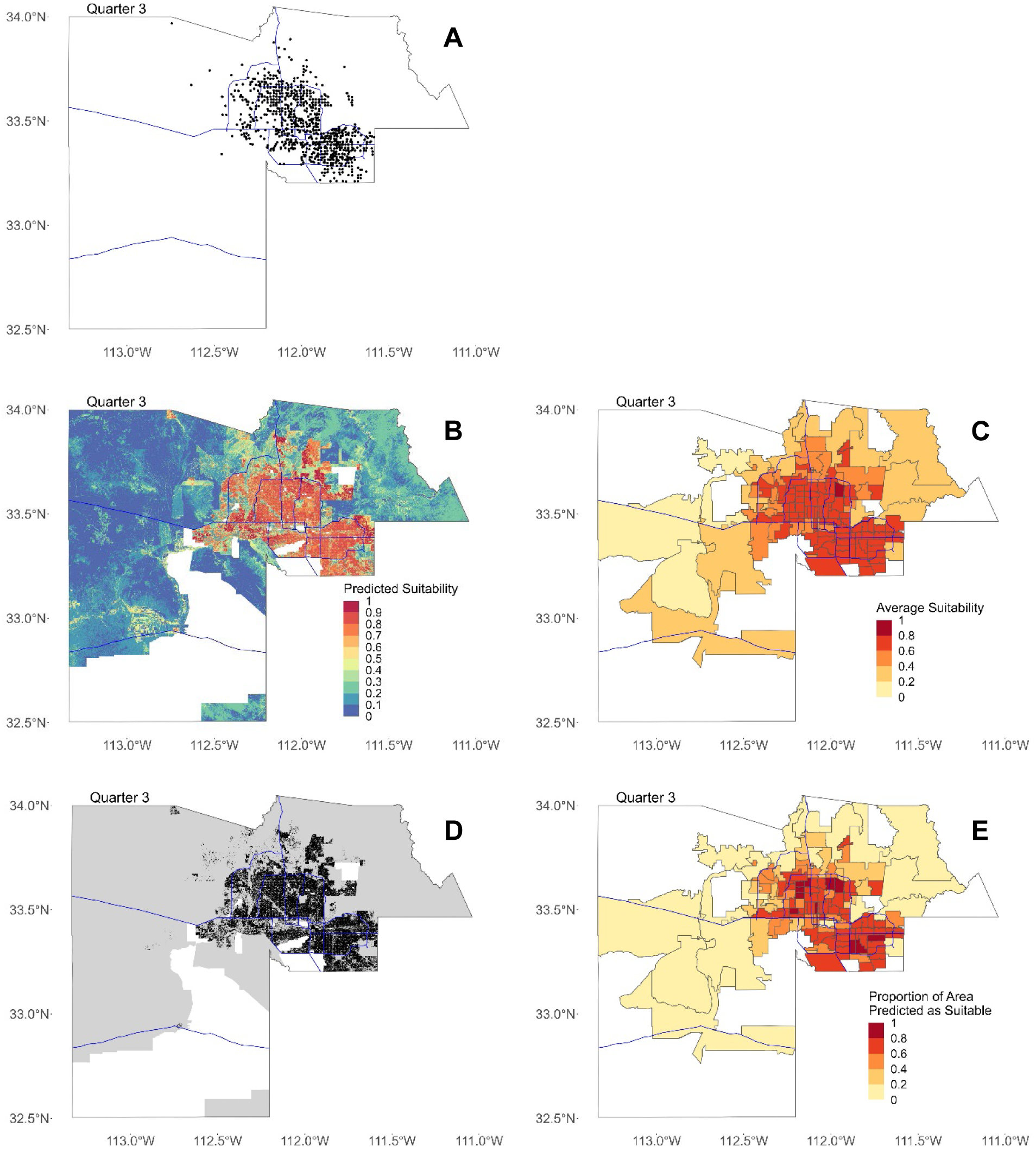
Maps for Quarter 3 (8/1/14—10/31/14). In all figures, blue lines represent major freeways in Maricopa County as a spatial reference. (**A**) Map of all locations where Ae. aegypti females were trapped in Maricopa County in Quarter 3. (**B**) Maxent output showing predicted suitability for 30-meter by 30-meter pixels. (**C**) Predicted suitability averaged by ZCTA. (**D**) Binary map showing areas predicted as suitable in black and not suitable in gray. In (B) and (C), areas with missing data where suitability was not predicted are shown in white. (**E**) Proportion of area in each ZCTA predicted as suitable after converting to binary map.

**Figure S4.**
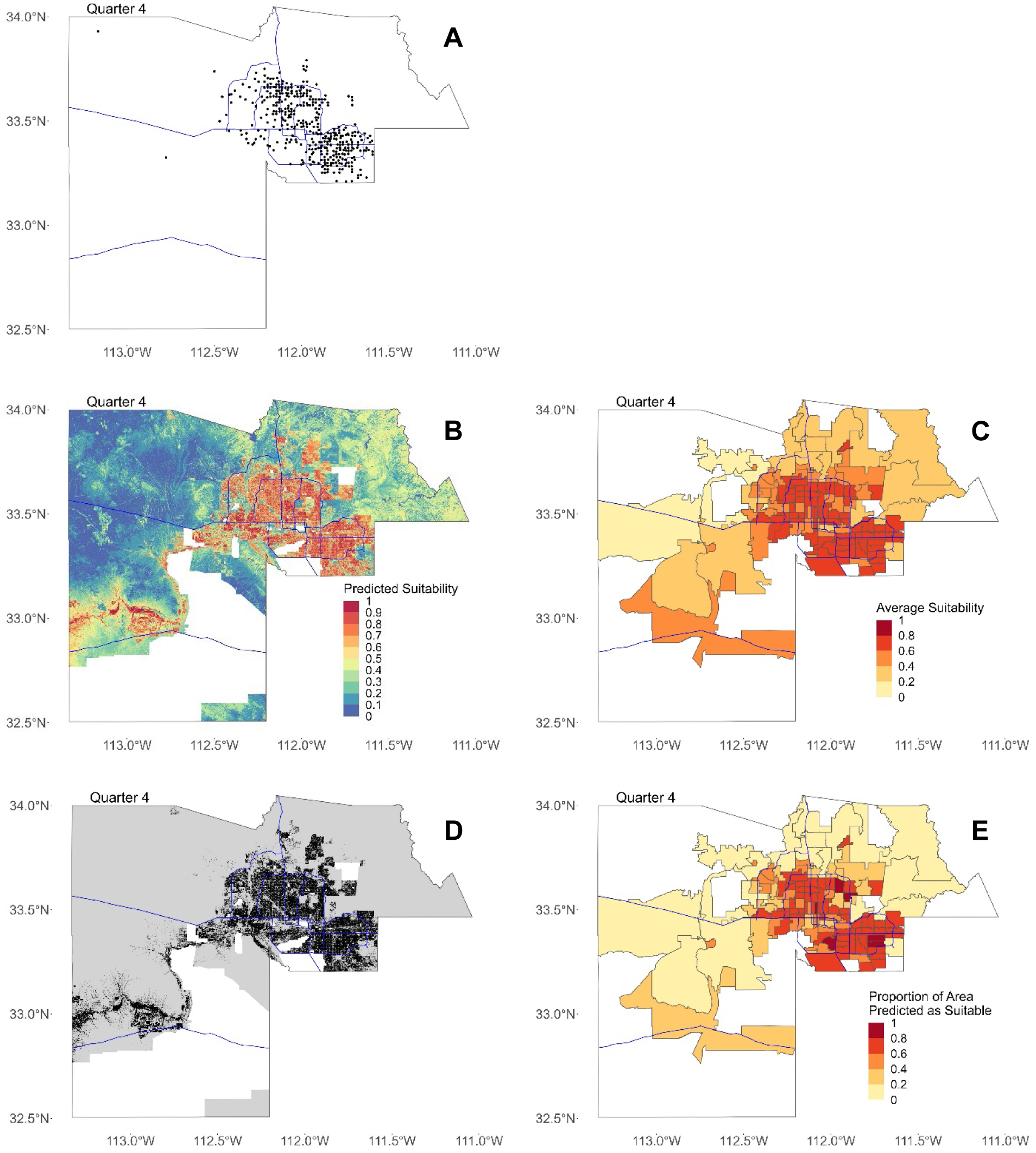
Maps for Quarter 4 (11/1/14—1/31/15). In all figures, blue lines represent major freeways in Maricopa County as a spatial reference. (**A**) Map of all locations where Ae. aegypti females were trapped in Maricopa County in Quarter 4. (**B**) Maxent output showing predicted suitability for 30-meter by 30-meter pixels. (**C**) Predicted suitability averaged by ZCTA. (**D**) Binary map showing areas predicted as suitable in black and not suitable in gray. In (B) and (C), areas with missing data where suitability was not predicted are shown in white. (**E**) Proportion of area in each ZCTA predicted as suitable after converting to binary map.

**Figure S5.**
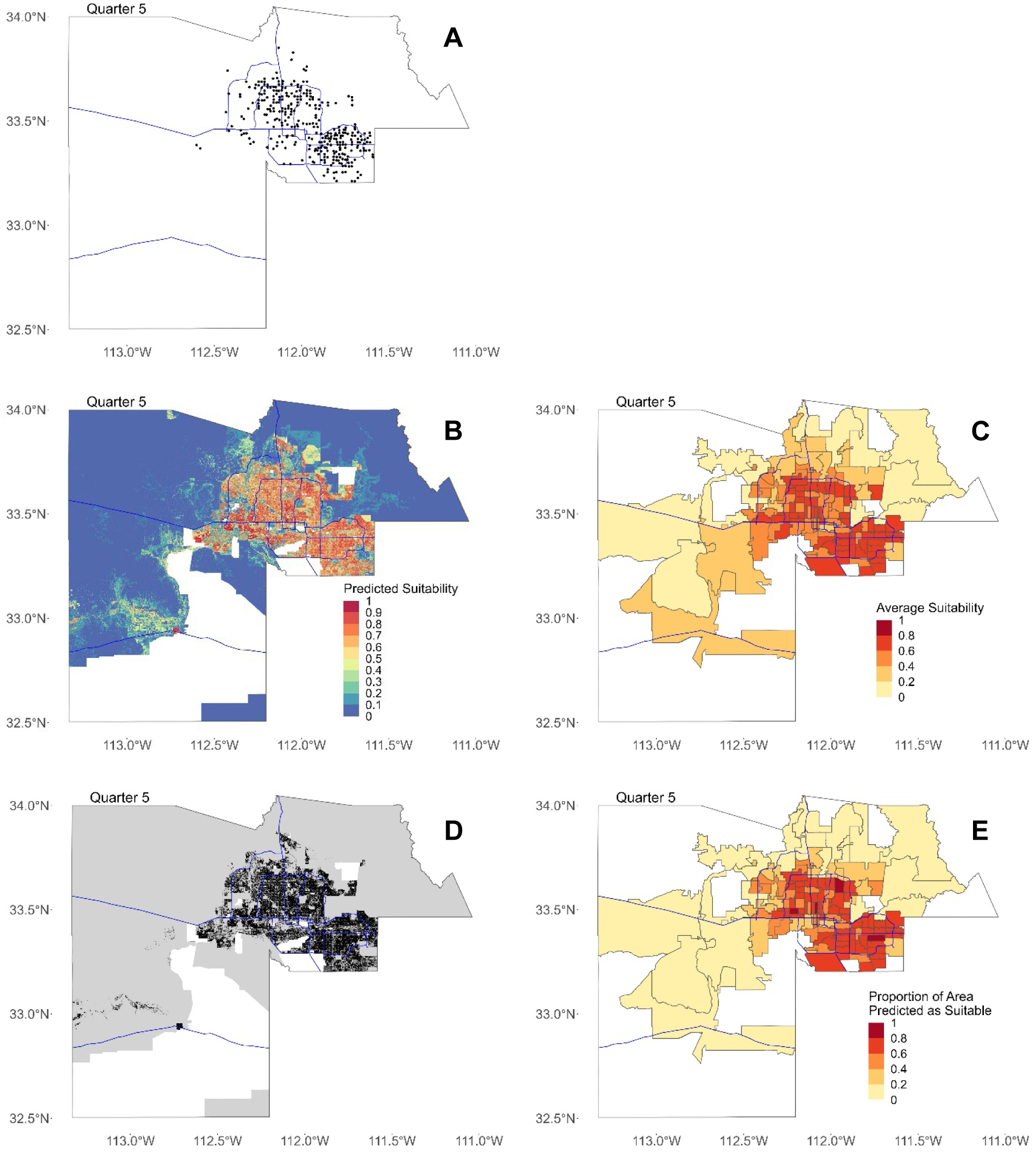
Maps for Quarter 5 (2/1/15—4/30/15). In all figures, blue lines represent major freeways in Maricopa County as a spatial reference. (**A**) Map of all locations where Ae. aegypti females were trapped in Maricopa County in Quarter 5. (**B**) Maxent output showing predicted suitability for 30-meter by 30-meter pixels. (**C**) Predicted suitability averaged by ZCTA. (**D**) Binary map showing areas predicted as suitable in black and not suitable in gray. In (B) and (C), areas with missing data where suitability was not predicted are shown in white. (**E**) Proportion of area in each ZCTA predicted as suitable after converting to binary map.

**Figure S6.**
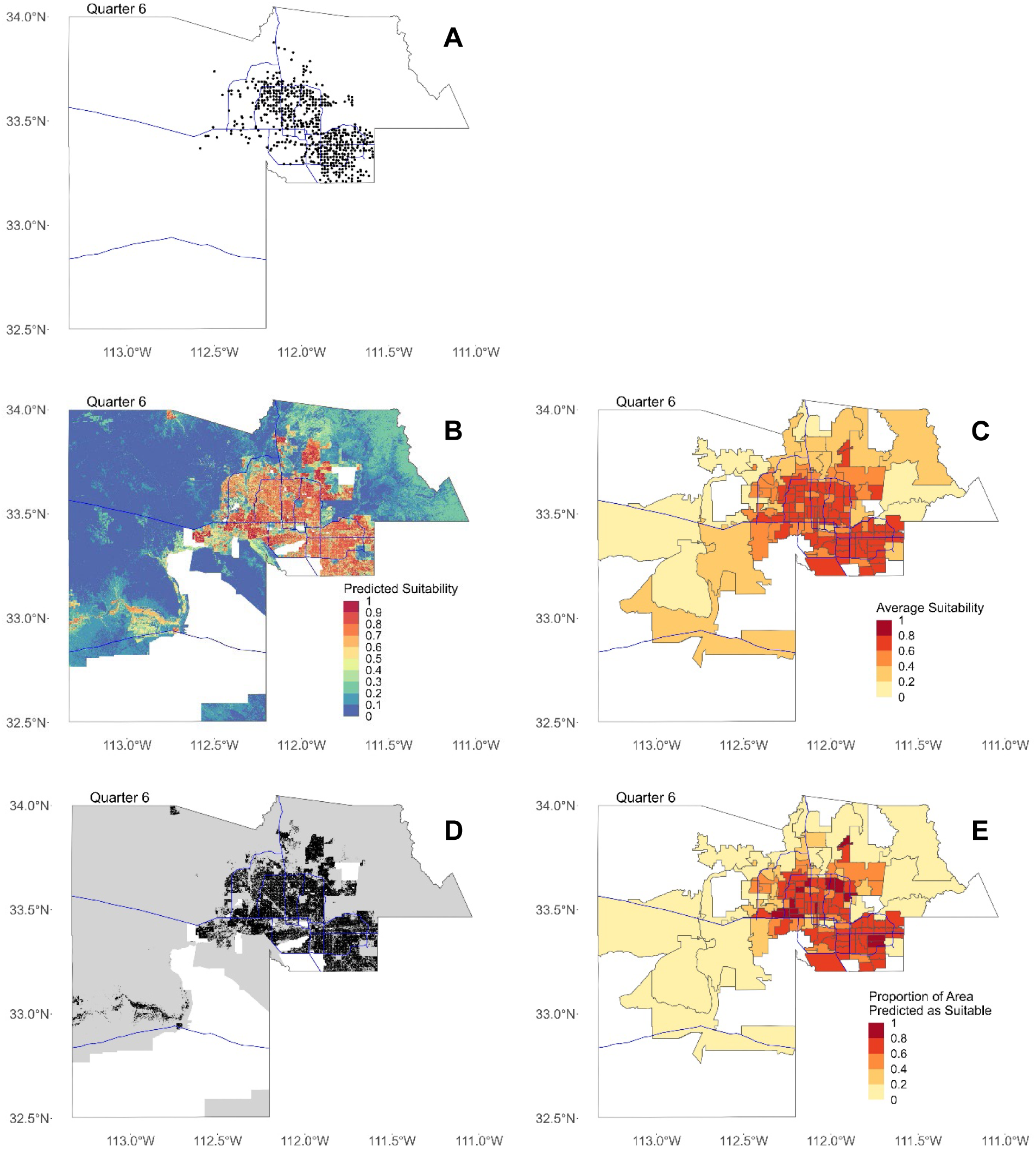
Maps for Quarter 6 (5/1/15—7/31/15). In all figures, blue lines represent major freeways in Maricopa County as a spatial reference. (**A**) Map of all locations where Ae. aegypti females were trapped in Maricopa County in Quarter 6. (**B**) Maxent output showing predicted suitability for 30-meter by 30-meter pixels. (**C**) Predicted suitability averaged by ZCTA. (**D**) Binary map showing areas predicted as suitable in black and not suitable in gray. In (B) and (C), areas with missing data where suitability was not predicted are shown in white. (**E**) Proportion of area in each ZCTA predicted as suitable after converting to binary map.

**Figure S7.**
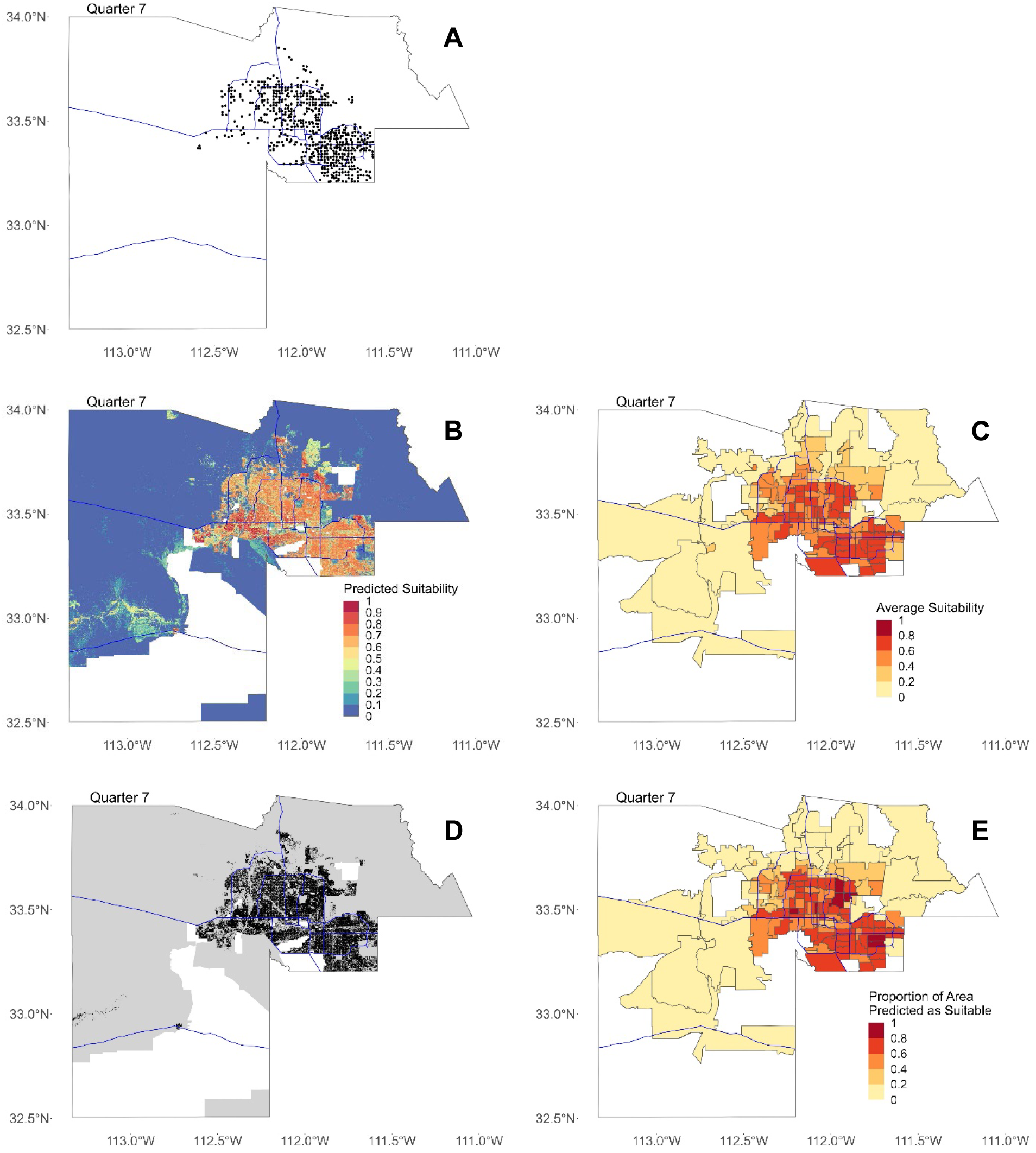
Maps for Quarter 7 (8/1/15—10/31/15). In all figures, blue lines represent major freeways in Maricopa County as a spatial reference. (**A**) Map of all locations where Ae. aegypti females were trapped in Maricopa County in Quarter 7. (**B**) Maxent output showing predicted suitability for 30-meter by 30-meter pixels. (**C**) Predicted suitability averaged by ZCTA. (**D**) Binary map showing areas predicted as suitable in black and not suitable in gray. In (B) and (C), areas with missing data where suitability was not predicted are shown in white. (**E**) Proportion of area in each ZCTA predicted as suitable after converting to binary map.

**Figure S8.**
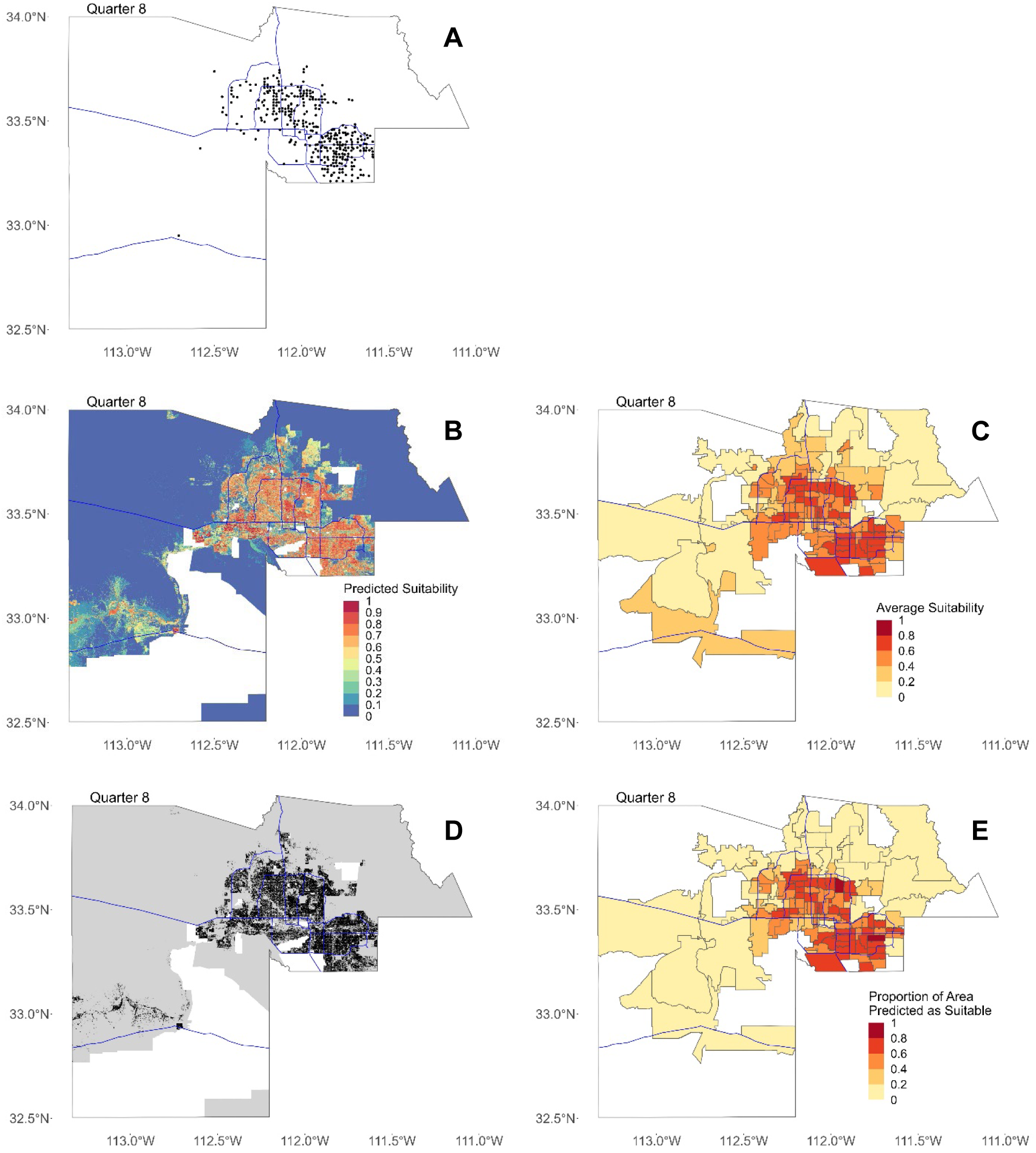
Maps for Quarter 8 (11/1/15—1/31/16). In all figures, blue lines represent major freeways in Maricopa County as a spatial reference. (**A**) Map of all locations where Ae. aegypti females were trapped in Maricopa County in Quarter 8. (**B**) Maxent output showing predicted suitability for 30-meter by 30-meter pixels. (**C**) Predicted suitability averaged by ZCTA. (**D**) Binary map showing areas predicted as suitable in black and not suitable in gray. In (B) and (C), areas with missing data where suitability was not predicted are shown in white. (**E**) Proportion of area in each ZCTA predicted as suitable after converting to binary map.

**Figure S9.**
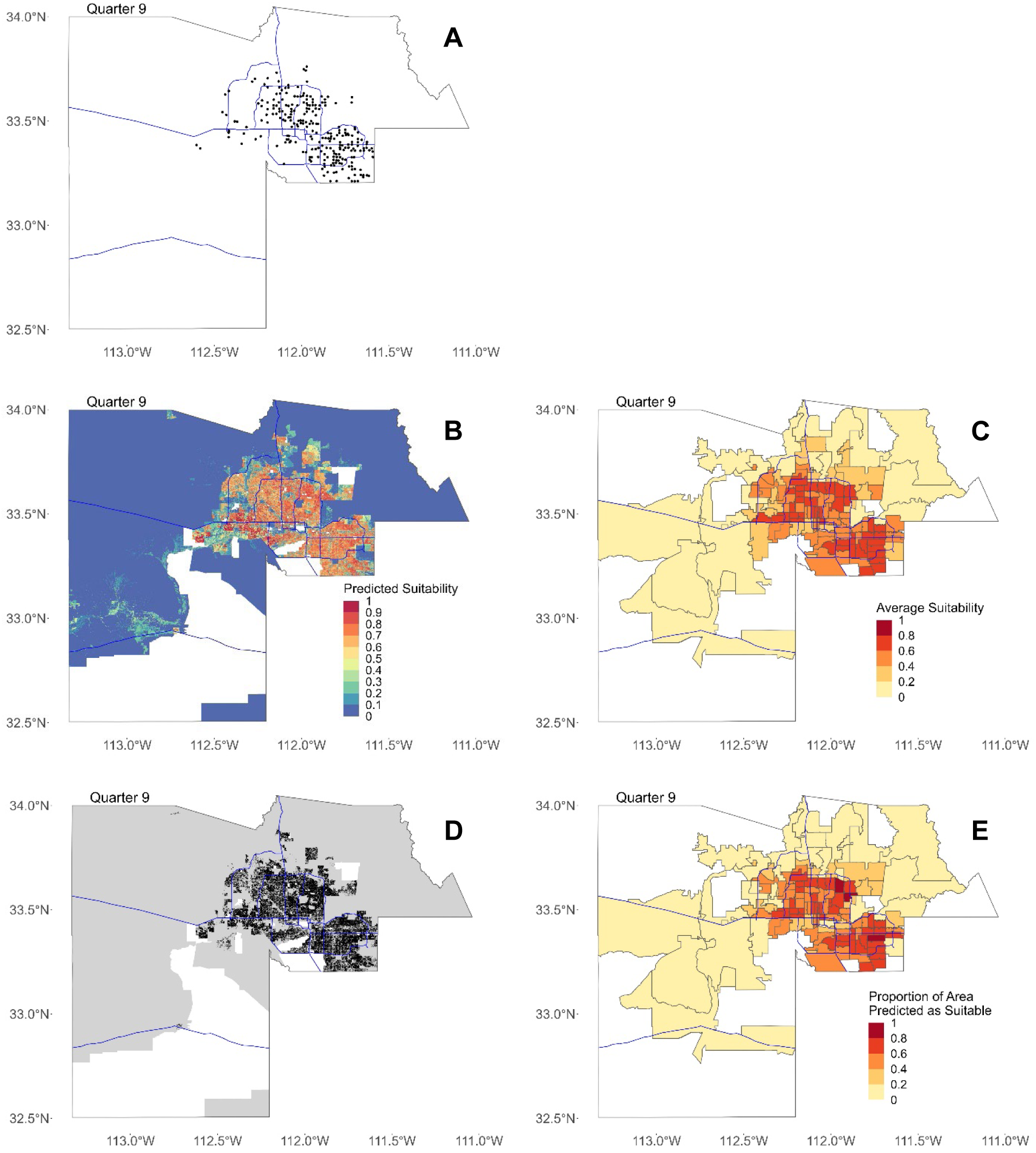
Maps for Quarter 9 (2/1/16—4/30/16). In all figures, blue lines represent major freeways in Maricopa County as a spatial reference. (**A**) Map of all locations where Ae. aegypti females were trapped in Maricopa County in Quarter 9. (**B**) Maxent output showing predicted suitability for 30-meter by 30-meter pixels. (**C**) Predicted suitability averaged by ZCTA. (**D**) Binary map showing areas predicted as suitable in black and not suitable in gray. In (B) and (C), areas with missing data where suitability was not predicted are shown in white. (**E**) Proportion of area in each ZCTA predicted as suitable after converting to binary map.

**Figure S10.**
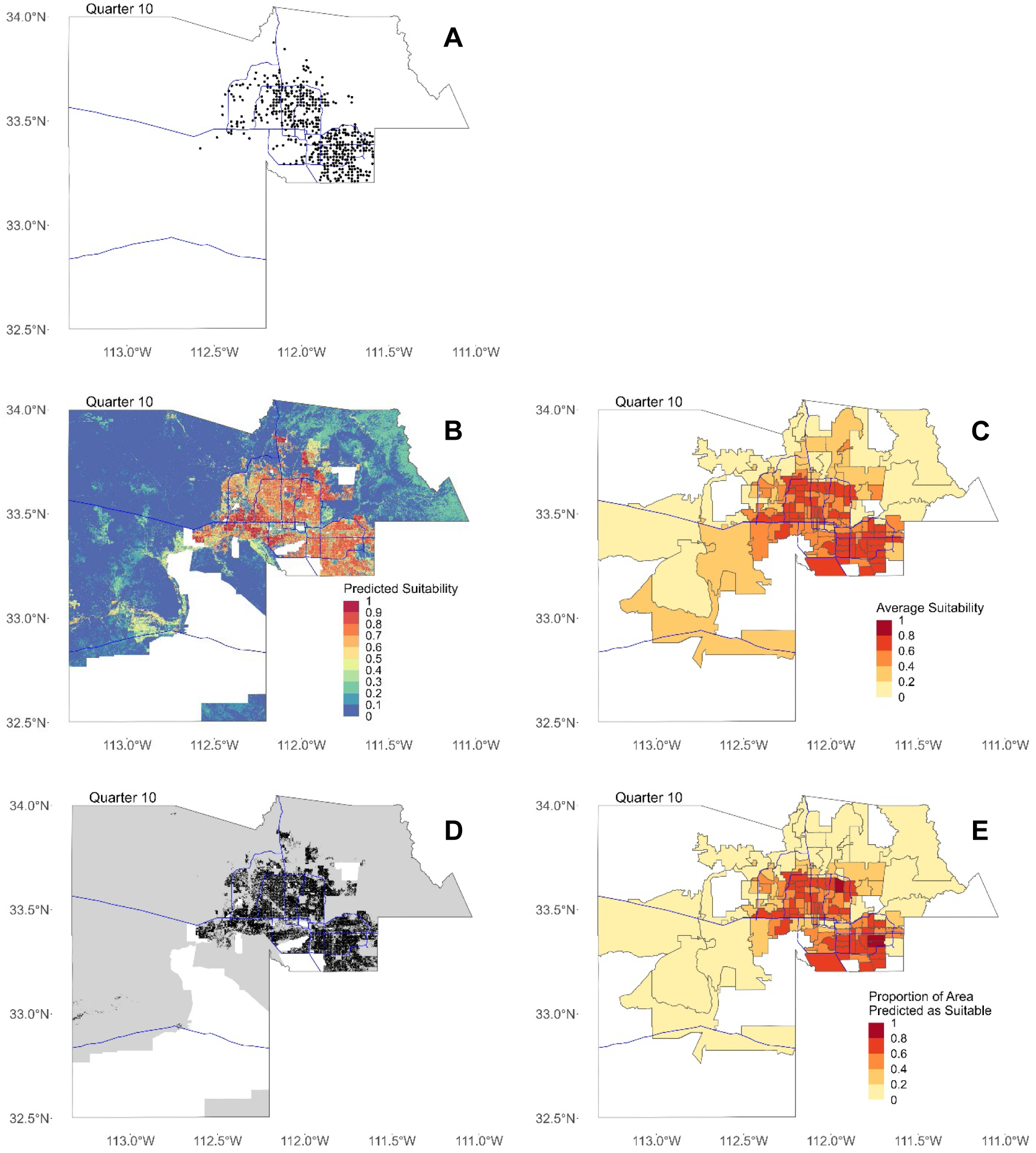
Maps for Quarter 10 (5/1/16—7/31/16). In all figures, blue lines represent major freeways in Maricopa County as a spatial reference. (**A**) Map of all locations where Ae. aegypti females were trapped in Maricopa County in Quarter 10. (**B**) Maxent output showing predicted suitability for 30-meter by 30-meter pixels. (**C**) Predicted suitability averaged by ZCTA. (**D**) Binary map showing areas predicted as suitable in black and not suitable in gray. In (B) and (C), areas with missing data where suitability was not predicted are shown in white. (**E**) Proportion of area in each ZCTA predicted as suitable after converting to binary map.

**Figure S11.**
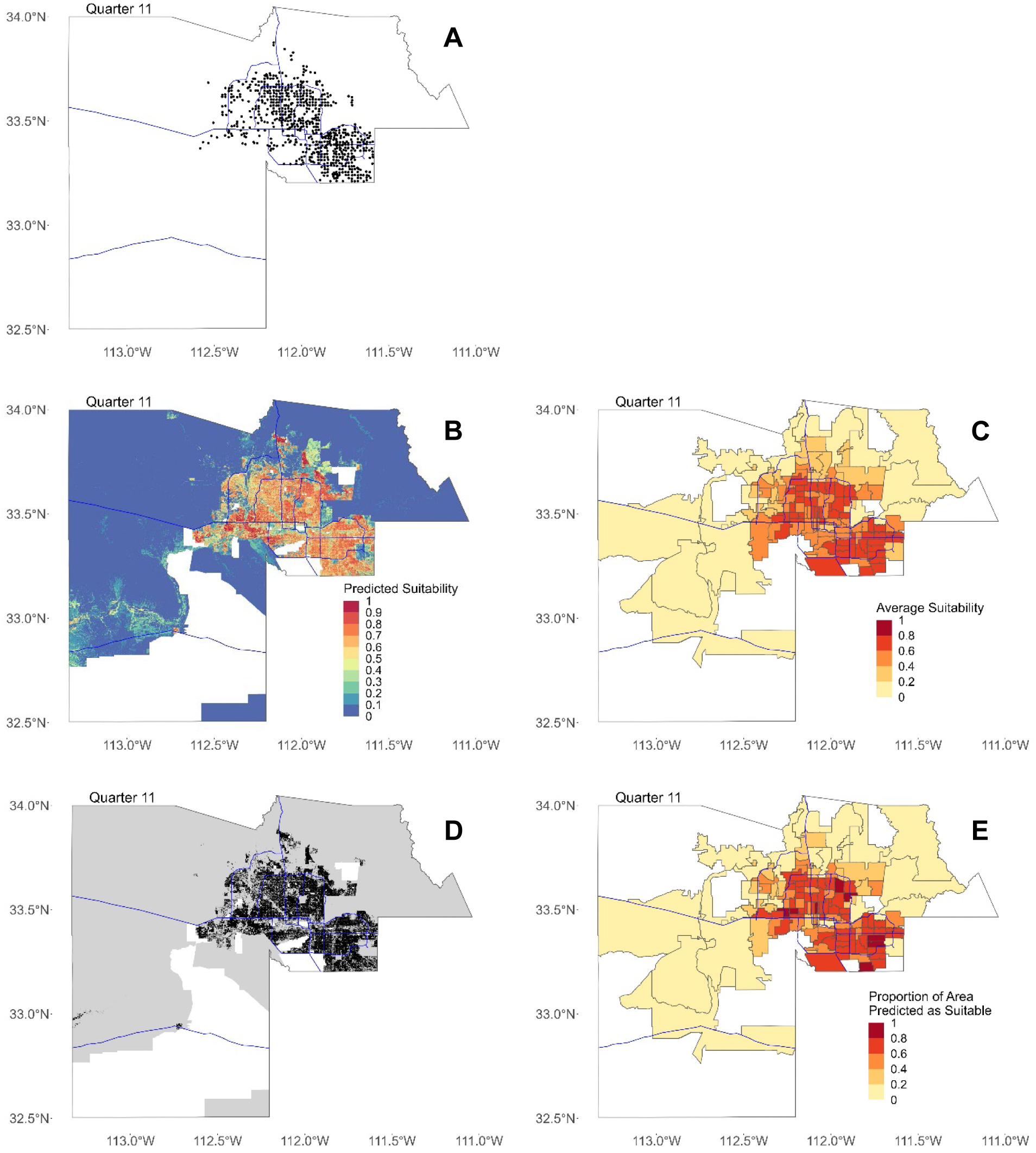
Maps for Quarter 11 (8/1/16—10/31/16). In all figures, blue lines represent major freeways in Maricopa County as a spatial reference. (**A**) Map of all locations where Ae. aegypti females were trapped in Maricopa County in Quarter 11. (**B**) Maxent output showing predicted suitability for 30-meter by 30-meter pixels. (**C**) Predicted suitability averaged by ZCTA. (**D**) Binary map showing areas predicted as suitable in black and not suitable in gray. In (B) and (C), areas with missing data where suitability was not predicted are shown in white. (**E**) Proportion of area in each ZCTA predicted as suitable after converting to binary map.

**Figure S12.**
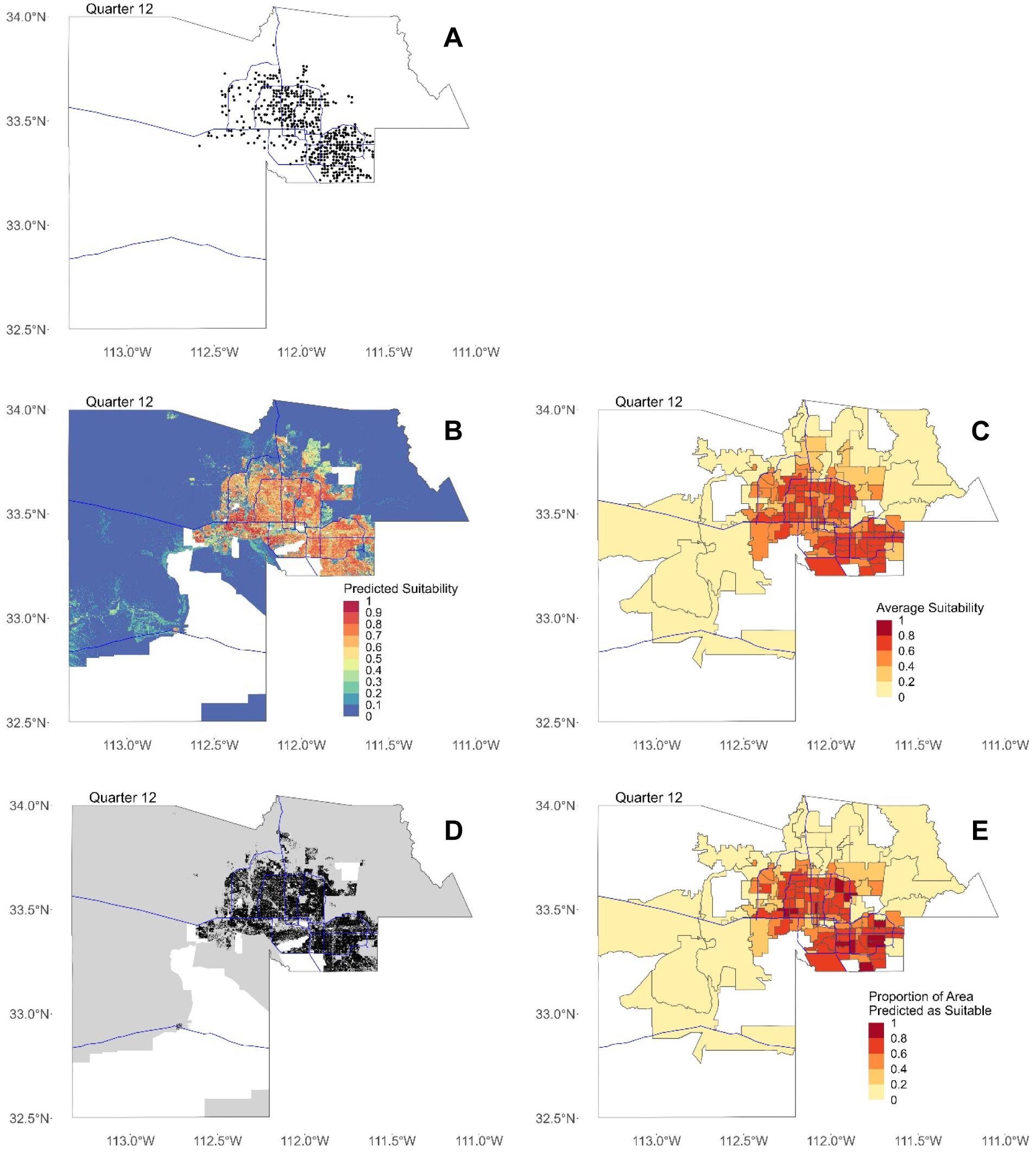
Maps for Quarter 12 (11/1/16—1/31/17). In all figures, blue lines represent major freeways in Maricopa County as a spatial reference. (**A**) Map of all locations where Ae. aegypti females were trapped in Maricopa County in Quarter 12. (**B**) Maxent output showing predicted suitability for 30-meter by 30-meter pixels. (**C**) Predicted suitability averaged by ZCTA. (**D**) Binary map showing areas predicted as suitable in black and not suitable in gray. In (B) and (C), areas with missing data where suitability was not predicted are shown in white. (**E**) Proportion of area in each ZCTA predicted as suitable after converting to binary map.

**Figure S13.**
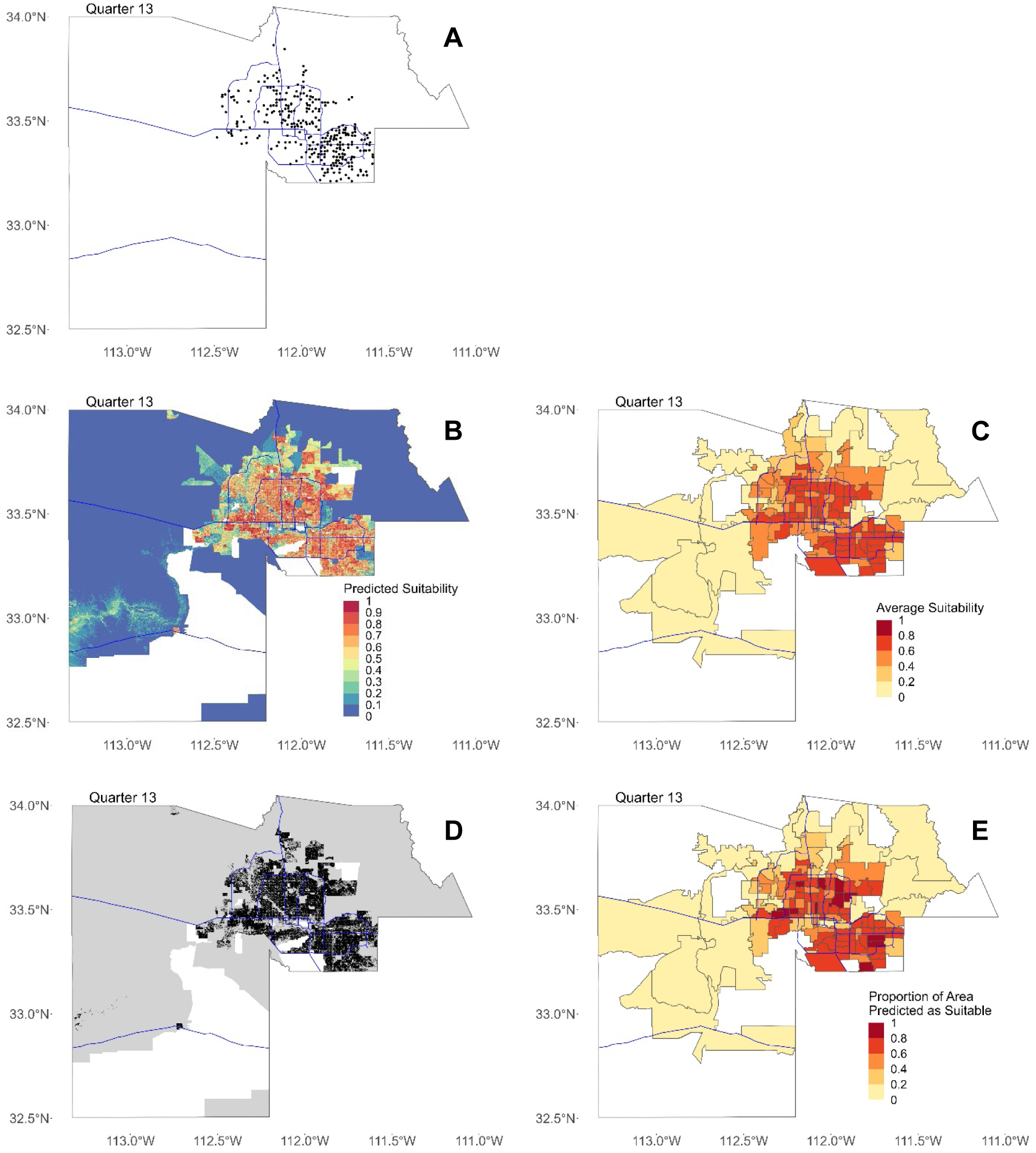
Maps for Quarter 13 (2/1/17—4/30/17). In all figures, blue lines represent major freeways in Maricopa County as a spatial reference. (**A**) Map of all locations where Ae. aegypti females were trapped in Maricopa County in Quarter 13. (**B**) Maxent output showing predicted suitability for 30-meter by 30-meter pixels. (**C**) Predicted suitability averaged by ZCTA. (**D**) Binary map showing areas predicted as suitable in black and not suitable in gray. In (B) and (C), areas with missing data where suitability was not predicted are shown in white. (**E**) Proportion of area in each ZCTA predicted as suitable after converting to binary map.

**Figure S14.**
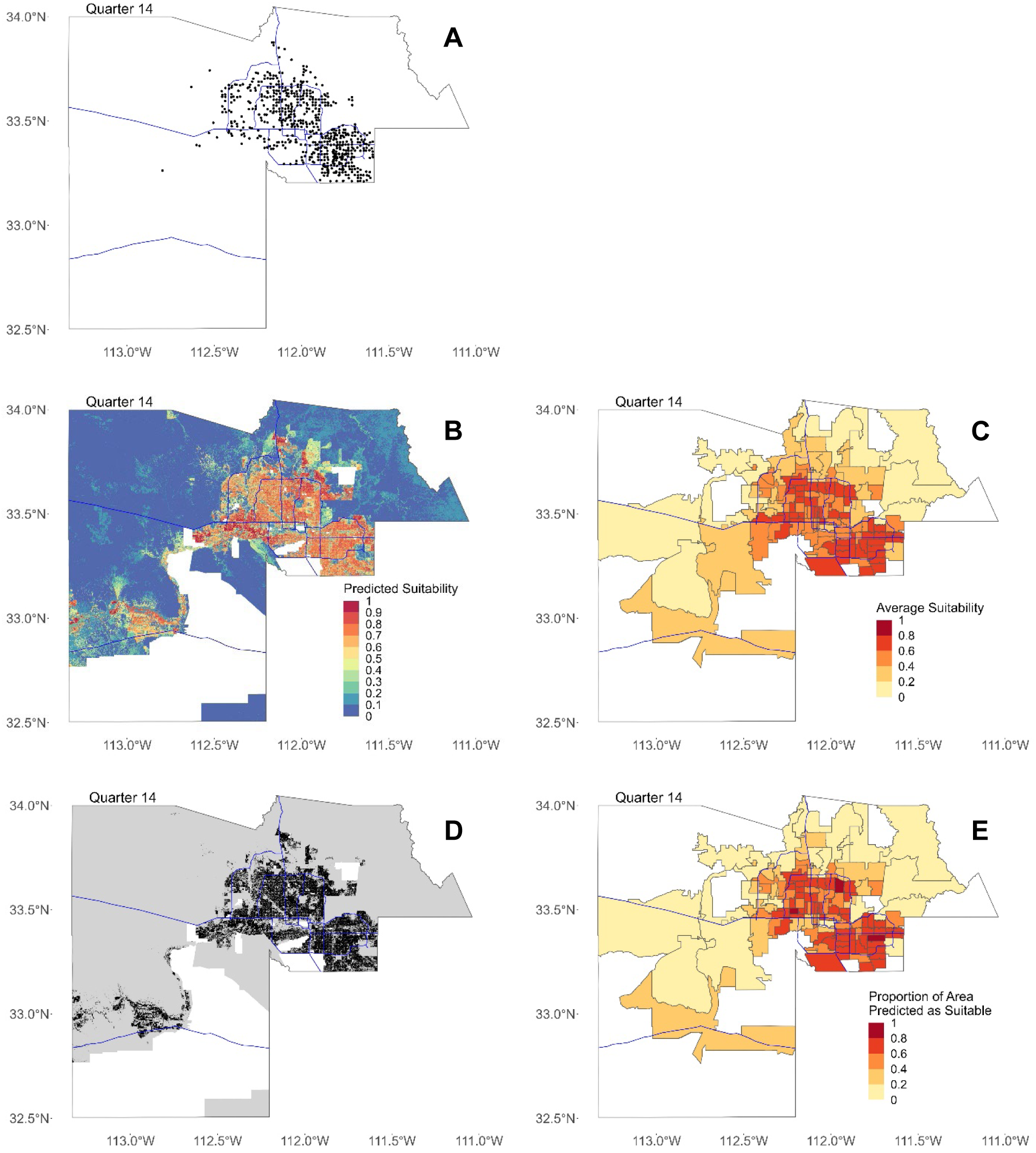
Maps for Quarter 14 (5/1/17—7/31/17). In all figures, blue lines represent major freeways in Maricopa County as a spatial reference. (**A**) Map of all locations where Ae. aegypti females were trapped in Maricopa County in Quarter 14. (**B**) Maxent output showing predicted suitability for 30-meter by 30-meter pixels. (**C**) Predicted suitability averaged by ZCTA. (**D**) Binary map showing areas predicted as suitable in black and not suitable in gray. In (B) and (C), areas with missing data where suitability was not predicted are shown in white. (**E**) Proportion of area in each ZCTA predicted as suitable after converting to binary map.

**Figure S15.**
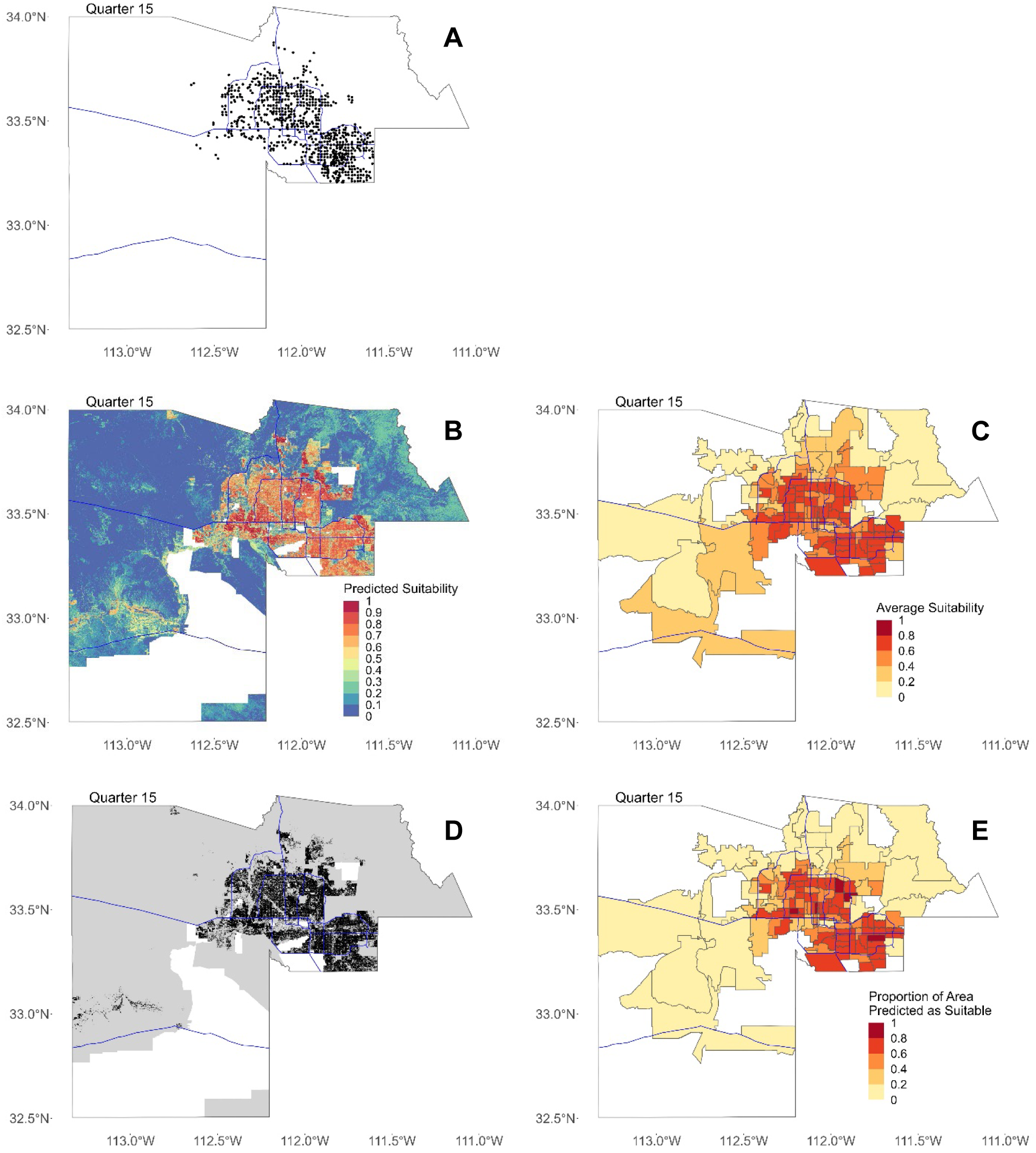
Maps for Quarter 15 (8/1/17—10/31/17). In all figures, blue lines represent major freeways in Maricopa County as a spatial reference. (**A**) Map of all locations where Ae. aegypti females were trapped in Maricopa County in Quarter 15. (**B**) Maxent output showing predicted suitability for 30-meter by 30-meter pixels. (**C**) Predicted suitability averaged by ZCTA. (**D**) Binary map showing areas predicted as suitable in black and not suitable in gray. In (B) and (C), areas with missing data where suitability was not predicted are shown in white. (**E**) Proportion of area in each ZCTA predicted as suitable after converting to binary map.

**Figure S16.**
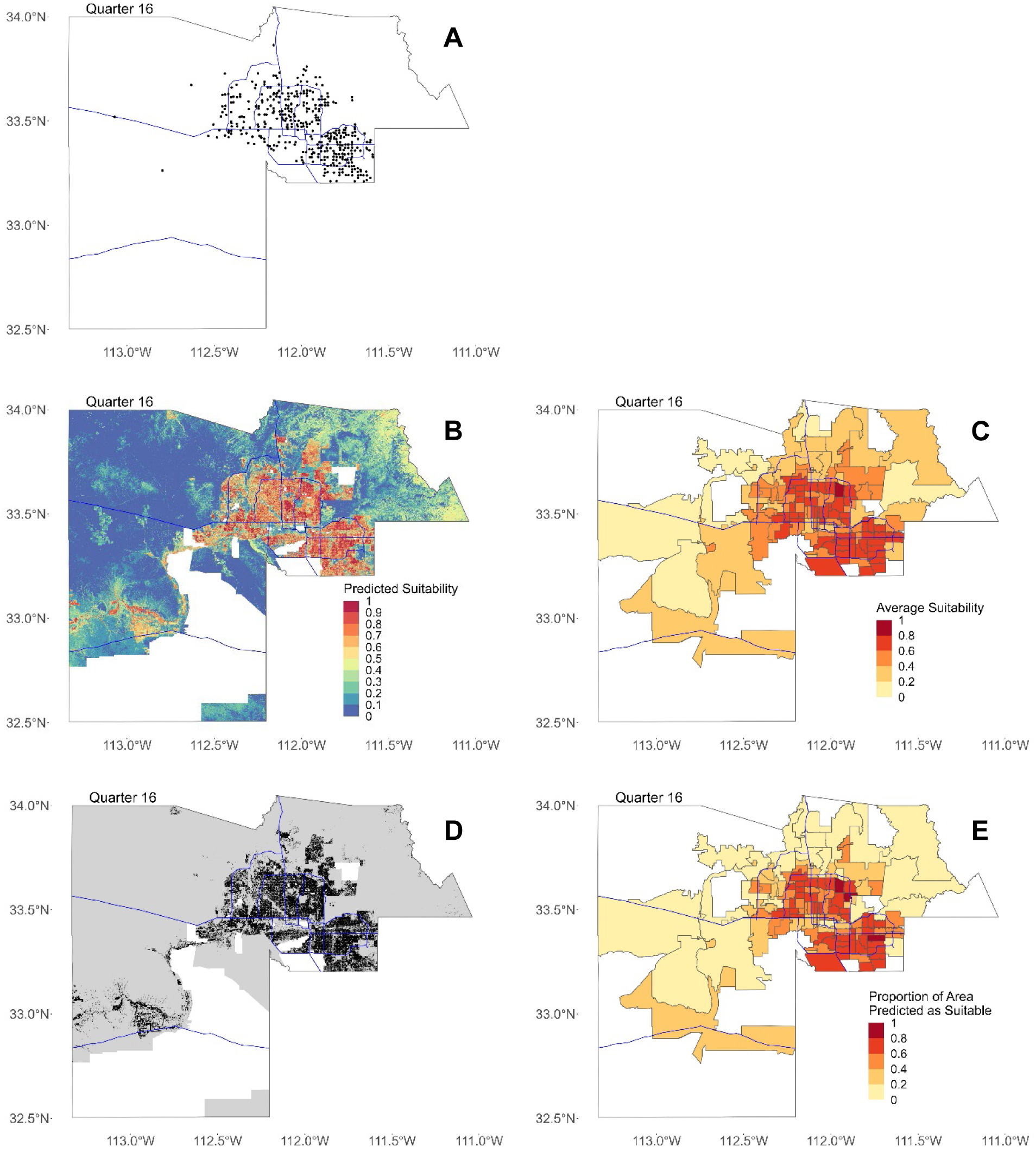
Maps for Quarter 16 (11/1/17—1/31/18). In all figures, blue lines represent major freeways in Maricopa County as a spatial reference. (**A**) Map of all locations where Ae. aegypti females were trapped in Maricopa County in Quarter 16. (**B**) Maxent output showing predicted suitability for 30-meter by 30-meter pixels. (**C**) Predicted suitability averaged by ZCTA. (**D**) Binary map showing areas predicted as suitable in black and not suitable in gray. In (B) and (C), areas with missing data where suitability was not predicted are shown in white. (**E**) Proportion of area in each ZCTA predicted as suitable after converting to binary map.

**Figure S17.**
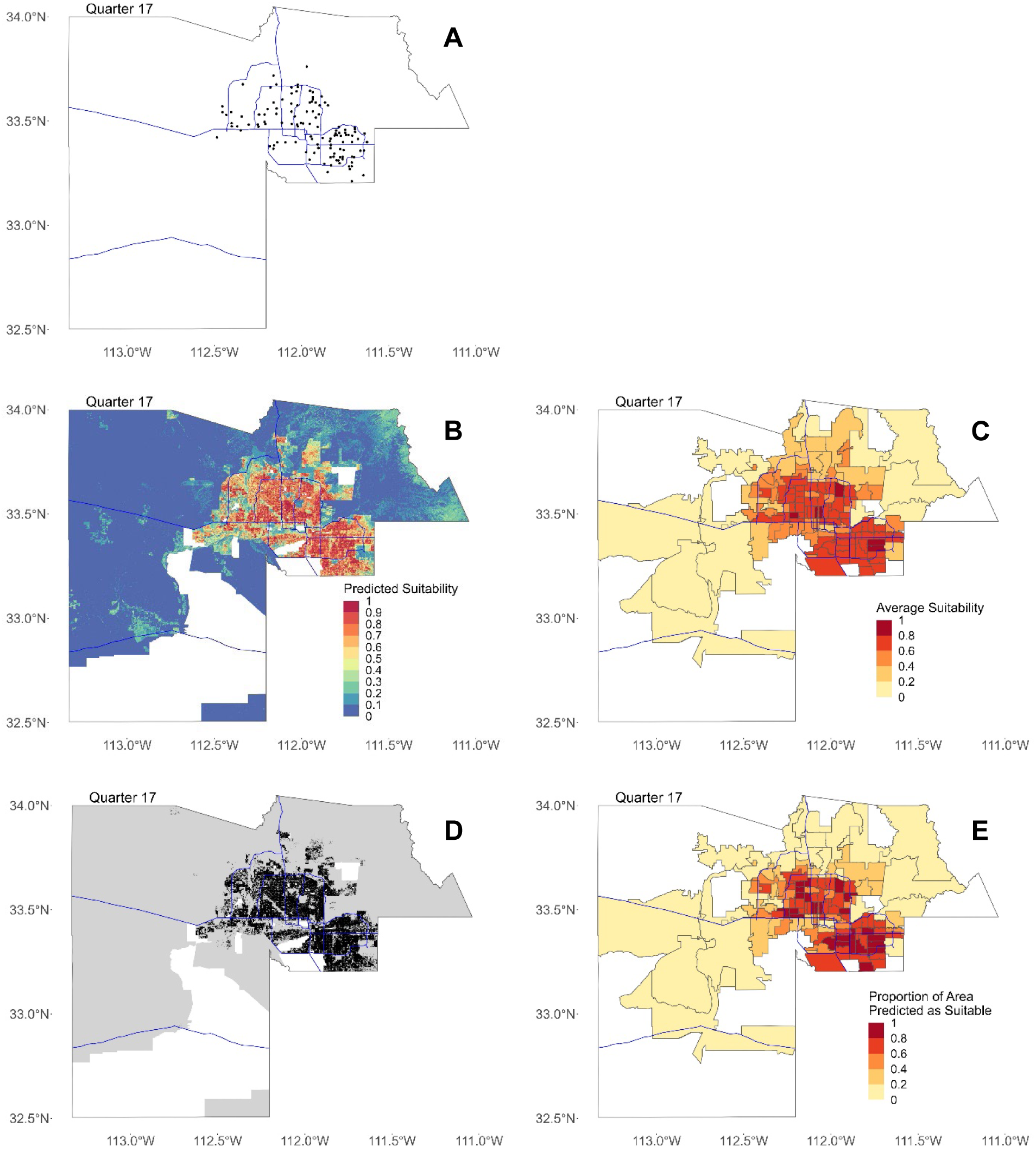
Maps for Quarter 17 (2/1/18—4/30/18). In all figures, blue lines represent major freeways in Maricopa County as a spatial reference. (**A**) Map of all locations where Ae. aegypti females were trapped in Maricopa County in Quarter 17. (**B**) Maxent output showing predicted suitability for 30-meter by 30-meter pixels. (**C**) Predicted suitability averaged by ZCTA. (**D**) Binary map showing areas predicted as suitable in black and not suitable in gray. In (B) and (C), areas with missing data where suitability was not predicted are shown in white. (**E**) Proportion of area in each ZCTA predicted as suitable after converting to binary map.

**Figure S18.**
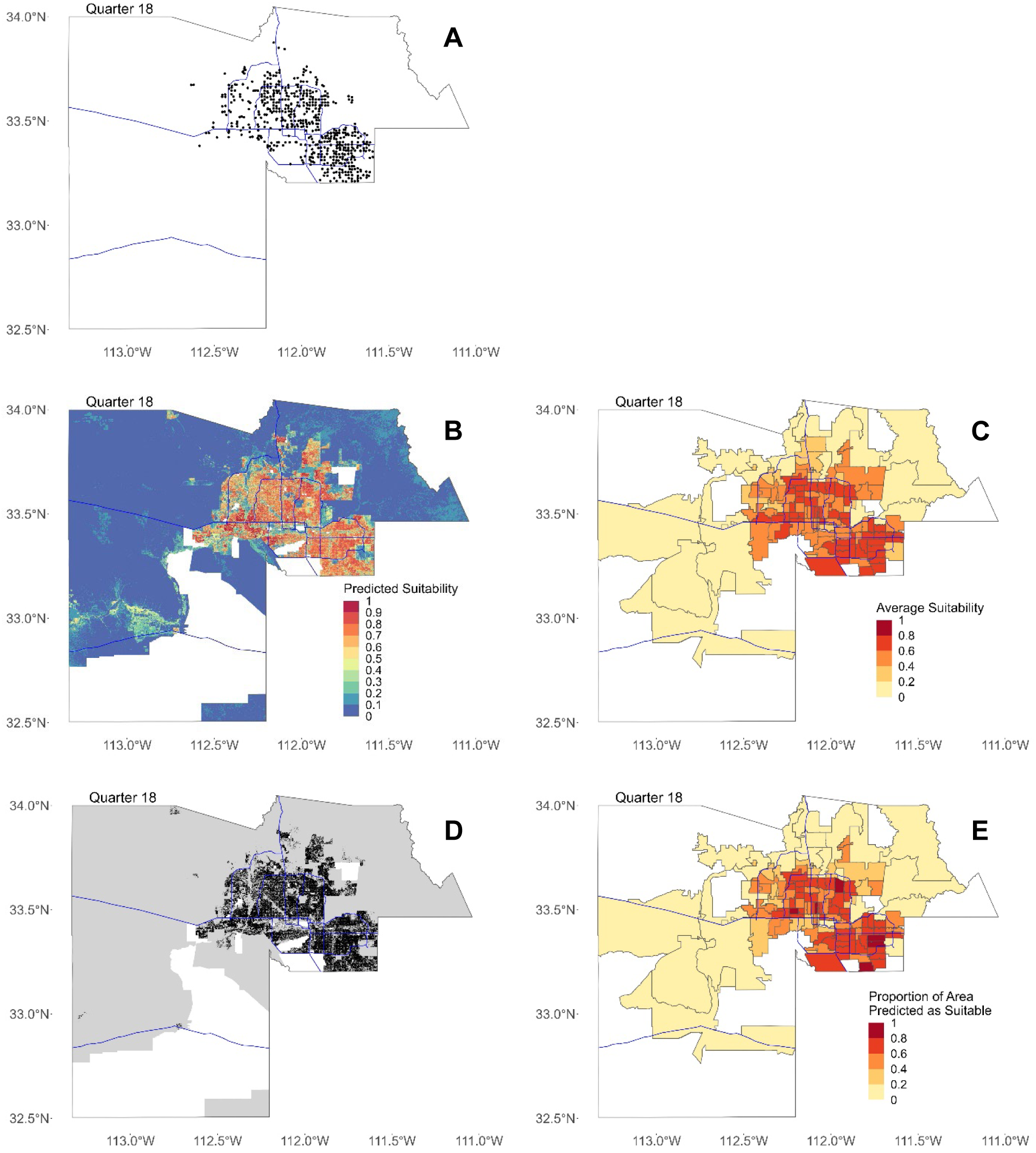
Maps for Quarter 18 (5/1/18—7/31/18). In all figures, blue lines represent major freeways in Maricopa County as a spatial reference. (**A**) Map of all locations where Ae. aegypti females were trapped in Maricopa County in Quarter 18. (**B**) Maxent output showing predicted suitability for 30-meter by 30-meter pixels. (**C**) Predicted suitability averaged by ZCTA. (**D**) Binary map showing areas predicted as suitable in black and not suitable in gray. In (B) and (C), areas with missing data where suitability was not predicted are shown in white. (**E**) Proportion of area in each ZCTA predicted as suitable after converting to binary map.

**Figure S19.**
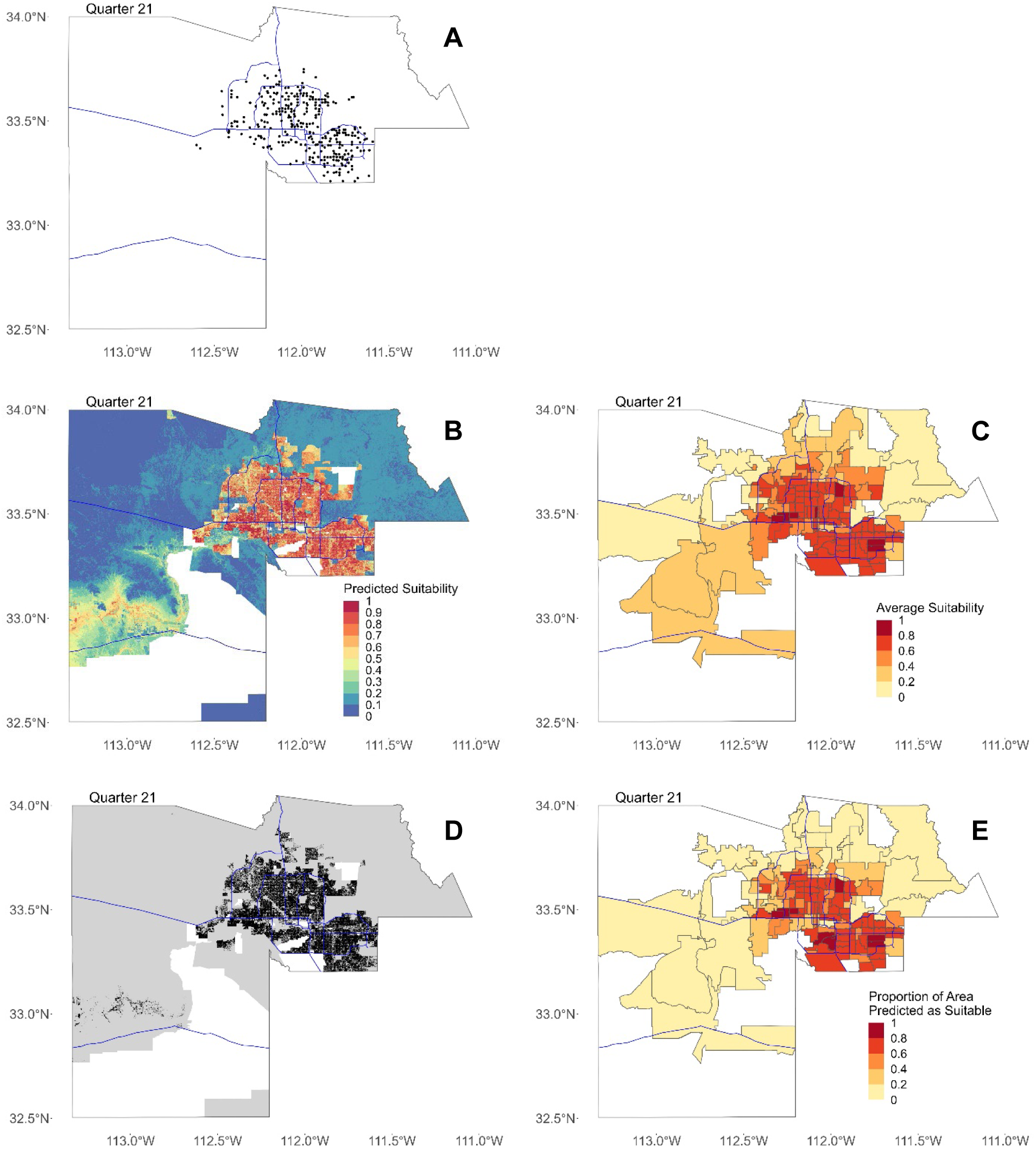
Maps for Quarter 21 (2/1/19—4/30/19). In all figures, blue lines represent major freeways in Maricopa County as a spatial reference. (**A**) Map of all locations where Ae. aegypti females were trapped in Maricopa County in Quarter 21. (**B**) Maxent output showing predicted suitability for 30-meter by 30-meter pixels. (**C**) Predicted suitability averaged by ZCTA. (**D**) Binary map showing areas predicted as suitable in black and not suitable in gray. In (B) and (C), areas with missing data where suitability was not predicted are shown in white. (**E**) Proportion of area in each ZCTA predicted as suitable after converting to binary map.

**Figure S20.**
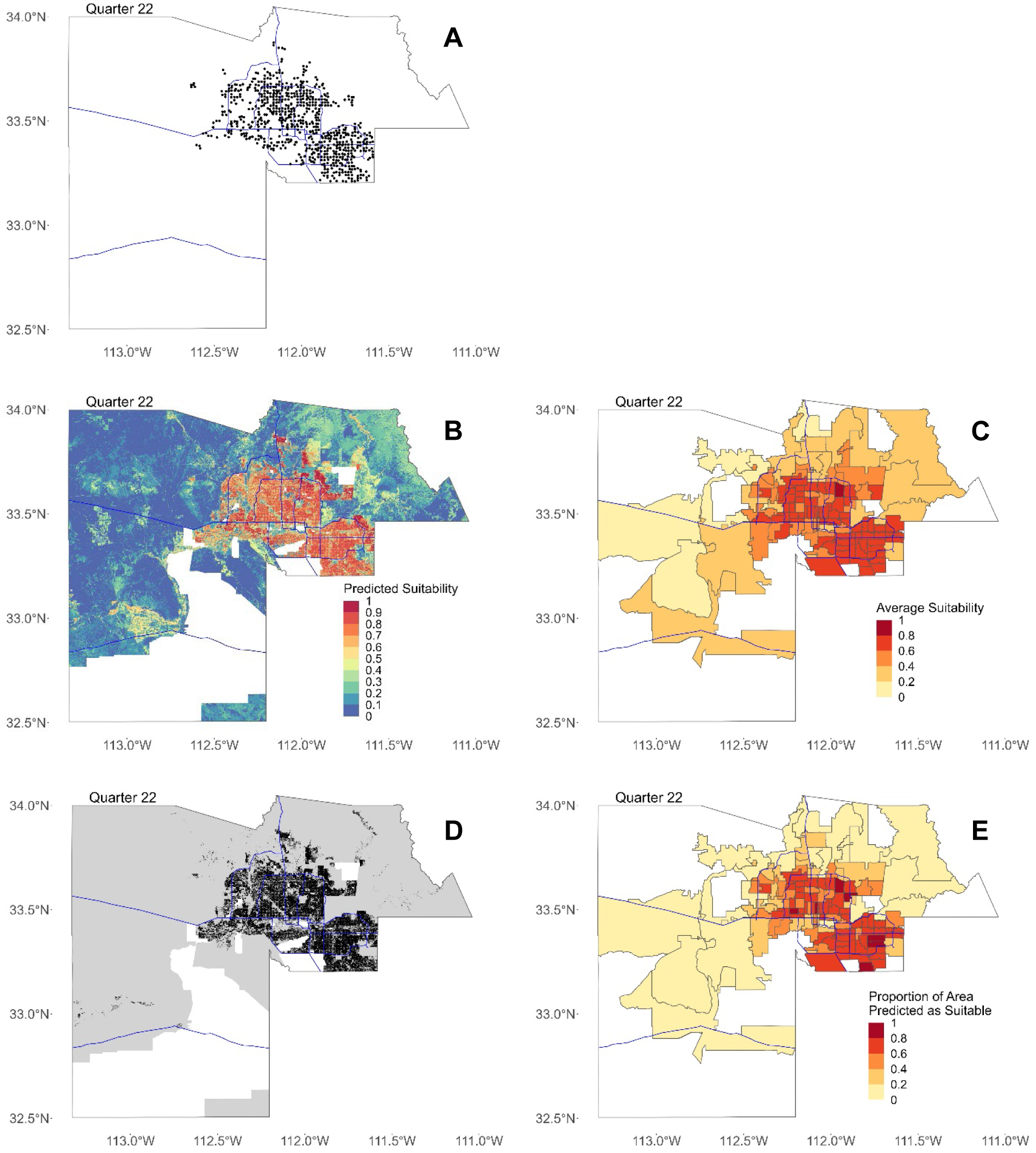
Maps for Quarter 22 (5/1/19—7/31/19). In all figures, blue lines represent major freeways in Maricopa County as a spatial reference. (**A**) Map of all locations where Ae. aegypti females were trapped in Maricopa County in Quarter 22. (**B**) Maxent output showing predicted suitability for 30-meter by 30-meter pixels. (**C**) Predicted suitability averaged by ZCTA. (**D**) Binary map showing areas predicted as suitable in black and not suitable in gray. In (B) and (C), areas with missing data where suitability was not predicted are shown in white. (**E**) Proportion of area in each ZCTA predicted as suitable after converting to binary map.

**Figure S21.**
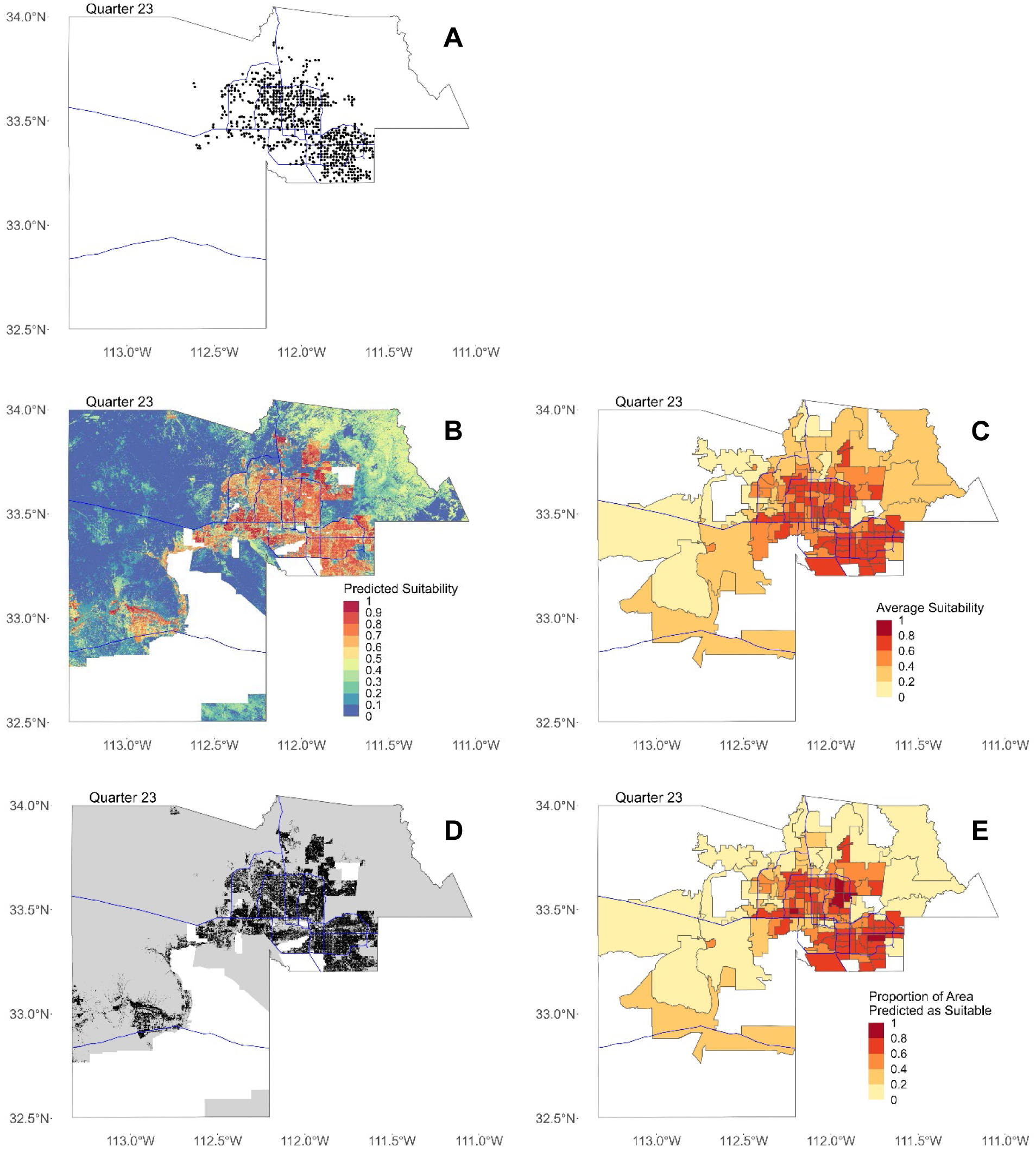
Maps for Quarter 23 (8/1/19—10/31/19). In all figures, blue lines represent major freeways in Maricopa County as a spatial reference. (**A**) Map of all locations where Ae. aegypti females were trapped in Maricopa County in Quarter 23. (**B**) Maxent output showing predicted suitability for 30-meter by 30-meter pixels. (**C**) Predicted suitability averaged by ZCTA. (**D**) Binary map showing areas predicted as suitable in black and not suitable in gray. In (B) and (C), areas with missing data where suitability was not predicted are shown in white. (**E**) Proportion of area in each ZCTA predicted as suitable after converting to binary map.

**Figure S22.**
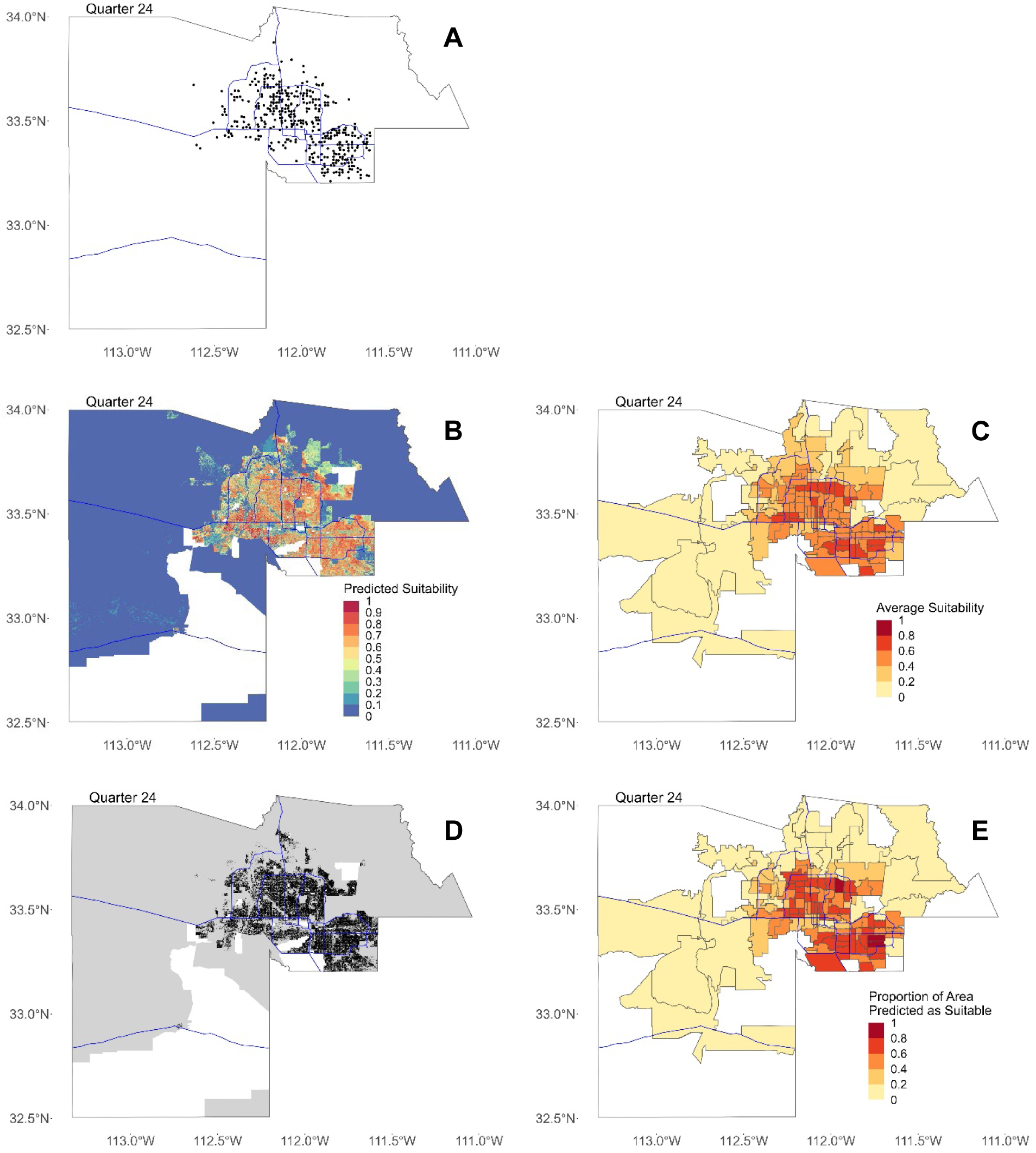
Maps for Quarter 24 (11/1/19—1/31/20). In all figures, blue lines represent major freeways in Maricopa County as a spatial reference. (**A**) Map of all locations where Ae. aegypti females were trapped in Maricopa County in Quarter 24. (**B**) Maxent output showing predicted suitability for 30-meter by 30-meter pixels. (**C**) Predicted suitability averaged by ZCTA. (**D**) Binary map showing areas predicted as suitable in black and not suitable in gray. In (B) and (C), areas with missing data where suitability was not predicted are shown in white. (**E**) Proportion of area in each ZCTA predicted as suitable after converting to binary map.

**Figure S23.**
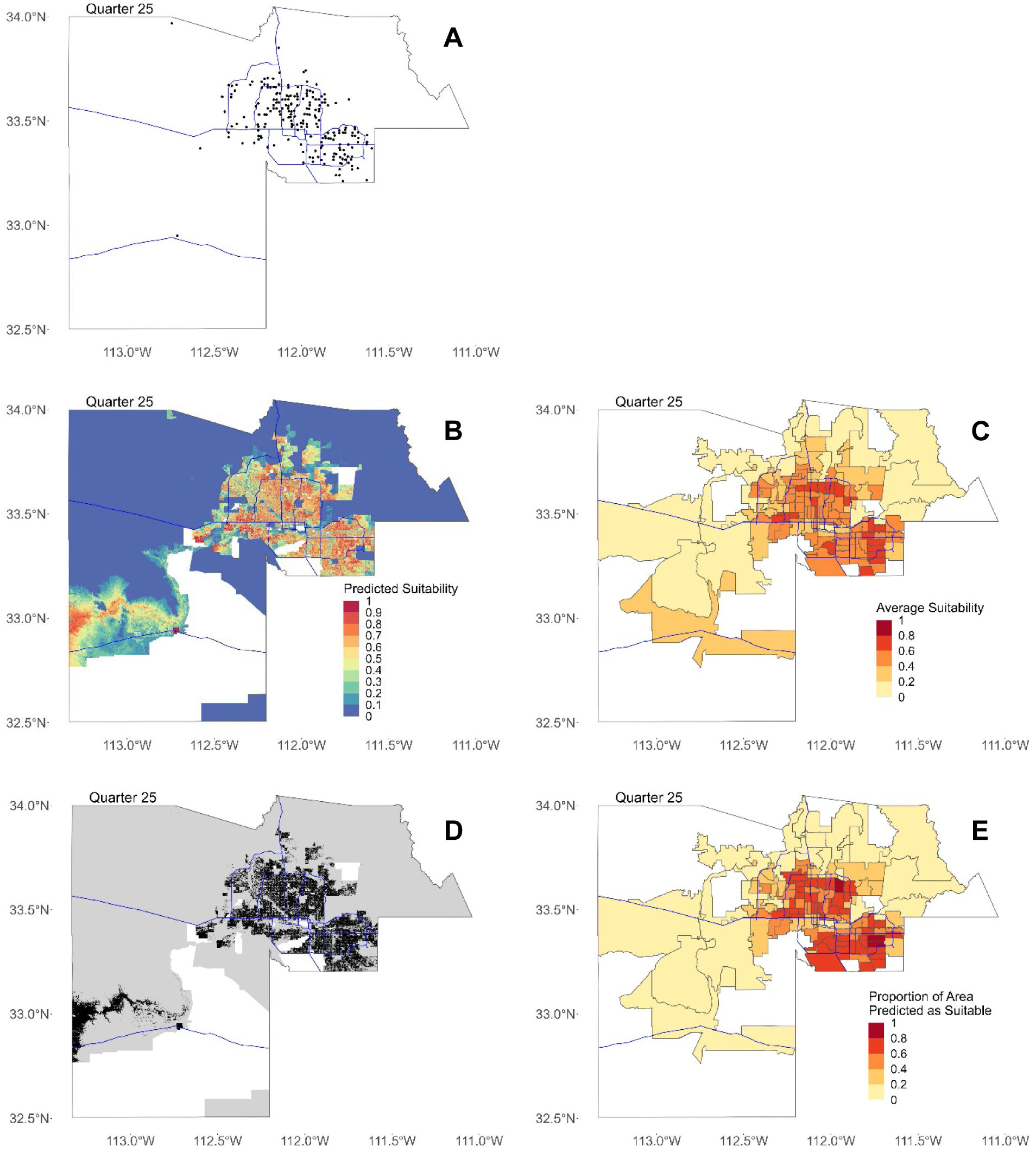
Maps for Quarter 25 (2/1/20—4/30/20). In all figures, blue lines represent major freeways in Maricopa County as a spatial reference. (**A**) Map of all locations where Ae. aegypti females were trapped in Maricopa County in Quarter 25. (**B**) Maxent output showing predicted suitability for 30-meter by 30-meter pixels. (**C**) Predicted suitability averaged by ZCTA. (**D**) Binary map showing areas predicted as suitable in black and not suitable in gray. In (B) and (C), areas with missing data where suitability was not predicted are shown in white. (**E**) Proportion of area in each ZCTA predicted as suitable after converting to binary map.

**Figure S24.**
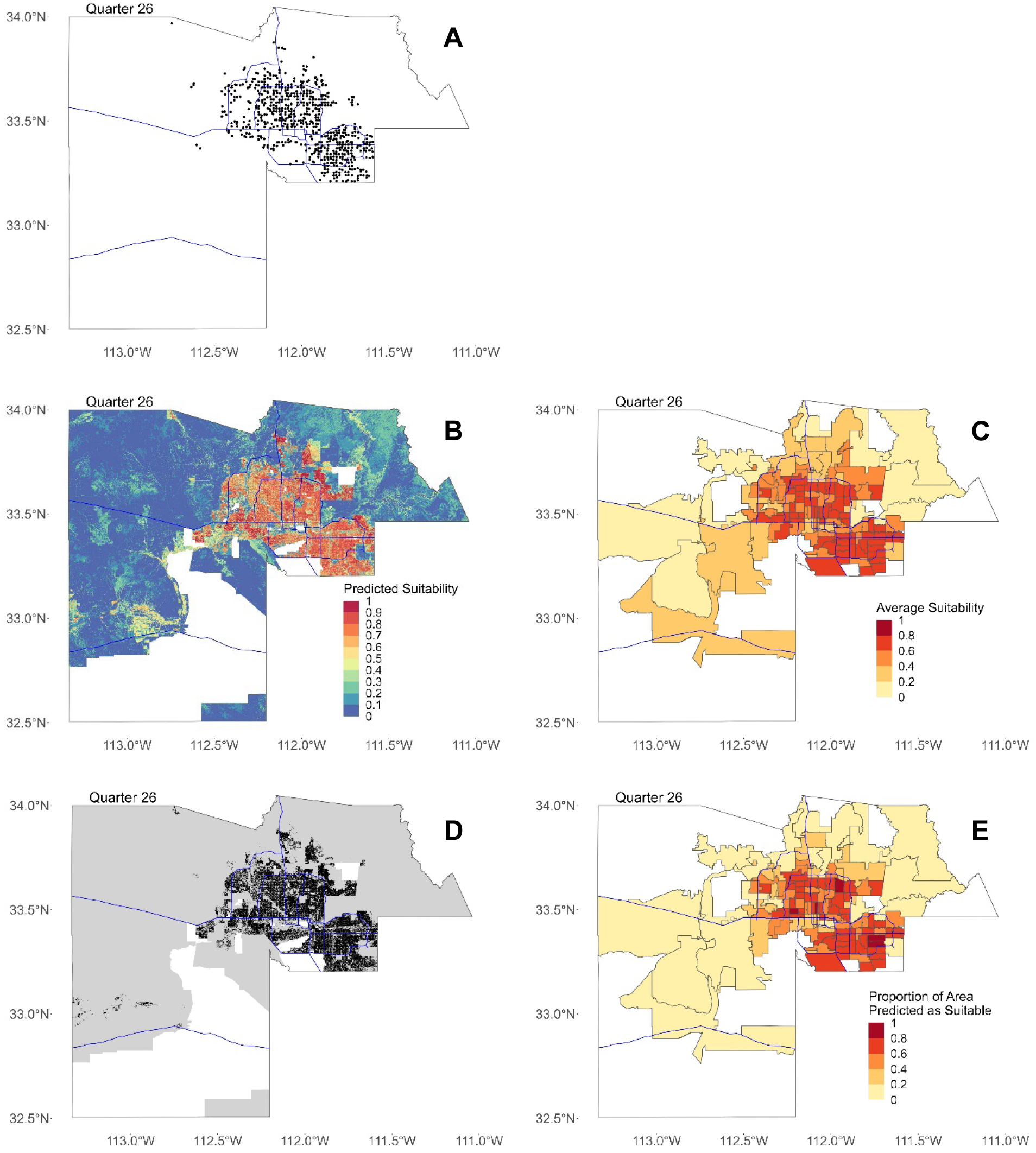
Maps for Quarter 26 (5/1/20—7/31/20). In all figures, blue lines represent major freeways in Maricopa County as a spatial reference. (**A**) Map of all locations where Ae. aegypti females were trapped in Maricopa County in Quarter 26. (**B**) Maxent output showing predicted suitability for 30-meter by 30-meter pixels. (**C**) Predicted suitability averaged by ZCTA. (**D**) Binary map showing areas predicted as suitable in black and not suitable in gray. In (B) and (C), areas with missing data where suitability was not predicted are shown in white. (**E**) Proportion of area in each ZCTA predicted as suitable after converting to binary map.

**Figure S25.**
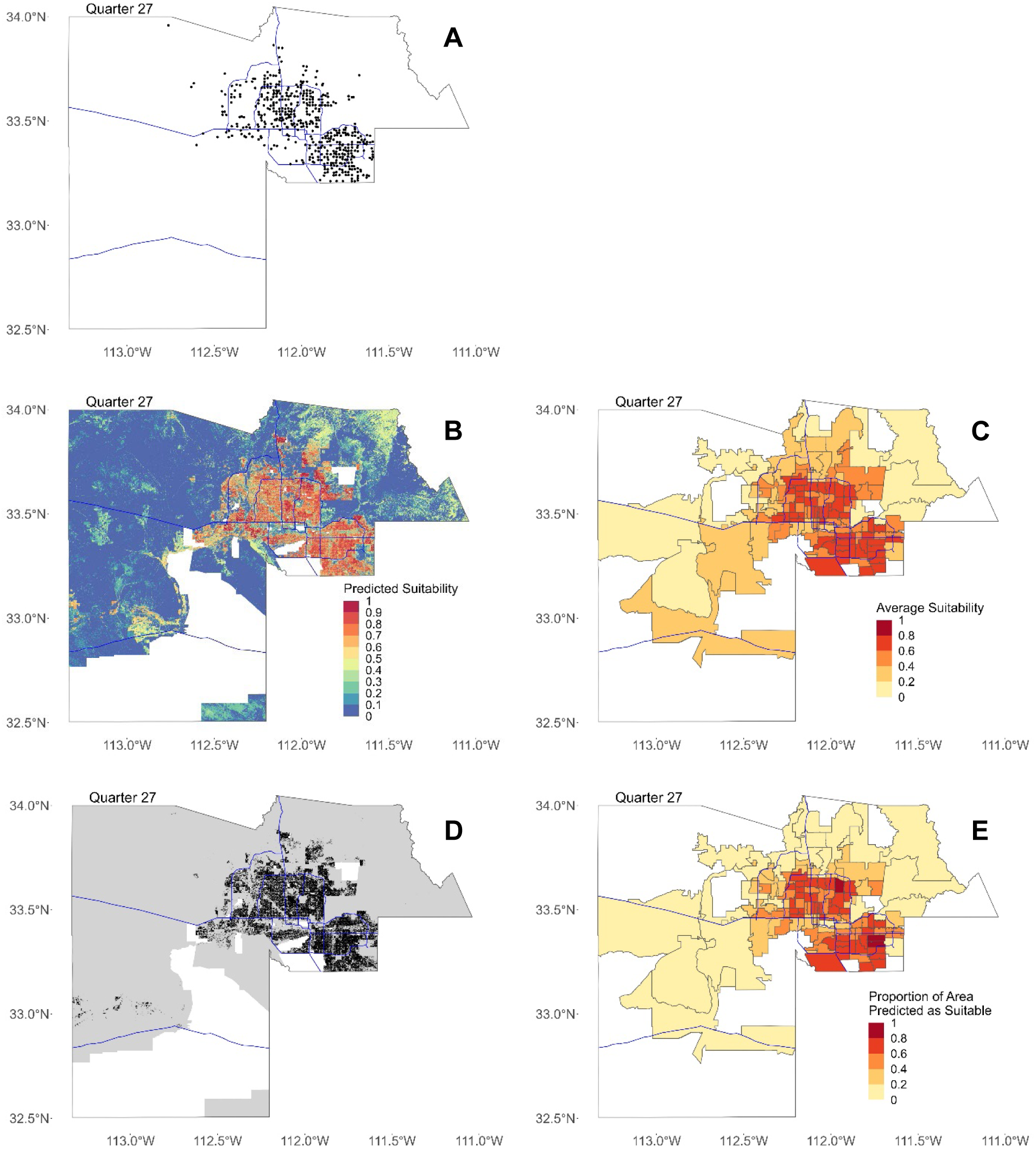
Maps for Quarter 27 (8/1/20—10/31/20). In all figures, blue lines represent major freeways in Maricopa County as a spatial reference. (**A**) Map of all locations where Ae. aegypti females were trapped in Maricopa County in Quarter 27. (**B**) Maxent output showing predicted suitability for 30-meter by 30-meter pixels. (**C**) Predicted suitability averaged by ZCTA. (**D**) Binary map showing areas predicted as suitable in black and not suitable in gray. In (B) and (C), areas with missing data where suitability was not predicted are shown in white. (**E**) Proportion of area in each ZCTA predicted as suitable after converting to binary map.

**Figure S26.**
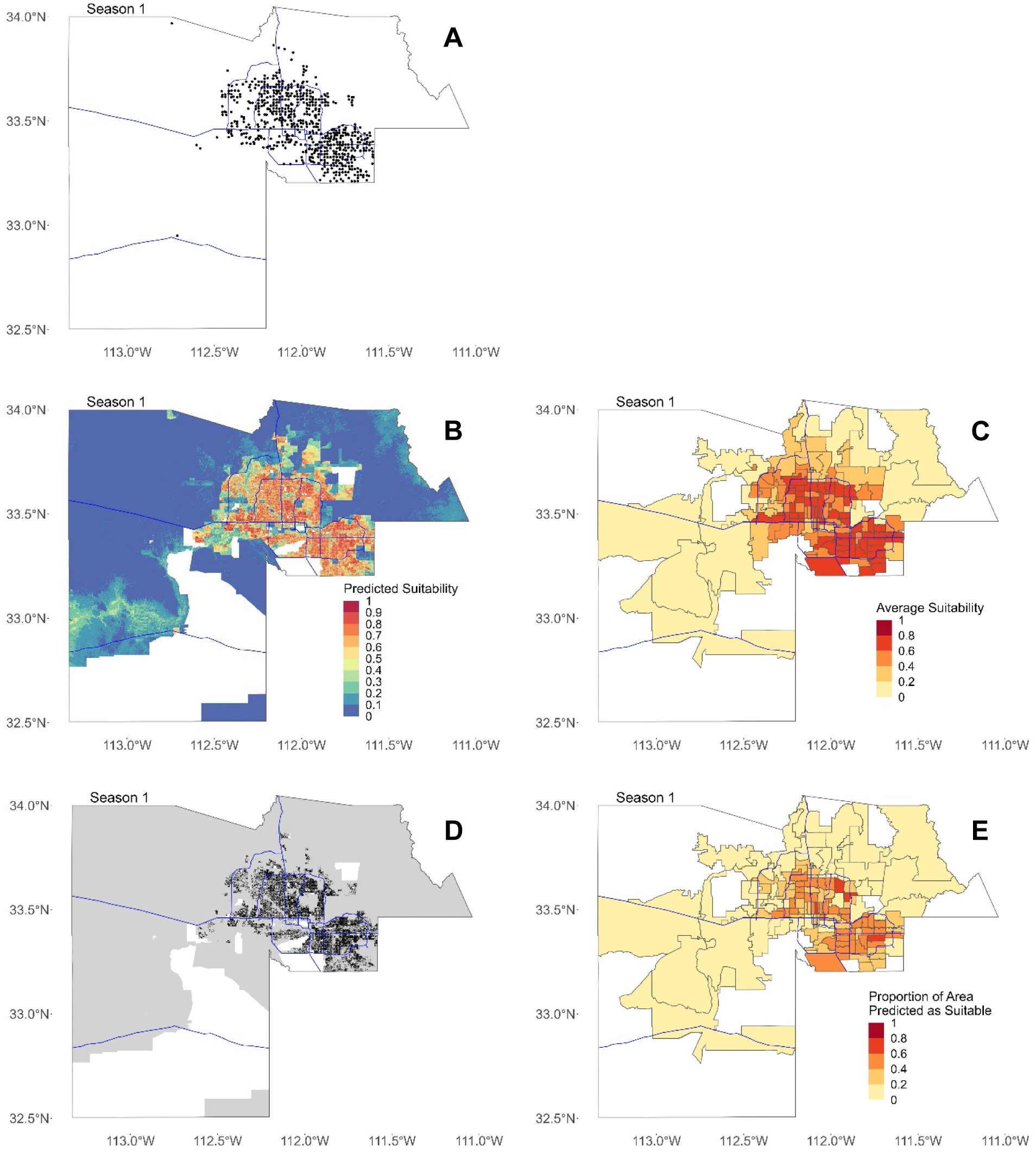
Maps for Season 1 (2/1-4/30 quarters). In all figures, blue lines represent major freeways in Maricopa County as a spatial reference. (**A**) Map of all locations where Ae. aegypti females were trapped in Maricopa County in Season 1. (**B**) Map showing mean predicted suitability for 30-meter by 30-meter pixels. (**C**) Mean predicted suitability averaged by ZCTA. (**D**) Binary map showing areas predicted as suitable for all quarters in black and not suitable in gray. In (B) and (C), areas with missing data where suitability was not predicted are shown in white. (**E**) Proportion of area in each ZCTA predicted as suitable after converting to binary map.

**Figure S27.**
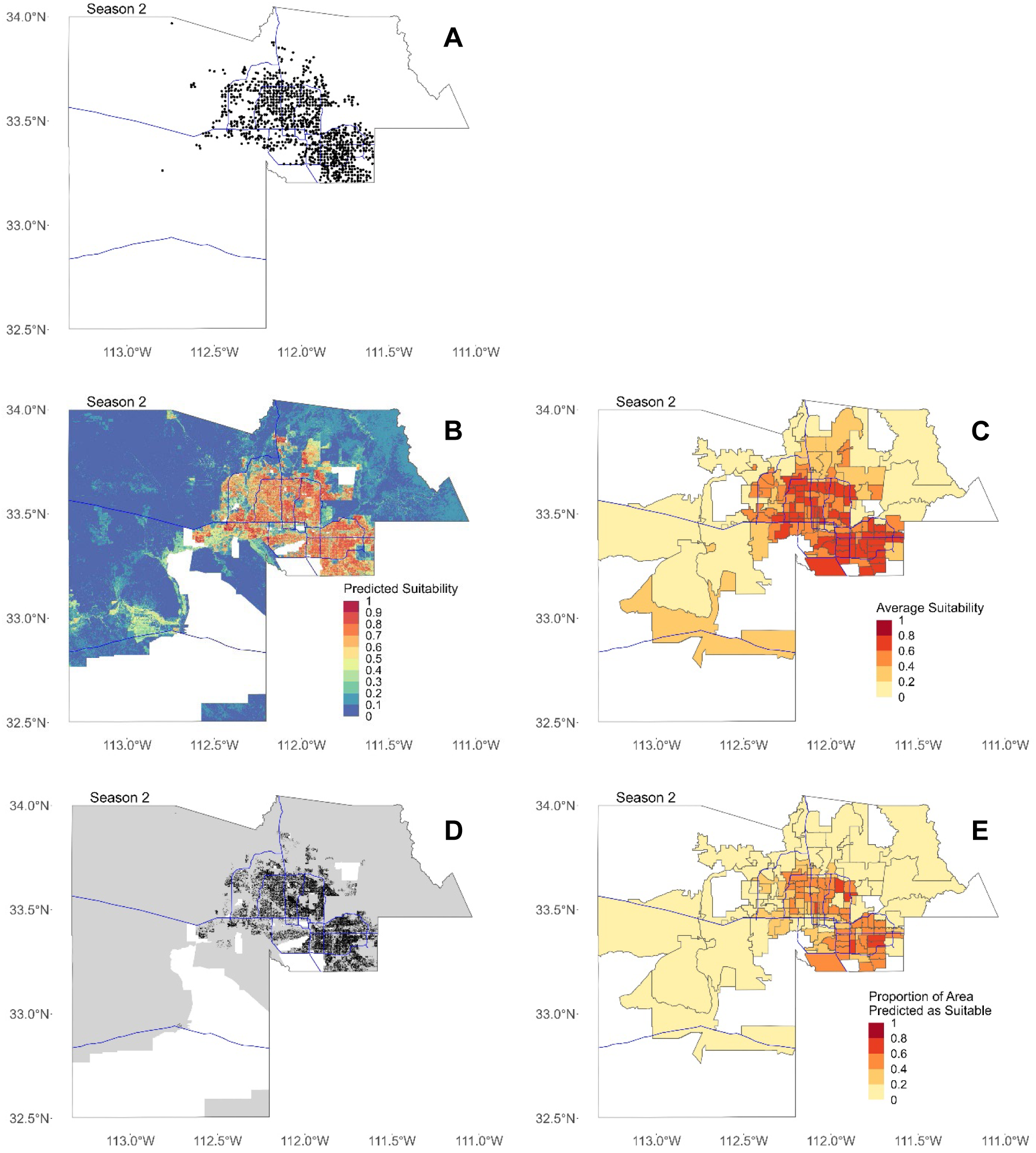
Maps for Season 2 (5/1-7/31 quarters). In all figures, blue lines represent major freeways in Maricopa County as a spatial reference. (**A**) Map of all locations where Ae. aegypti females were trapped in Maricopa County in Season 2. (**B**) Map showing mean predicted suitability for 30-meter by 30-meter pixels. (**C**) Mean predicted suitability averaged by ZCTA. (**D**) Binary map showing areas predicted as suitable for all quarters in black and not suitable in gray. In (B) and (C), areas with missing data where suitability was not predicted are shown in white. (**E**) Proportion of area in each ZCTA predicted as suitable after converting to binary map.

**Figure S28.**
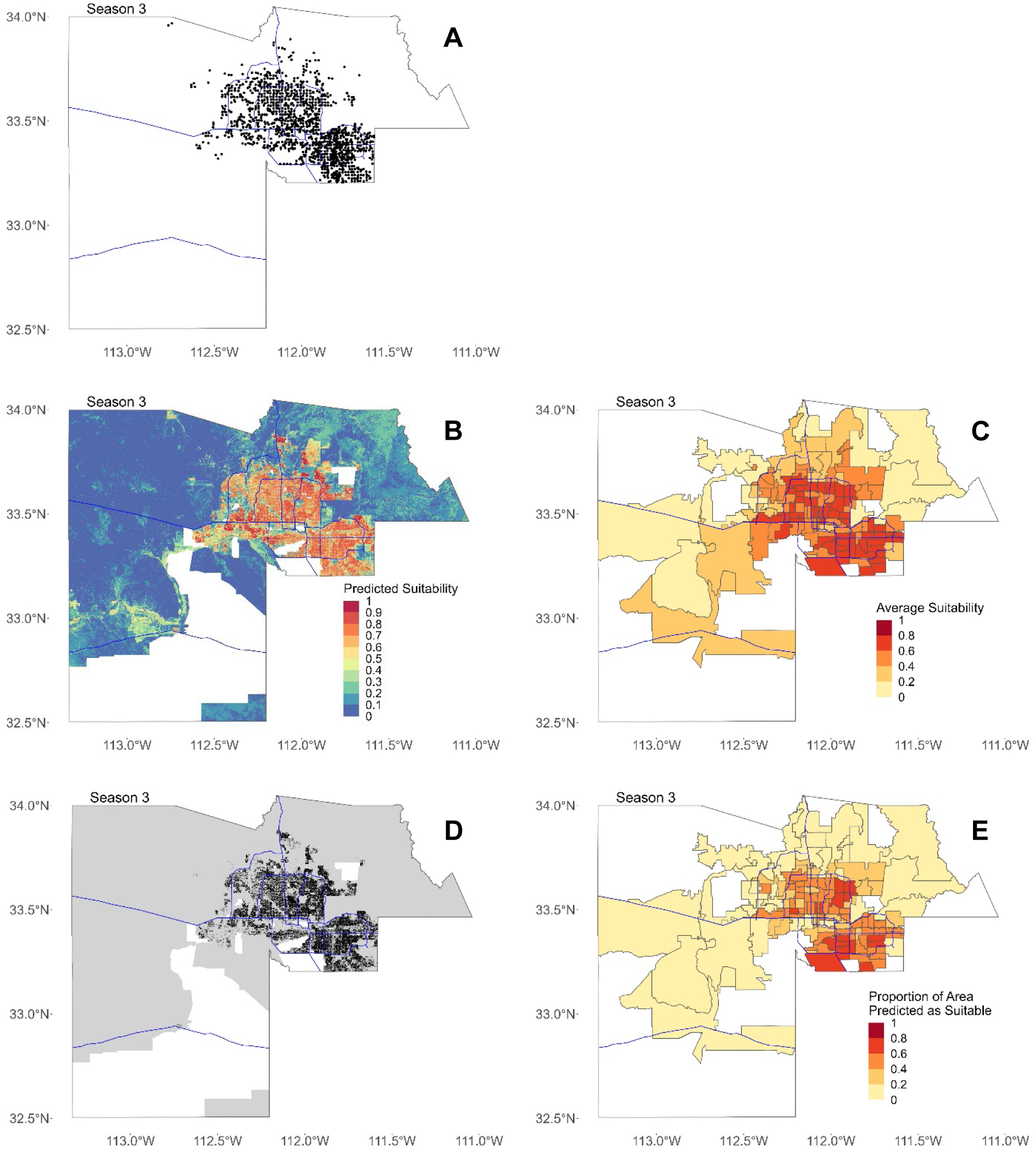
Maps for Season 3 (8/1-10/31 quarters). In all figures, blue lines represent major freeways in Maricopa County as a spatial reference. (**A**) Map of all locations where Ae. aegypti females were trapped in Maricopa County in Season 3. (**B**) Map showing mean predicted suitability for 30-meter by 30-meter pixels. (**C**) Mean predicted suitability averaged by ZCTA. (**D**) Binary map showing areas predicted as suitable for all quarters in black and not suitable in gray. In (B) and (C), areas with missing data where suitability was not predicted are shown in white. (**E**) Proportion of area in each ZCTA predicted as suitable after converting to binary map.

**Figure S29.**
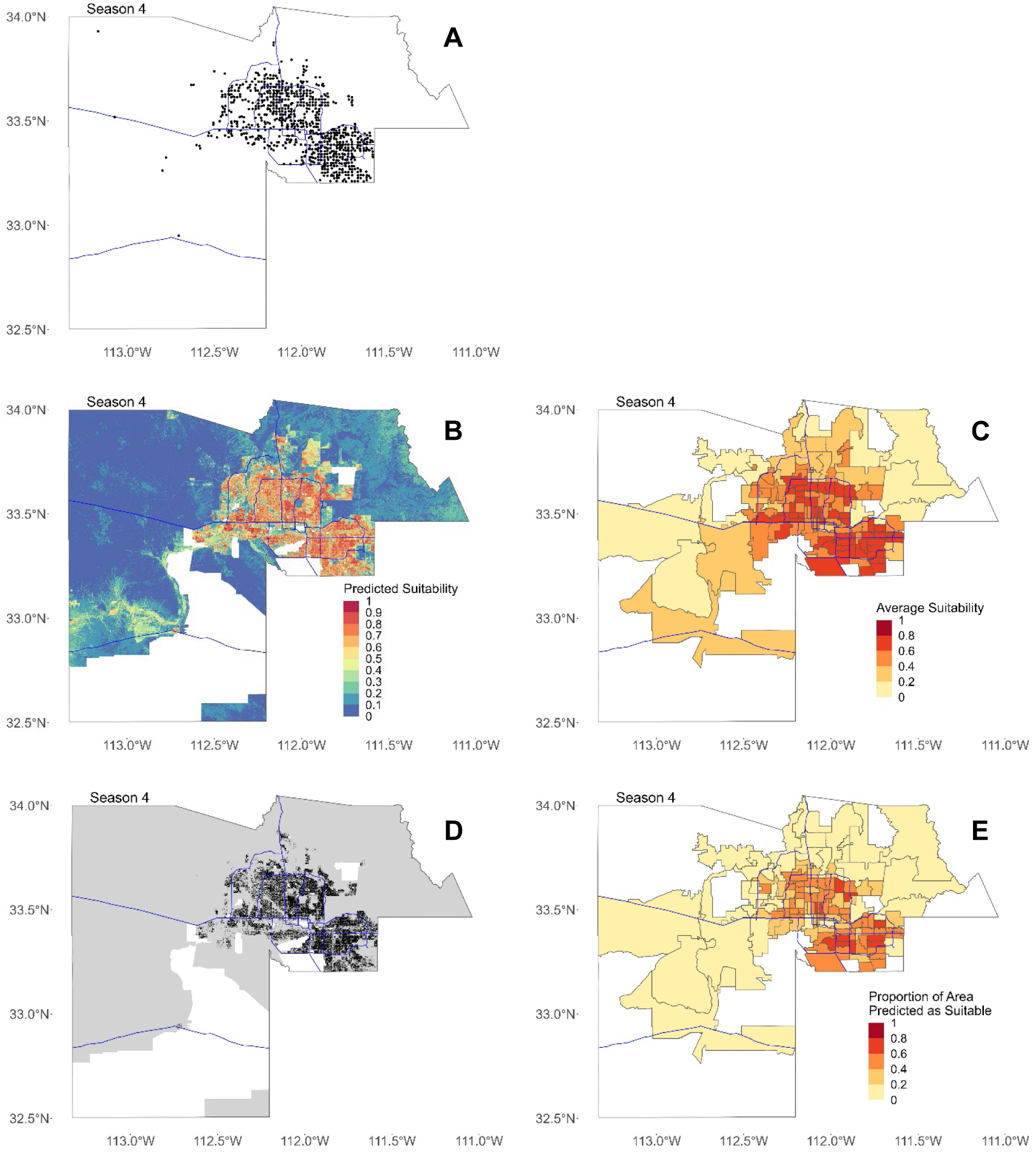
Maps for Season 4 (11/1-1/31 quarters). In all figures, blue lines represent major freeways in Maricopa County as a spatial reference. (**A**) Map of all locations where Ae. aegypti females were trapped in Maricopa County in Season 4. (**B**) Map showing mean predicted suitability for 30-meter by 30-meter pixels. (**C**) Mean predicted suitability averaged by ZCTA. (**D**) Binary map showing areas predicted as suitable for all quarters in black and not suitable in gray. In (B) and (C), areas with missing data where suitability was not predicted are shown in white. (**E**) Proportion of area in each ZCTA predicted as suitable after converting to binary map.

**Figure S30.**
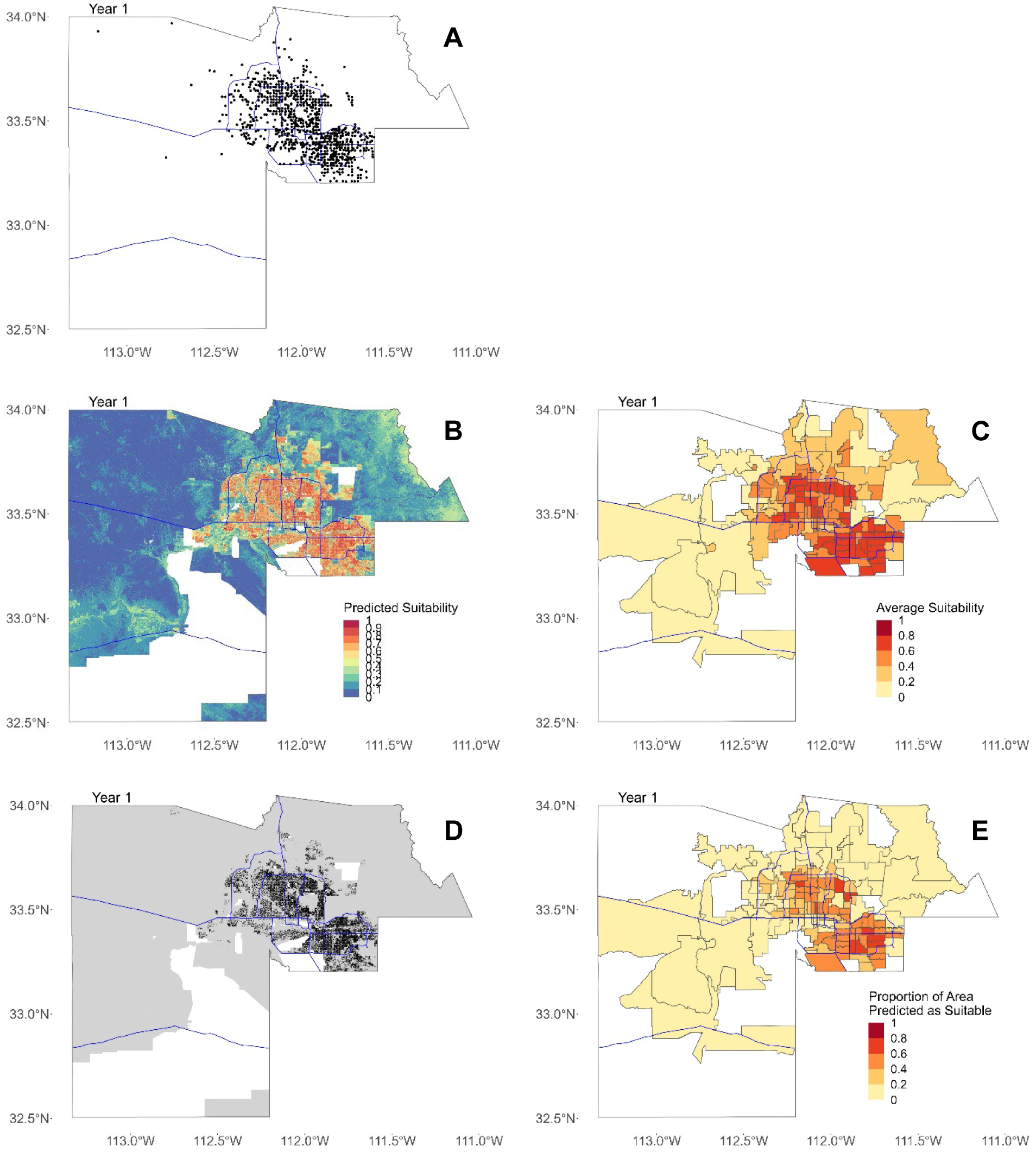
Maps for Year 1 (2/1/14-1/31/15). In all figures, blue lines represent major freeways in Maricopa County as a spatial reference. (**A**) Map of all locations where Ae. aegypti females were trapped in Maricopa County in Year 1. (**B**) Map showing mean predicted suitability for 30-meter by 30-meter pixels. (**C**) Mean predicted suitability averaged by ZCTA. (**D**) Binary map showing areas predicted as suitable for all quarters in black and not suitable in gray. In (B) and (C), areas with missing data where suitability was not predicted are shown in white. (**E**) Proportion of area in each ZCTA predicted as suitable after converting to binary map.

**Figure S31.**
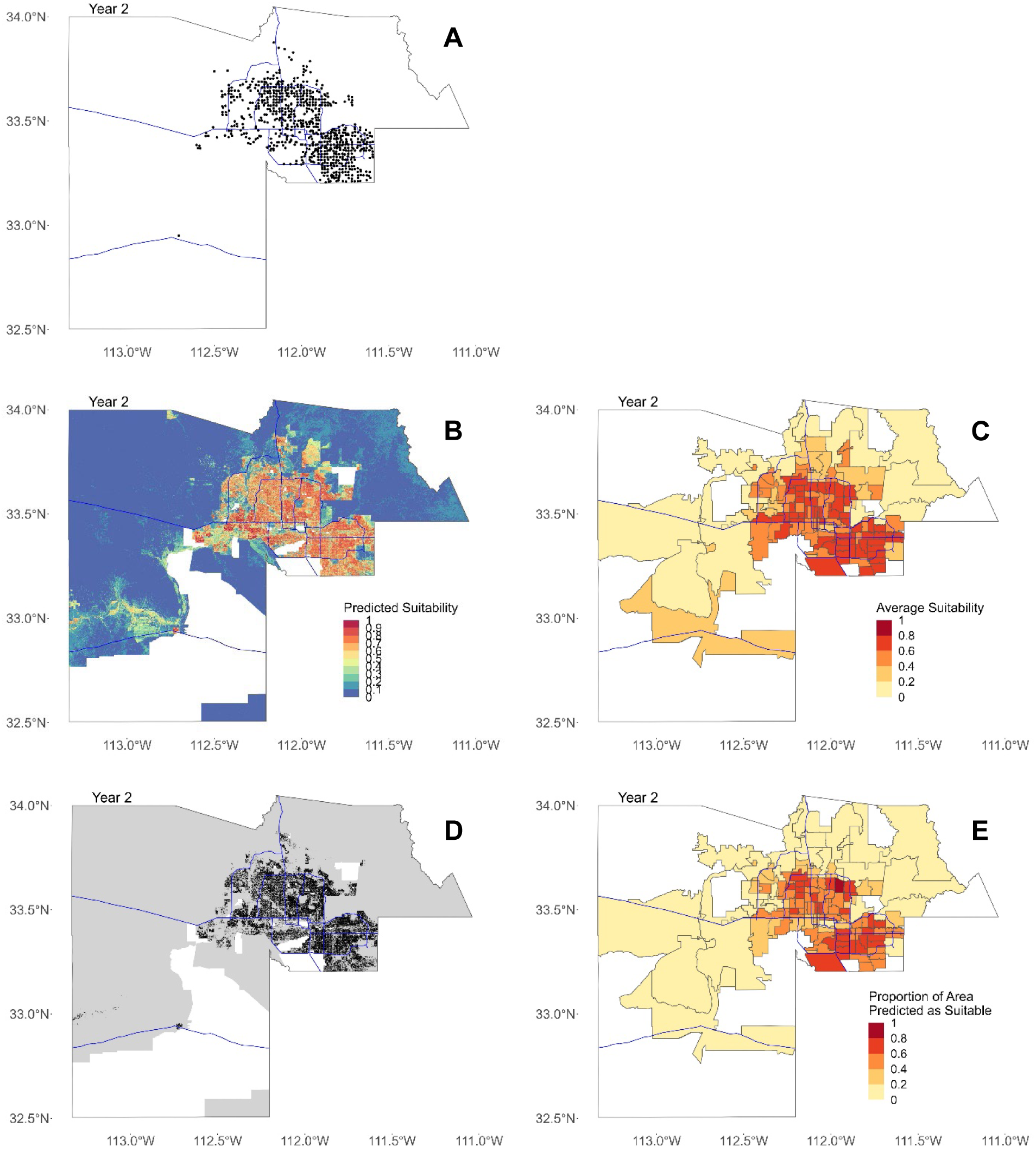
Maps for Year 2 (2/1/15-1/31/16). In all figures, blue lines represent major freeways in Maricopa County as a spatial reference. (**A**) Map of all locations where Ae. aegypti females were trapped in Maricopa County in Year 2. (**B**) Map showing mean predicted suitability for 30-meter by 30-meter pixels. (**C**) Mean predicted suitability averaged by ZCTA. (**D**) Binary map showing areas predicted as suitable for all quarters in black and not suitable in gray. In (B) and (C), areas with missing data where suitability was not predicted are shown in white. (**E**) Proportion of area in each ZCTA predicted as suitable after converting to binary map.

**Figure S32.**
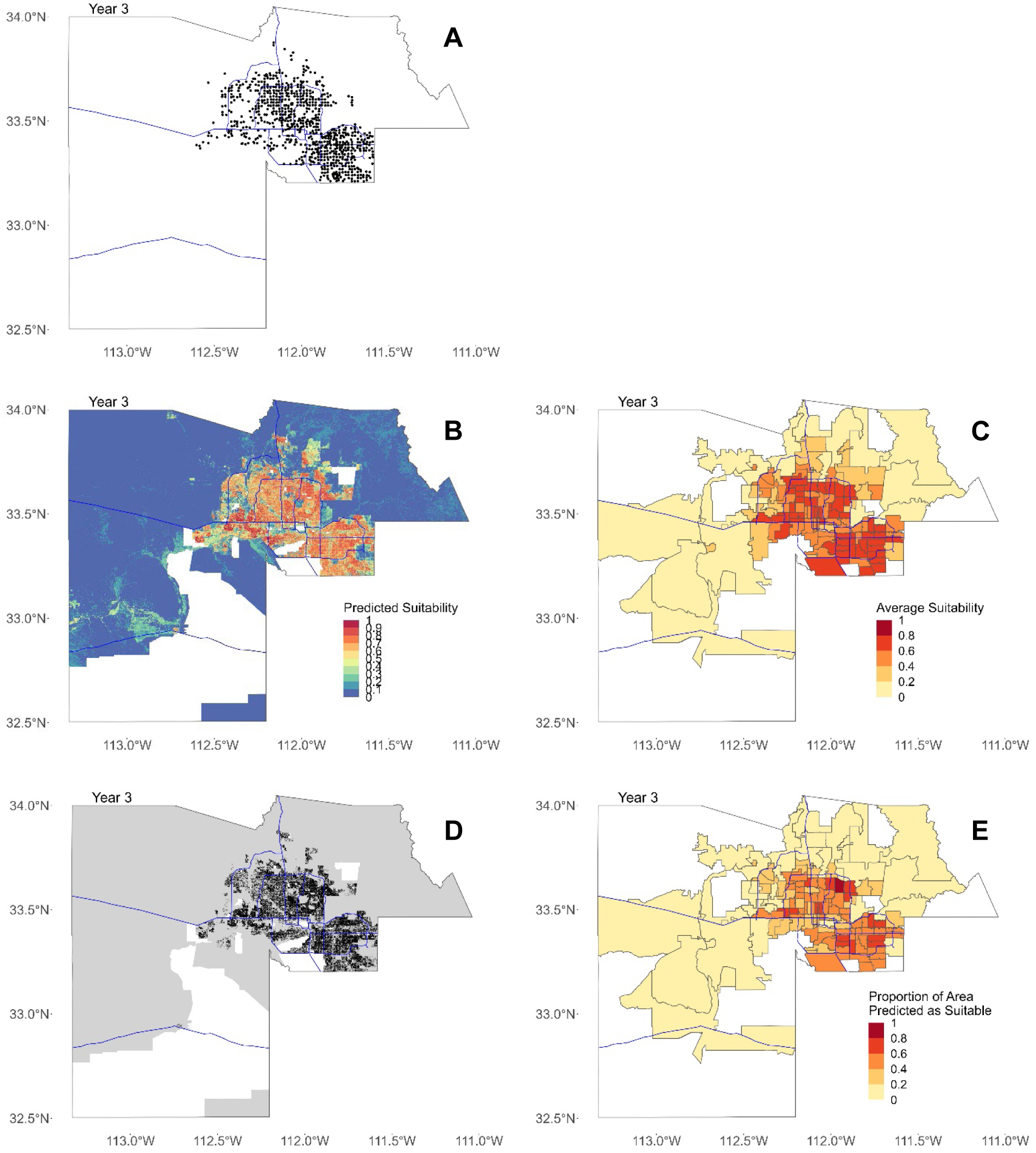
Maps for Year 3 (2/1/16-1/31/17). In all figures, blue lines represent major freeways in Maricopa County as a spatial reference. (**A**) Map of all locations where Ae. aegypti females were trapped in Maricopa County in Year 3. (**B**) Map showing mean predicted suitability for 30-meter by 30-meter pixels. (**C**) Mean predicted suitability averaged by ZCTA. (**D**) Binary map showing areas predicted as suitable for all quarters in black and not suitable in gray. In (B) and (C), areas with missing data where suitability was not predicted are shown in white. (**E**) Proportion of area in each ZCTA predicted as suitable after converting to binary map.

**Figure S33.**
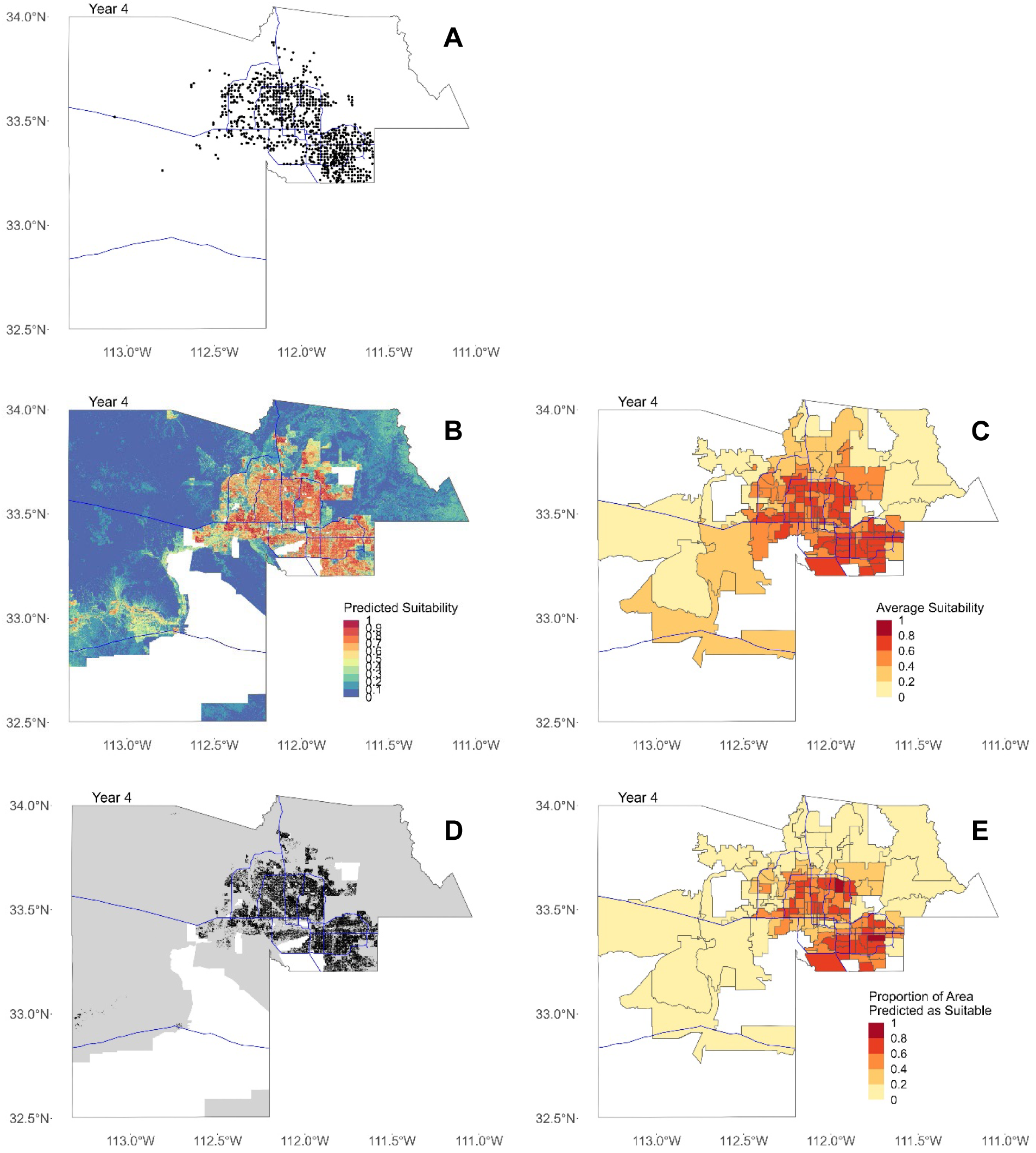
Maps for Year 4 (2/1/17-1/31/18). In all figures, blue lines represent major freeways in Maricopa County as a spatial reference. (**A**) Map of all locations where Ae. aegypti females were trapped in Maricopa County in Year 4. (**B**) Map showing mean predicted suitability for 30-meter by 30-meter pixels. (**C**) Mean predicted suitability averaged by ZCTA. (**D**) Binary map showing areas predicted as suitable for all quarters in black and not suitable in gray. In (B) and (C), areas with missing data where suitability was not predicted are shown in white. (**E**) Proportion of area in each ZCTA predicted as suitable after converting to binary map.

**Figure S34.**
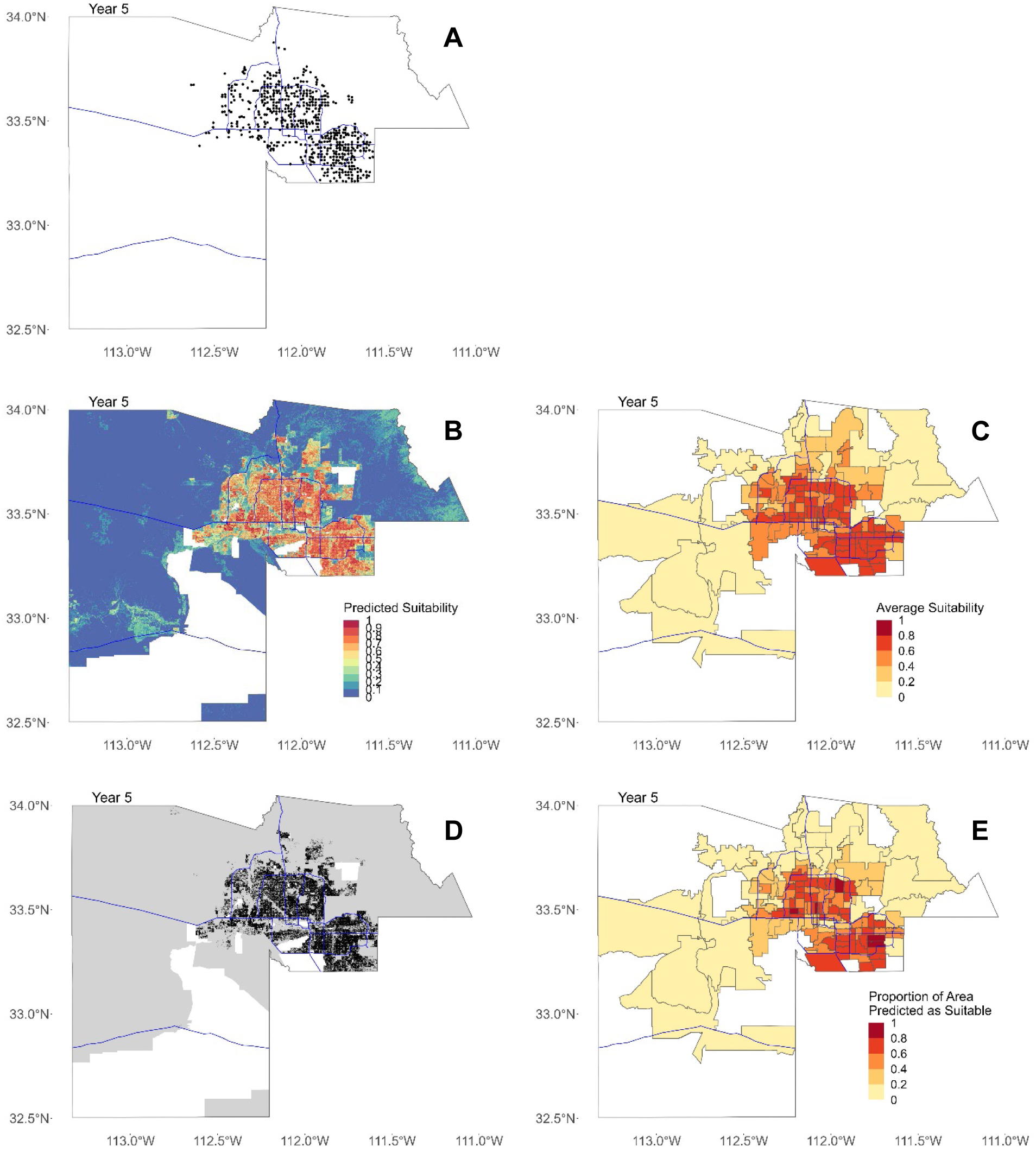
Maps for Year 5 (2/1/18-7/31/18). In all figures, blue lines represent major freeways in Maricopa County as a spatial reference. (**A**) Map of all locations where Ae. aegypti females were trapped in Maricopa County in Year 5. (**B**) Map showing mean predicted suitability for 30-meter by 30-meter pixels. (**C**) Mean predicted suitability averaged by ZCTA. (**D**) Binary map showing areas predicted as suitable for all quarters in black and not suitable in gray. In (B) and (C), areas with missing data where suitability was not predicted are shown in white. (**E**) Proportion of area in each ZCTA predicted as suitable after converting to binary map.

**Figure S35.**
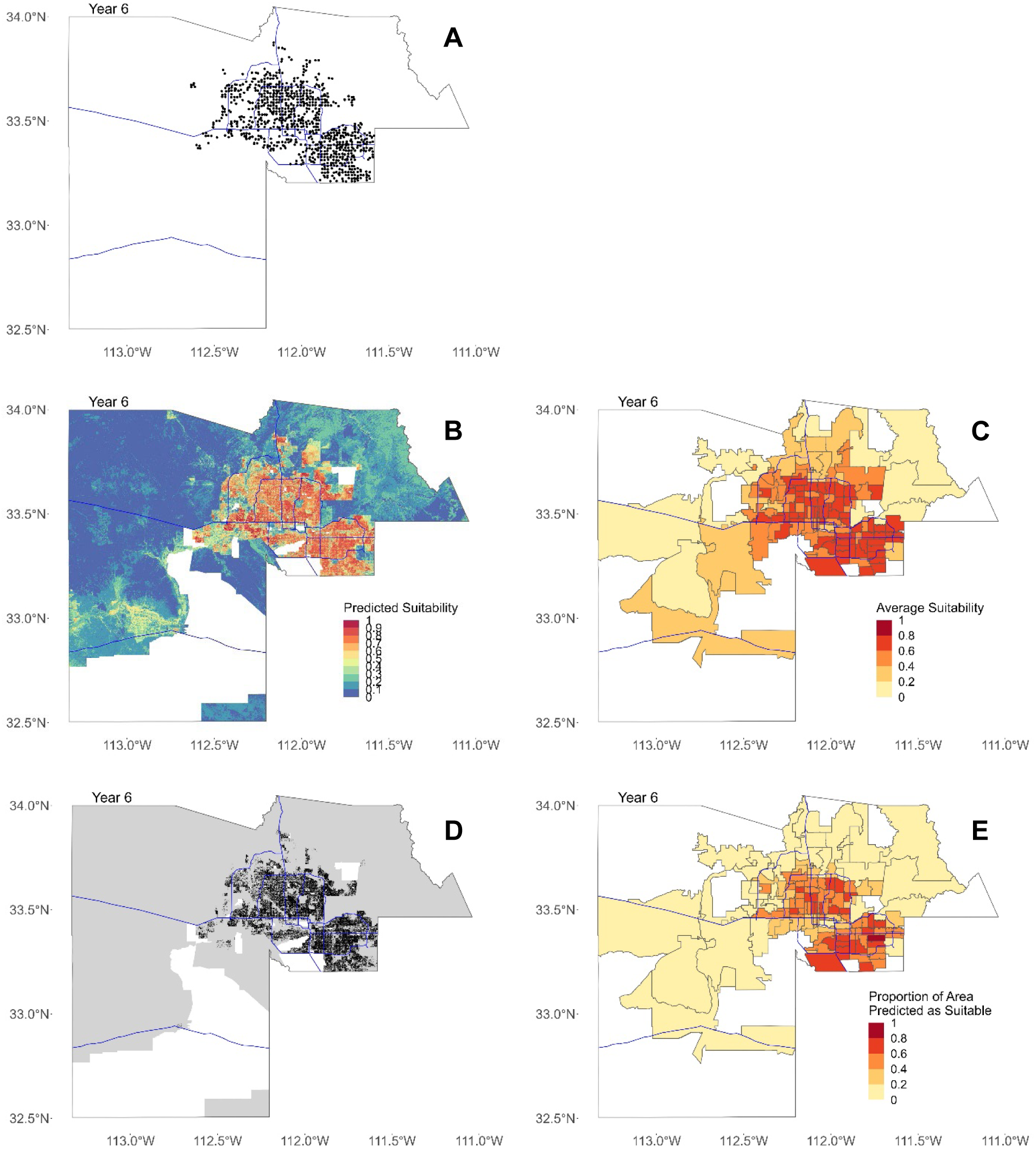
Maps for Year 6 (2/1/19-1/31/20). In all figures, blue lines represent major freeways in Maricopa County as a spatial reference. (**A**) Map of all locations where Ae. aegypti females were trapped in Maricopa County in Year 6. (**B**) Map showing mean predicted suitability for 30-meter by 30-meter pixels. (**C**) Mean predicted suitability averaged by ZCTA. (**D**) Binary map showing areas predicted as suitable for all quarters in black and not suitable in gray. In (B) and (C), areas with missing data where suitability was not predicted are shown in white. (**E**) Proportion of area in each ZCTA predicted as suitable after converting to binary map.

**Figure S36.**
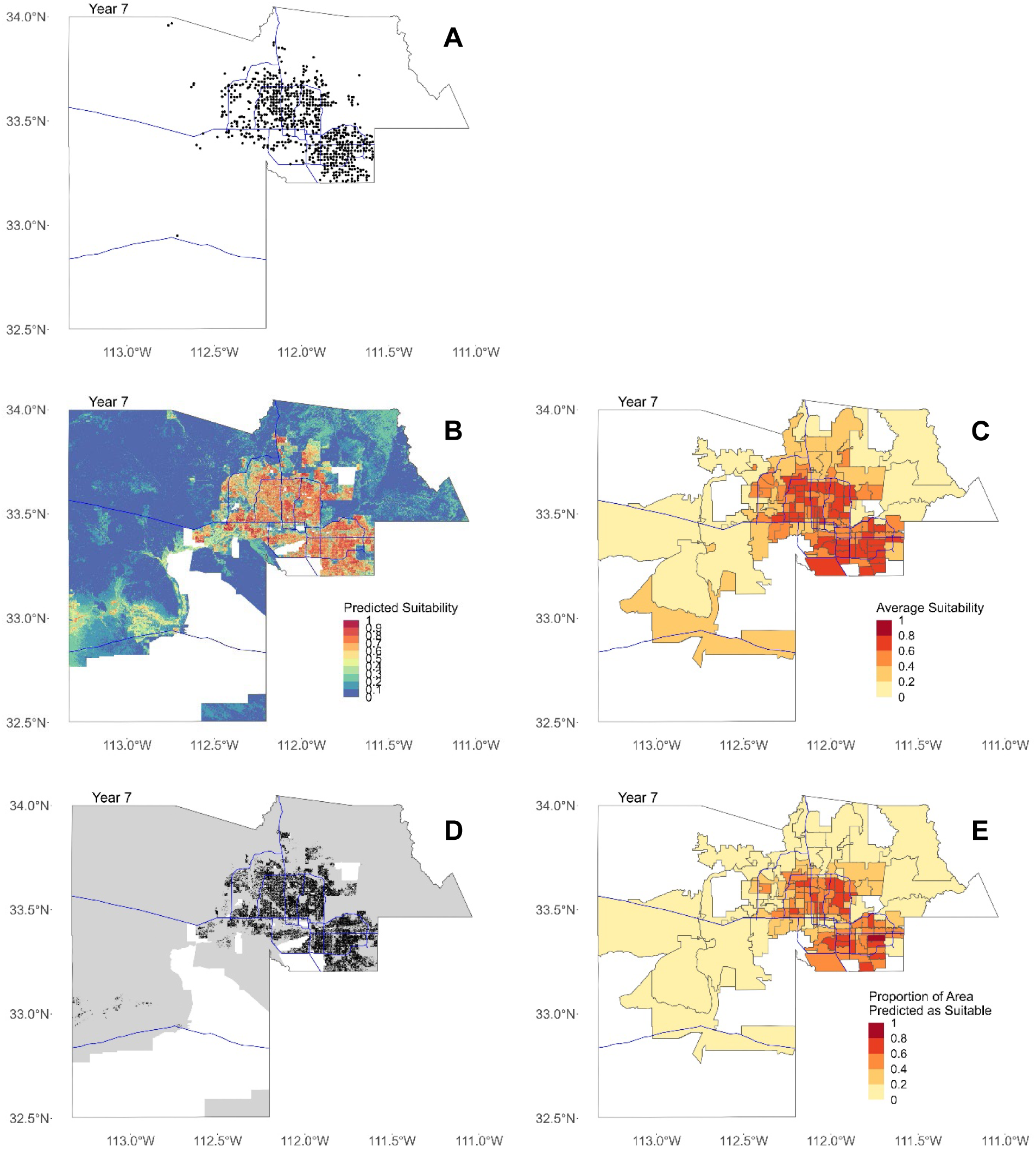
Maps for Year 7 (2/1/20-10/31/20). In all figures, blue lines represent major freeways in Maricopa County as a spatial reference. (**A**) Map of all locations where Ae. aegypti females were trapped in Maricopa County in Year 7. (**B**) Map showing mean predicted suitability for 30-meter by 30-meter pixels. (**C**) Mean predicted suitability averaged by ZCTA. (**D**) Binary map showing areas predicted as suitable for all quarters in black and not suitable in gray. In (B) and (C), areas with missing data where suitability was not predicted are shown in white. (**E**) Proportion of area in each ZCTA predicted as suitable after converting to binary map.

**Figure S37.**
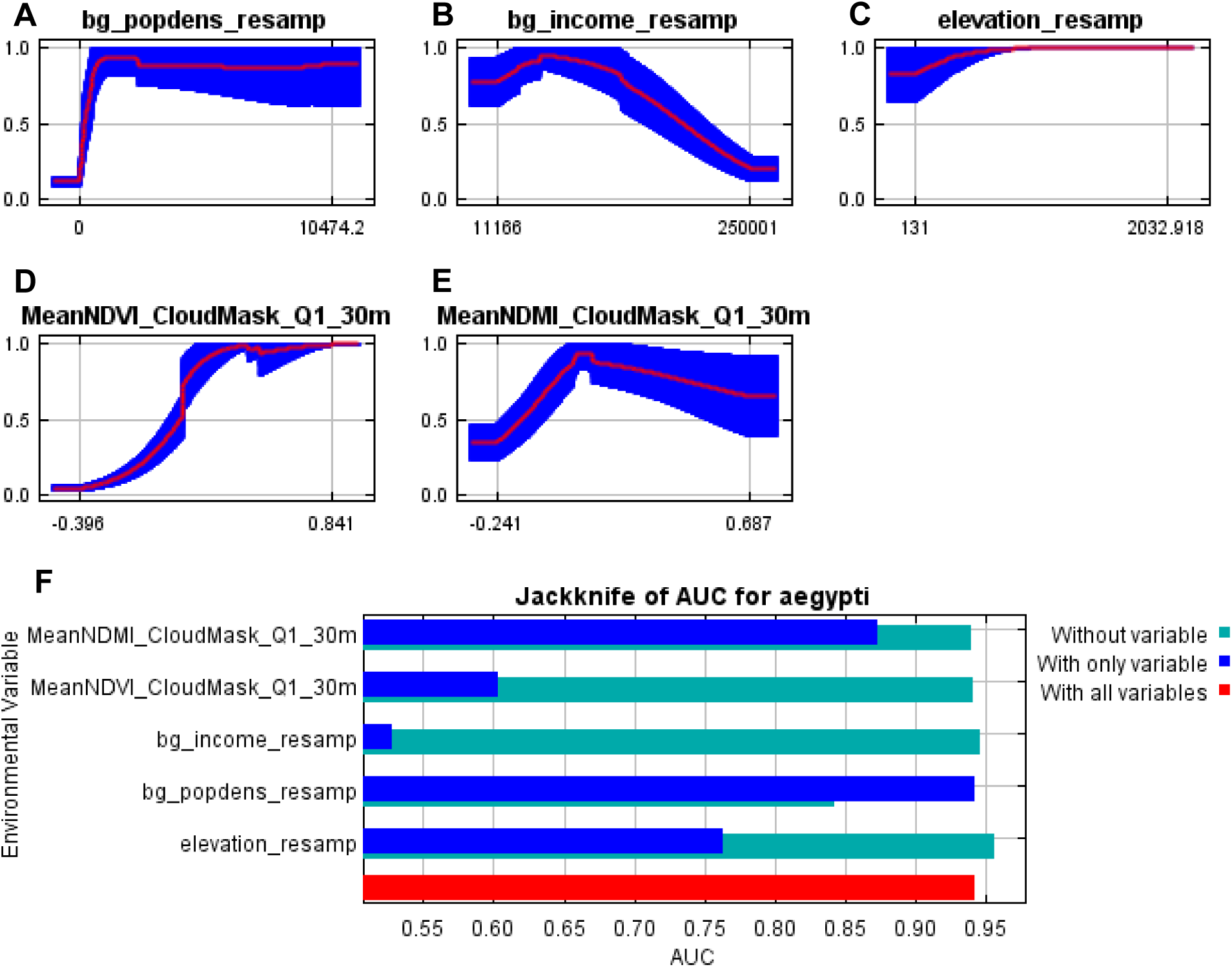
Response curves for Quarter 1 (2/1/14—4/30/14). (**A**) through (**E**) show Maxent response curves for the predictors Population Density, Median Income, Elevation, mean Normalized Difference Vegetation Index (NDVI), and mean Normalized Difference Moisture Index (NDMI). Response curves are shown for each predictor when all other predictors are held constant. Mean response from 10 folds is shown with a red line, and +/- one standard deviation is shown in blue. (**F**) Maxent output showing the average change in the Area Under Curve (AUC) value when only that predictor is included (dark blue), when only that predictor is removed (green), and when all predictors were included (red).

**Figure S38.**
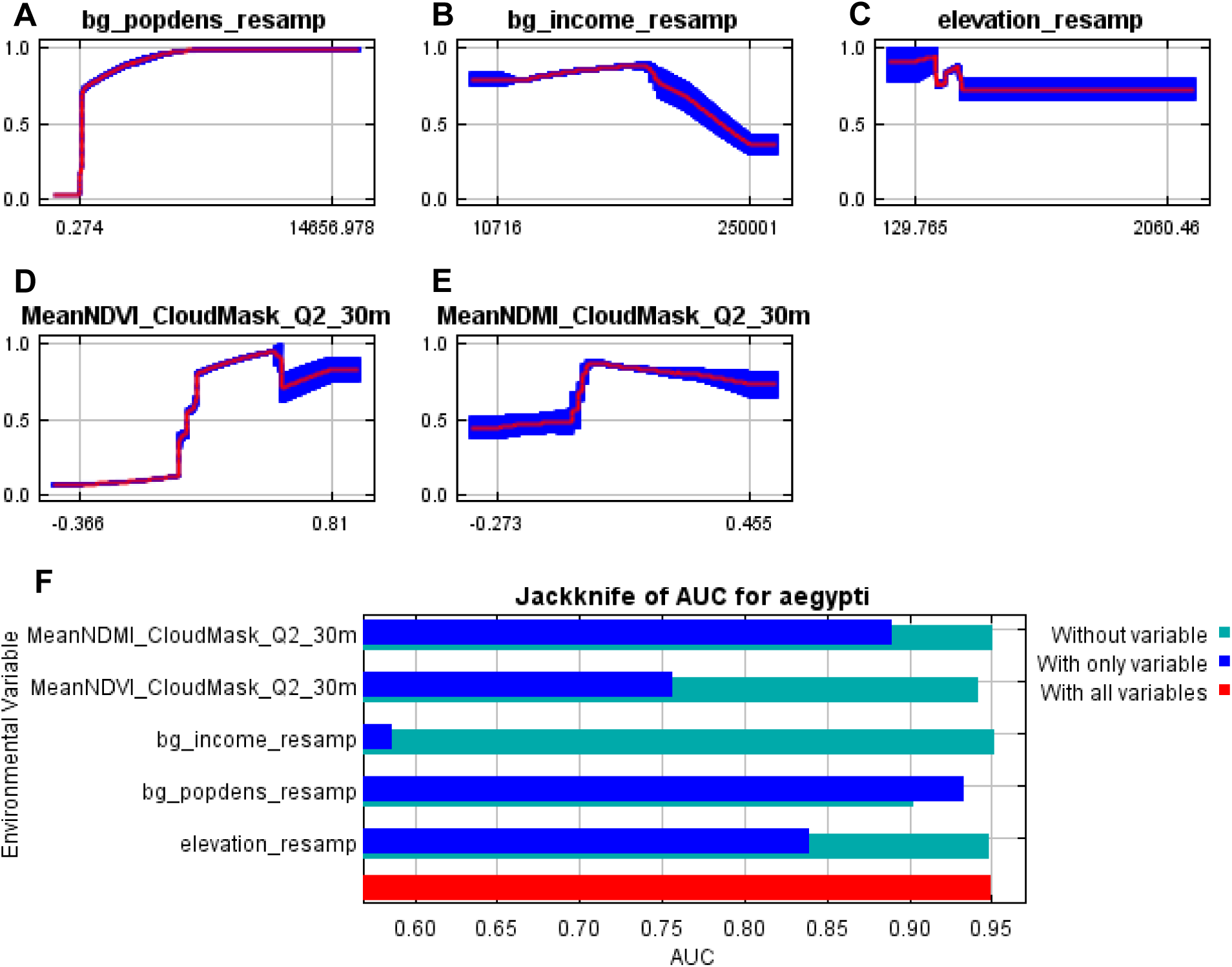
Response curves for Quarter 2 (5/1/14—7/31/14). (**A**) through (**E**) show Maxent response curves for the predictors Population Density, Median Income, Elevation, mean Normalized Difference Vegetation Index (NDVI), and mean Normalized Difference Moisture Index (NDMI). Response curves are shown for each predictor when all other predictors are held constant. Mean response from 10 folds is shown with a red line, and +/- one standard deviation is shown in blue. (**F**) Maxent output showing the average change in the Area Under Curve (AUC) value when only that predictor is included (dark blue), when only that predictor is removed (green), and when all predictors were included (red).

**Figure S39.**
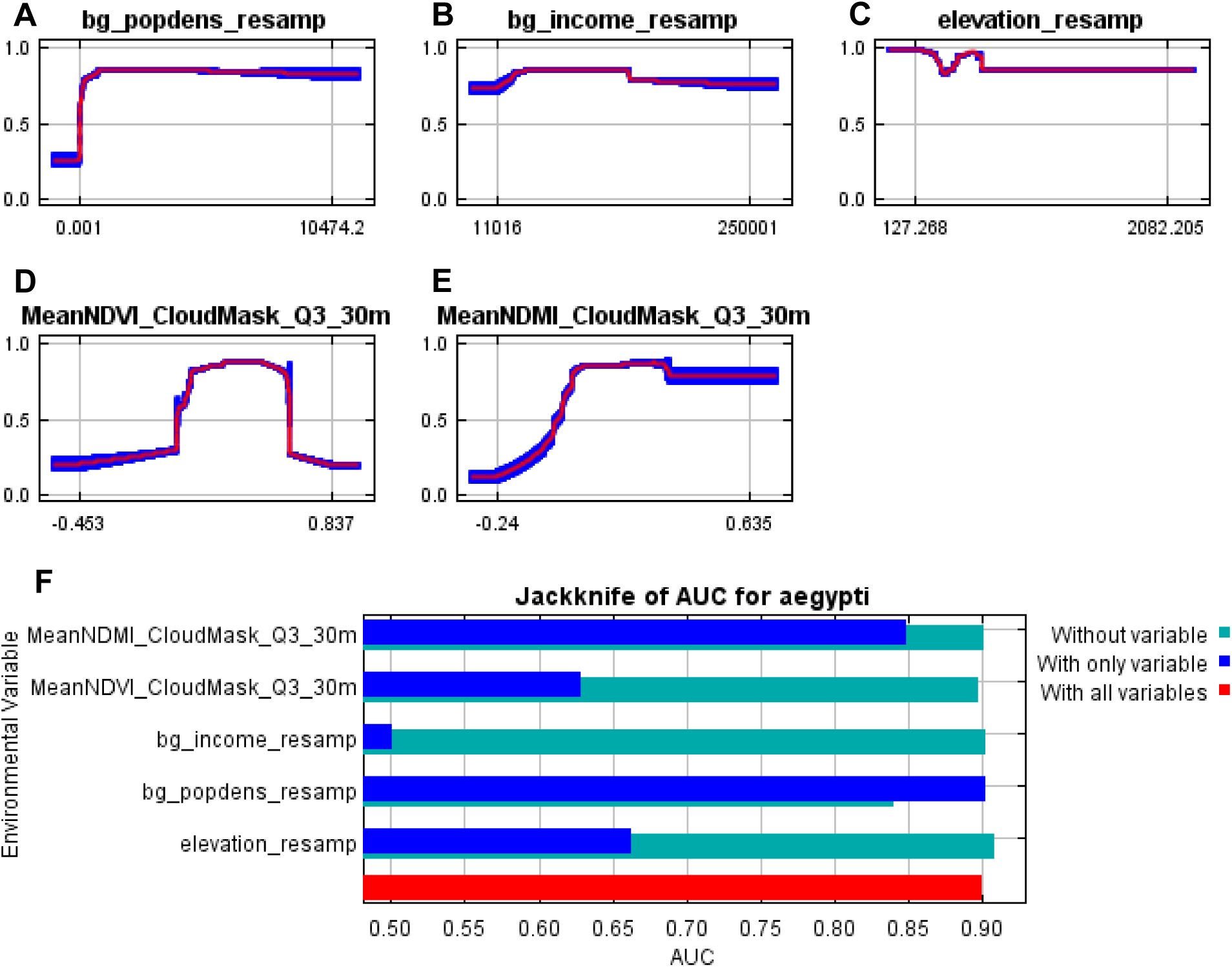
Response curves for Quarter 3 (8/1/14—10/31/14). (**A**) through (**E**) show Maxent response curves for the predictors Population Density, Median Income, Elevation, mean Normalized Difference Vegetation Index (NDVI), and mean Normalized Difference Moisture Index (NDMI). Response curves are shown for each predictor when all other predictors are held constant. Mean response from 10 folds is shown with a red line, and +/- one standard deviation is shown in blue. (**F**) Maxent output showing the average change in the Area Under Curve (AUC) value when only that predictor is included (dark blue), when only that predictor is removed (green), and when all predictors were included (red).

**Figure S40.**
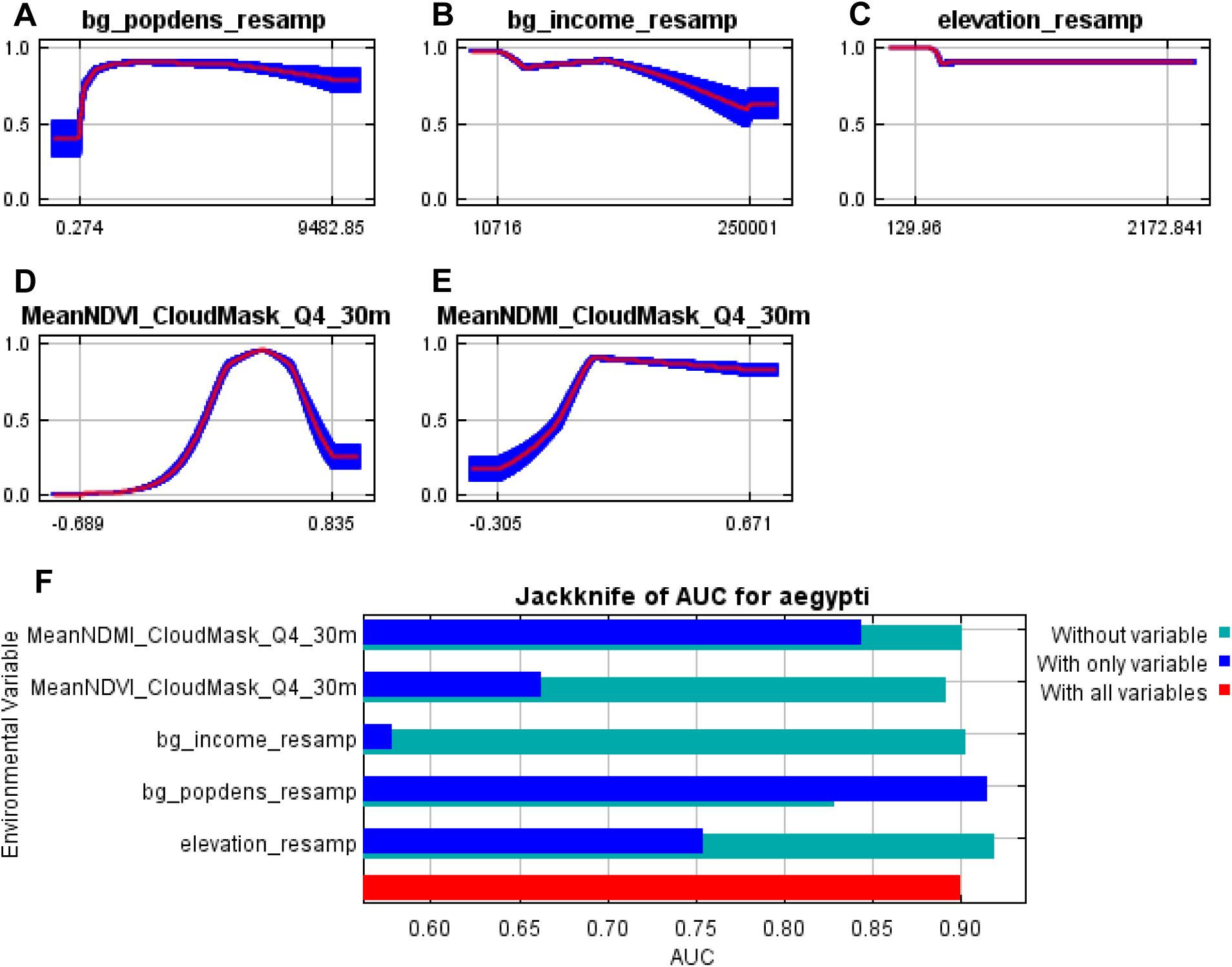
Response curves for Quarter 4 (11/1/14—1/31/15). (**A**) through (**E**) show Maxent response curves for the predictors Population Density, Median Income, Elevation, mean Normalized Difference Vegetation Index (NDVI), and mean Normalized Difference Moisture Index (NDMI). Response curves are shown for each predictor when all other predictors are held constant. Mean response from 10 folds is shown with a red line, and +/- one standard deviation is shown in blue. (**F**) Maxent output showing the average change in the Area Under Curve (AUC) value when only that predictor is included (dark blue), when only that predictor is removed (green), and when all predictors were included (red).

**Figure S41.**
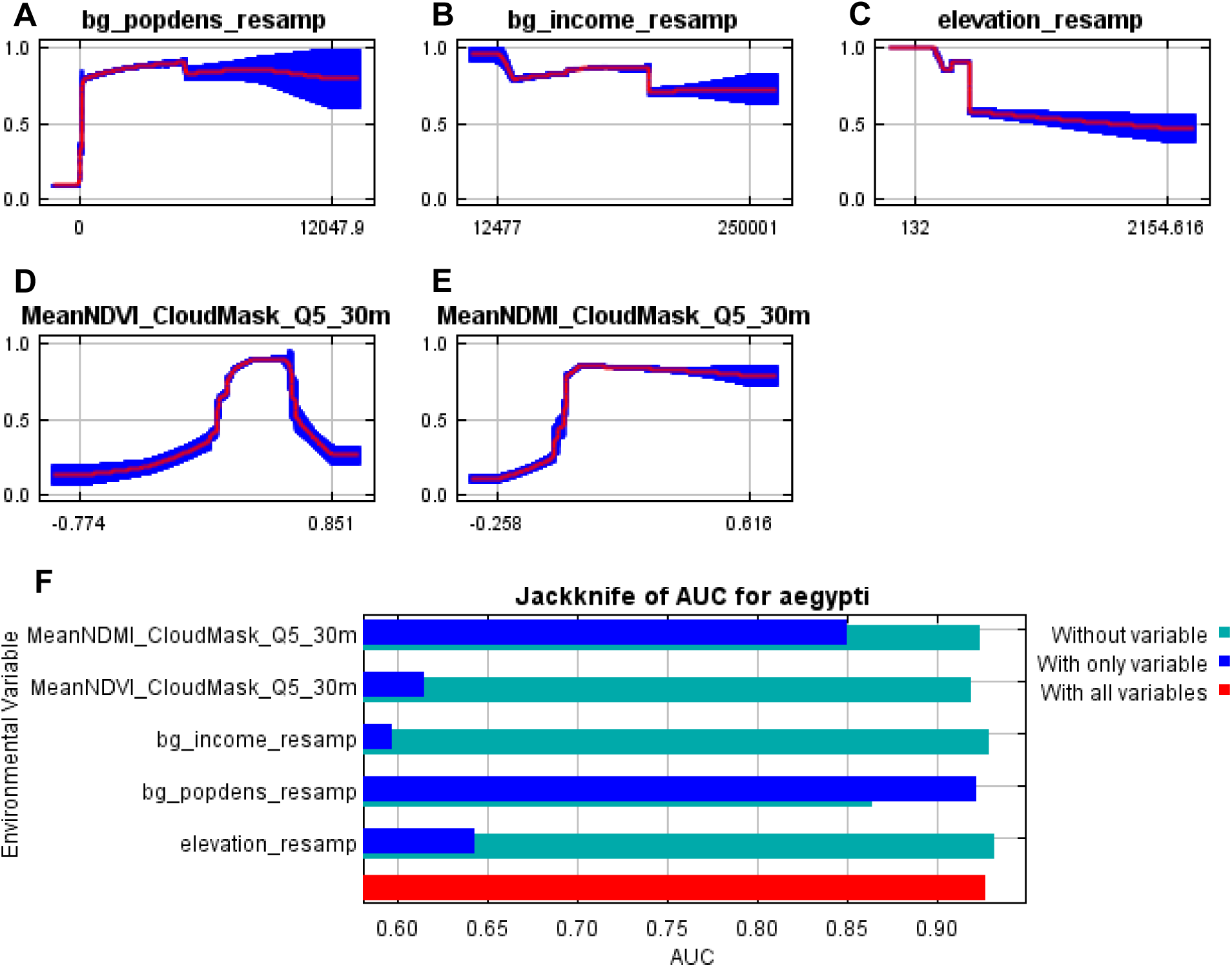
Response curves for Quarter 5 (2/1/15—4/30/15). (**A**) through (**E**) show Maxent response curves for the predictors Population Density, Median Income, Elevation, mean Normalized Difference Vegetation Index (NDVI), and mean Normalized Difference Moisture Index (NDMI). Response curves are shown for each predictor when all other predictors are held constant. Mean response from 10 folds is shown with a red line, and +/- one standard deviation is shown in blue. (**F**) Maxent output showing the average change in the Area Under Curve (AUC) value when only that predictor is included (dark blue), when only that predictor is removed (green), and when all predictors were included (red).

**Figure S42.**
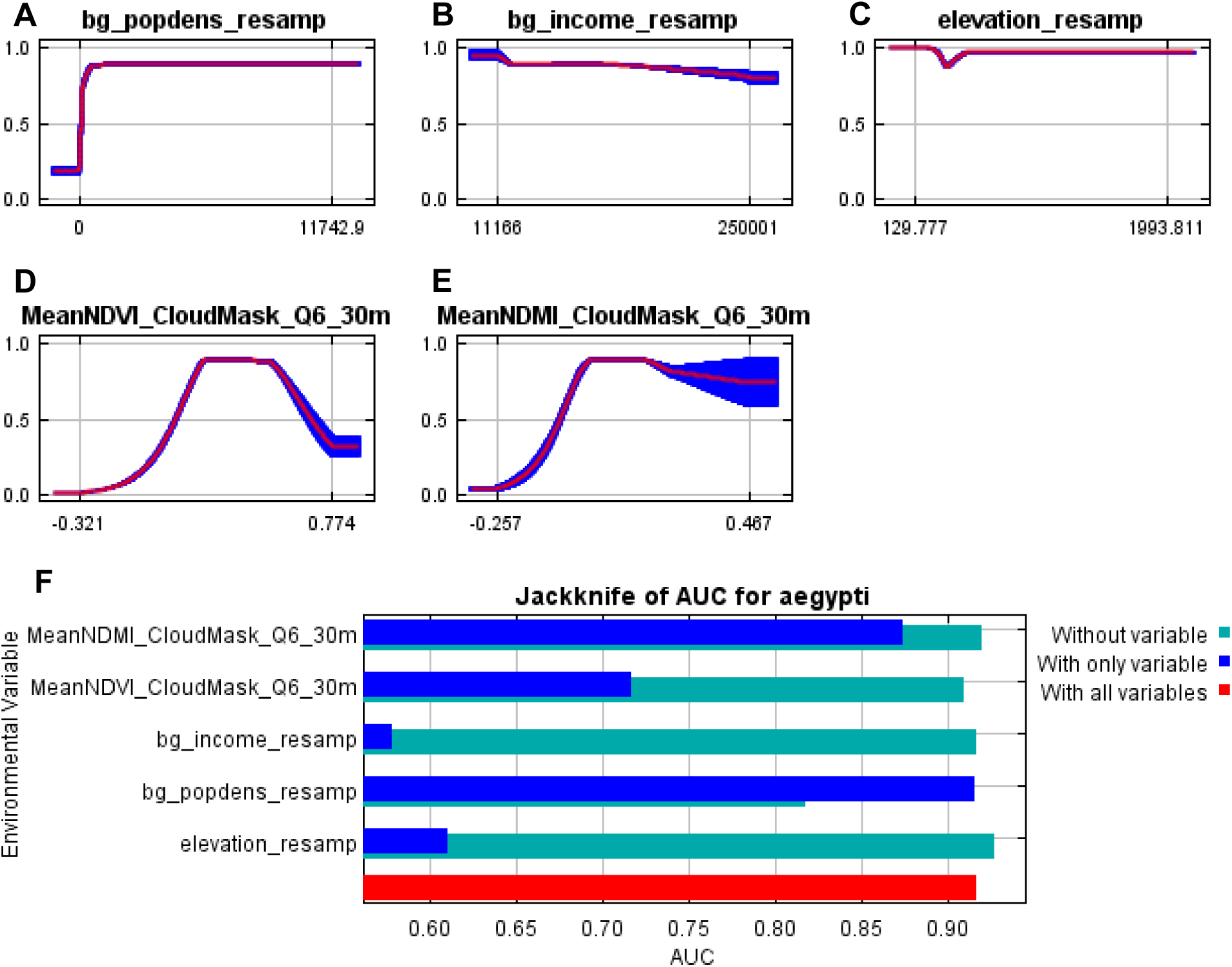
Response curves for Quarter 6 (5/1/15—7/31/15). (**A**) through (**E**) show Maxent response curves for the predictors Population Density, Median Income, Elevation, mean Normalized Difference Vegetation Index (NDVI), and mean Normalized Difference Moisture Index (NDMI). Response curves are shown for each predictor when all other predictors are held constant. Mean response from 10 folds is shown with a red line, and +/- one standard deviation is shown in blue. (**F**) Maxent output showing the average change in the Area Under Curve (AUC) value when only that predictor is included (dark blue), when only that predictor is removed (green), and when all predictors were included (red).

**Figure S43.**
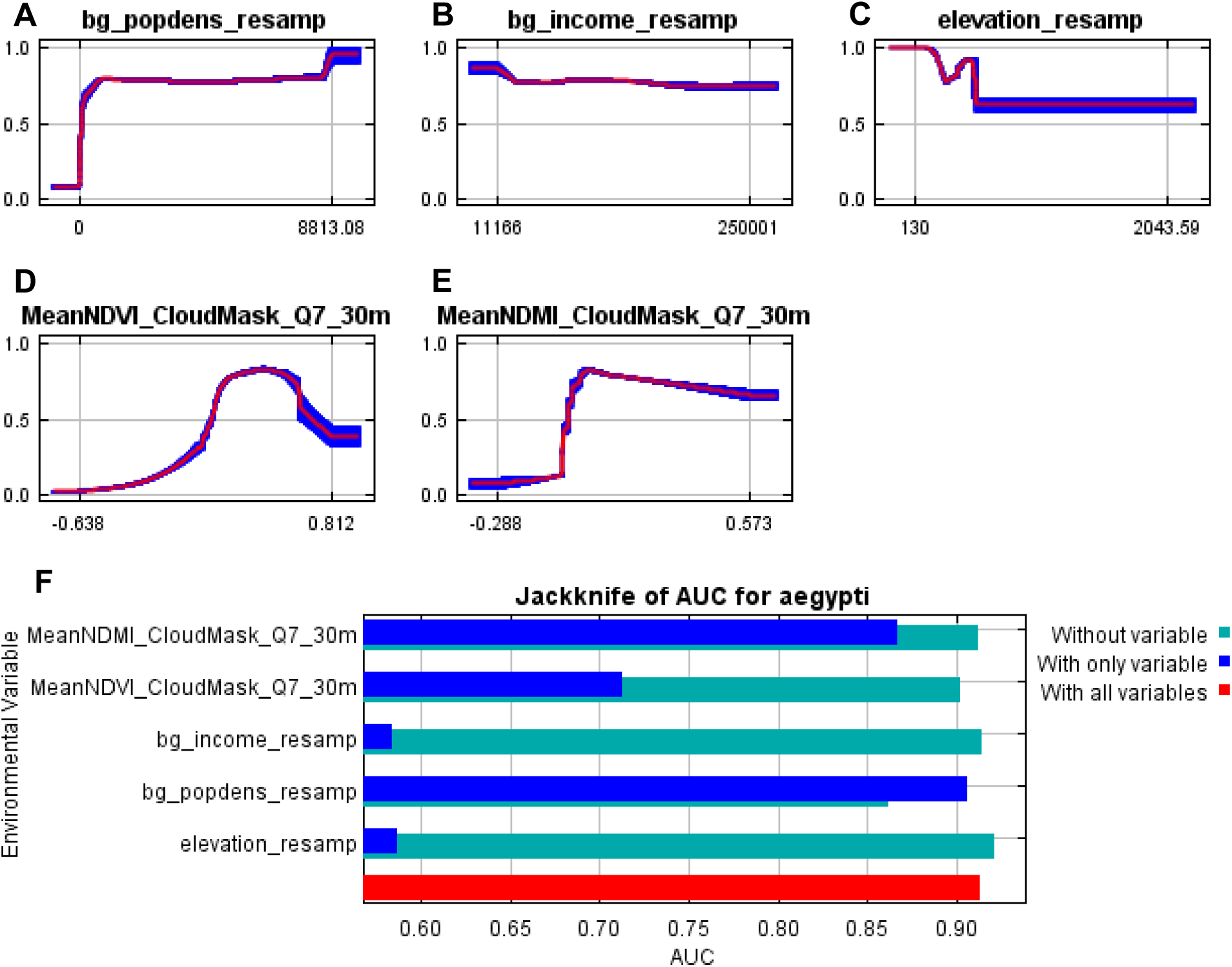
Response curves for Quarter 7 (8/1/15—10/31/15). (**A**) through (**E**) show Maxent response curves for the predictors Population Density, Median Income, Elevation, mean Normalized Difference Vegetation Index (NDVI), and mean Normalized Difference Moisture Index (NDMI). Response curves are shown for each predictor when all other predictors are held constant. Mean response from 10 folds is shown with a red line, and +/- one standard deviation is shown in blue. (**F**) Maxent output showing the average change in the Area Under Curve (AUC) value when only that predictor is included (dark blue), when only that predictor is removed (green), and when all predictors were included (red).

**Figure S44.**
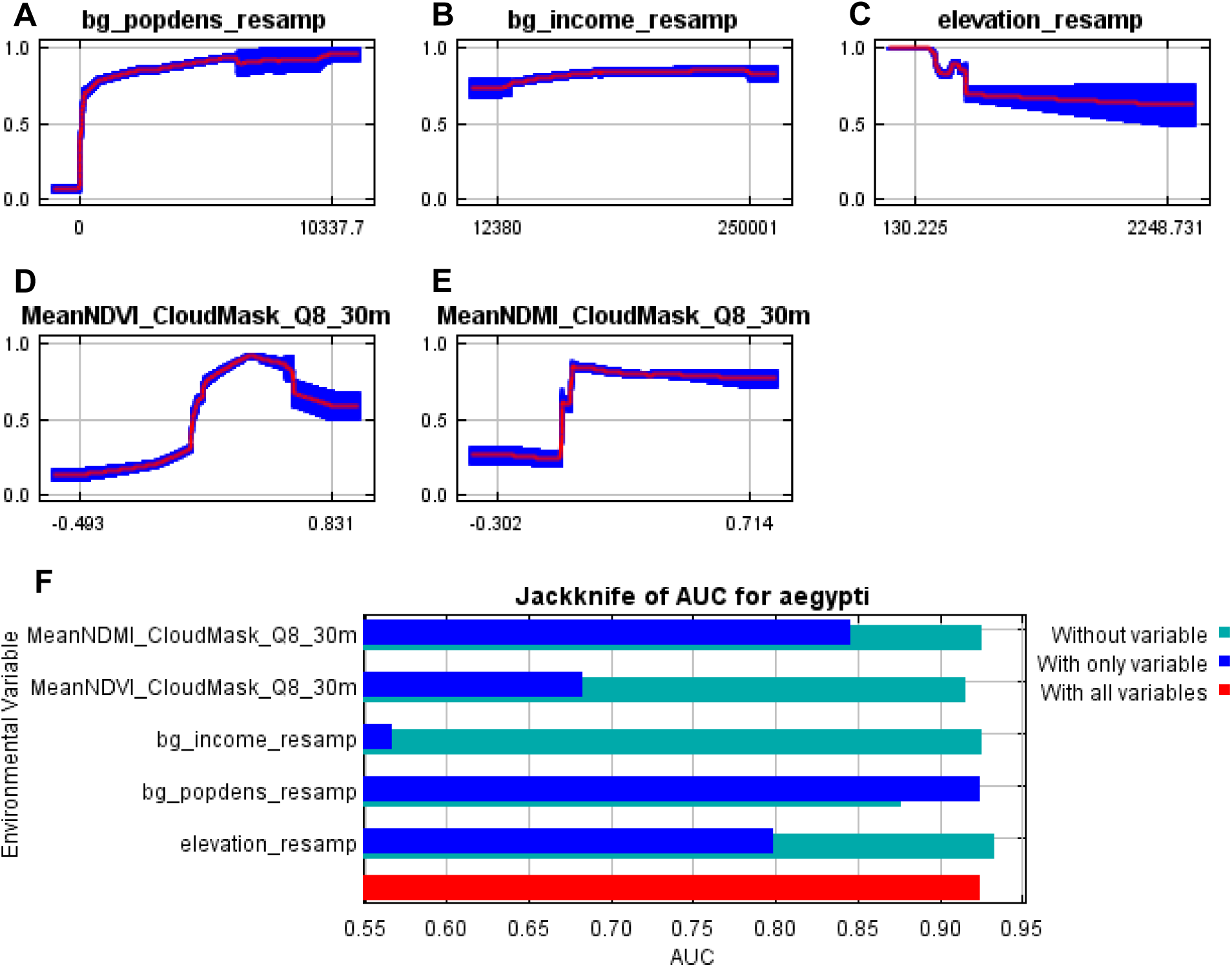
Response curves for Quarter 8 (11/1/15—1/31/16). (**A**) through (**E**) show Maxent response curves for the predictors Population Density, Median Income, Elevation, mean Normalized Difference Vegetation Index (NDVI), and mean Normalized Difference Moisture Index (NDMI). Response curves are shown for each predictor when all other predictors are held constant. Mean response from 10 folds is shown with a red line, and +/- one standard deviation is shown in blue. (**F**) Maxent output showing the average change in the Area Under Curve (AUC) value when only that predictor is included (dark blue), when only that predictor is removed (green), and when all predictors were included (red).

**Figure S45.**
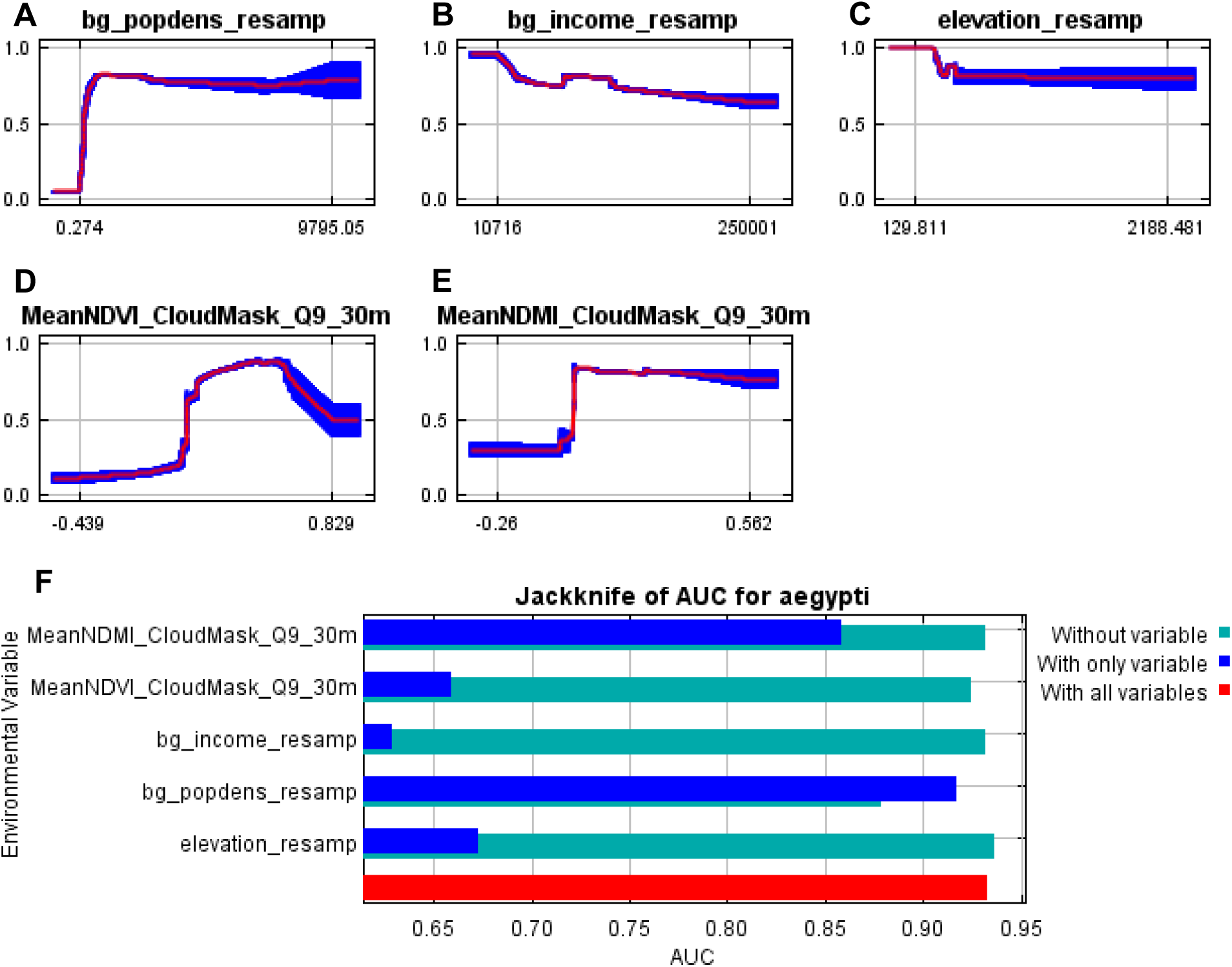
Response curves for Quarter 9 (2/1/16—4/30/16). (**A**) through (**E**) show Maxent response curves for the predictors Population Density, Median Income, Elevation, mean Normalized Difference Vegetation Index (NDVI), and mean Normalized Difference Moisture Index (NDMI). Response curves are shown for each predictor when all other predictors are held constant. Mean response from 10 folds is shown with a red line, and +/- one standard deviation is shown in blue. (**F**) Maxent output showing the average change in the Area Under Curve (AUC) value when only that predictor is included (dark blue), when only that predictor is removed (green), and when all predictors were included (red).

**Figure S46.**
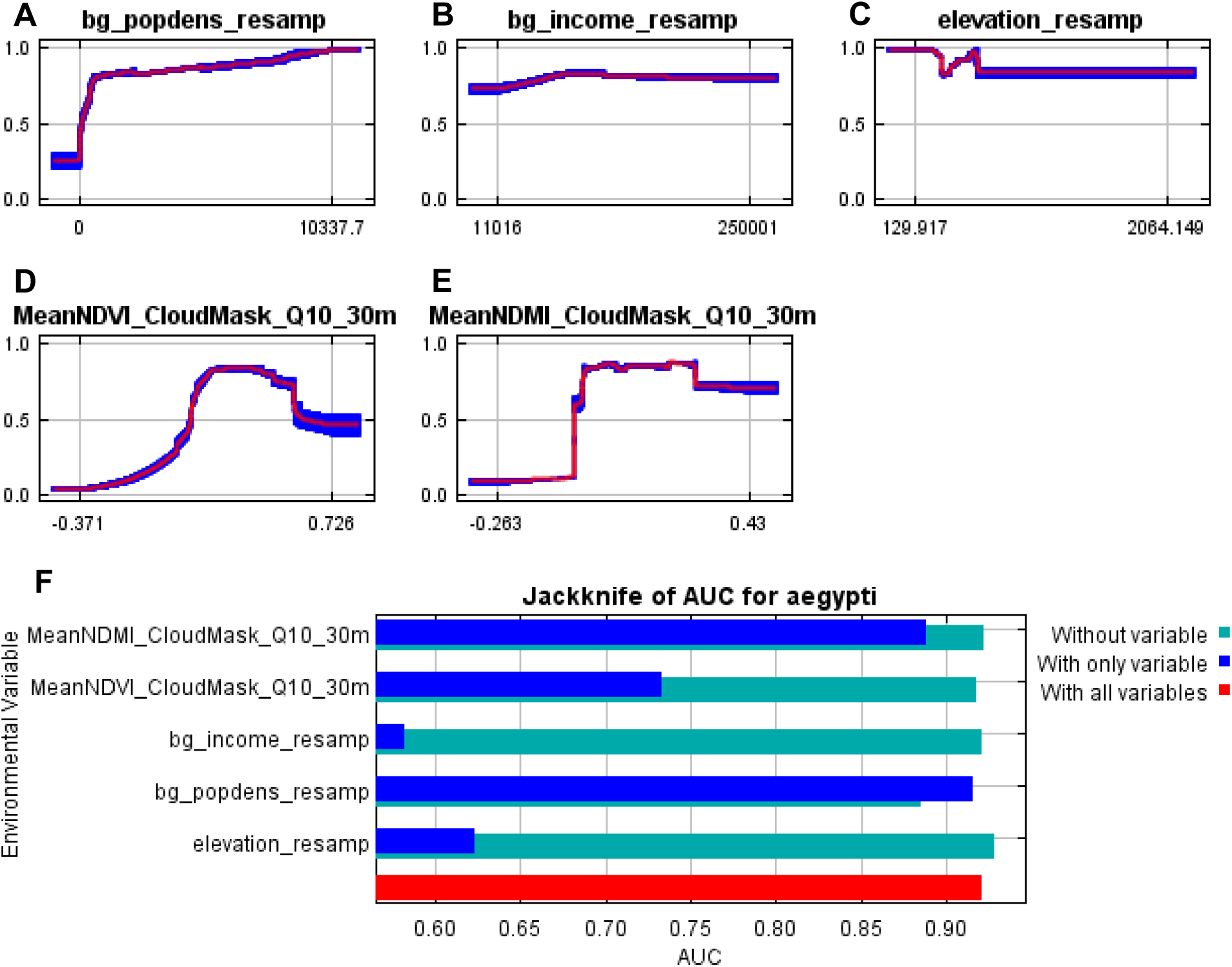
Response curves for Quarter 10 (5/1/16—7/31/16). (**A**) through (**E**) show Maxent response curves for the predictors Population Density, Median Income, Elevation, mean Normalized Difference Vegetation Index (NDVI), and mean Normalized Difference Moisture Index (NDMI). Response curves are shown for each predictor when all other predictors are held constant. Mean response from 10 folds is shown with a red line, and +/- one standard deviation is shown in blue. (**F**) Maxent output showing the average change in the Area Under Curve (AUC) value when only that predictor is included (dark blue), when only that predictor is removed (green), and when all predictors were included (red).

**Figure S47.**
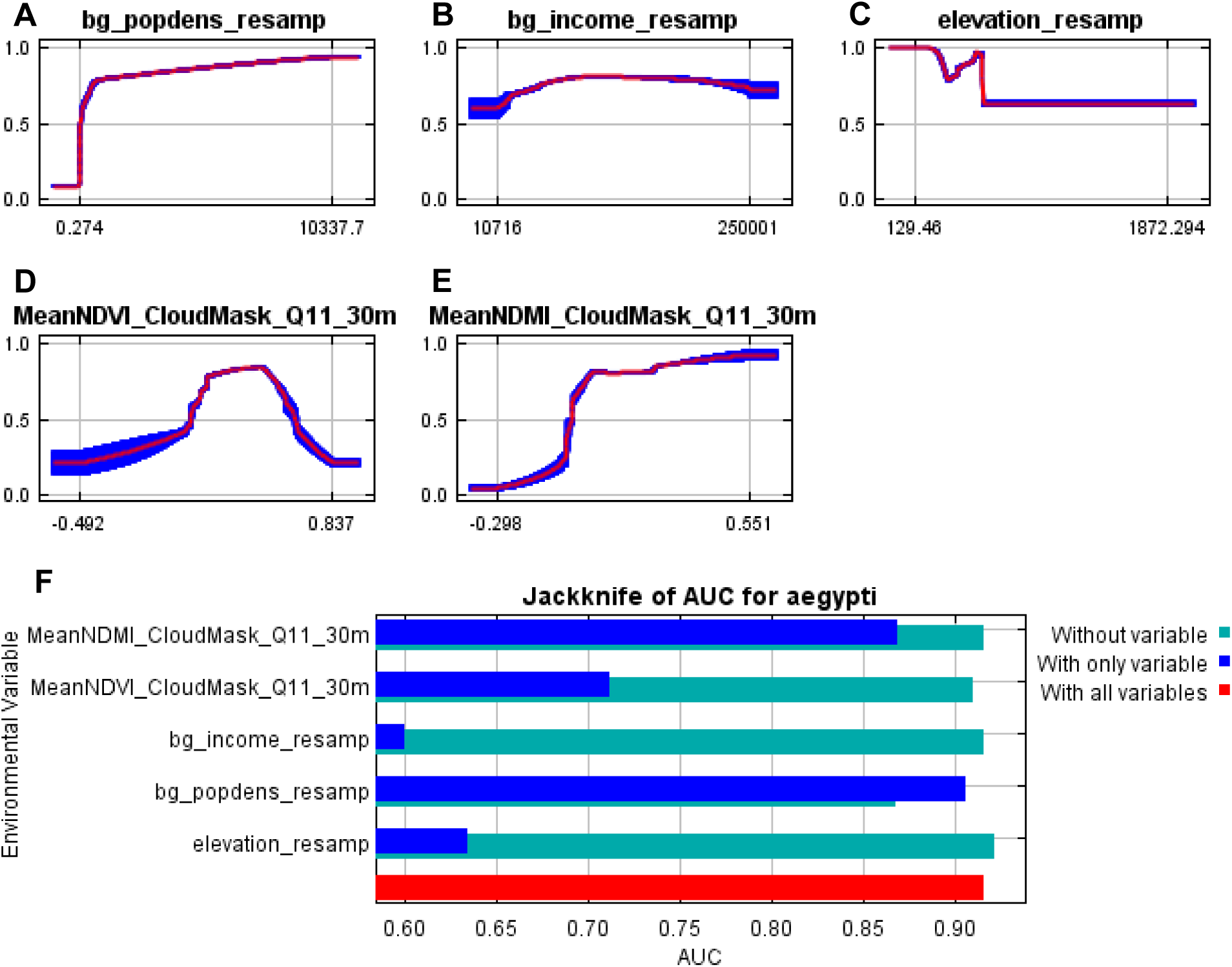
Response curves for Quarter 11 (8/1/16—10/31/16). (**A**) through (**E**) show Maxent response curves for the predictors Population Density, Median Income, Elevation, mean Normalized Difference Vegetation Index (NDVI), and mean Normalized Difference Moisture Index (NDMI). Response curves are shown for each predictor when all other predictors are held constant. Mean response from 10 folds is shown with a red line, and +/- one standard deviation is shown in blue. (**F**) Maxent output showing the average change in the Area Under Curve (AUC) value when only that predictor is included (dark blue), when only that predictor is removed (green), and when all predictors were included (red).

**Figure S48.**
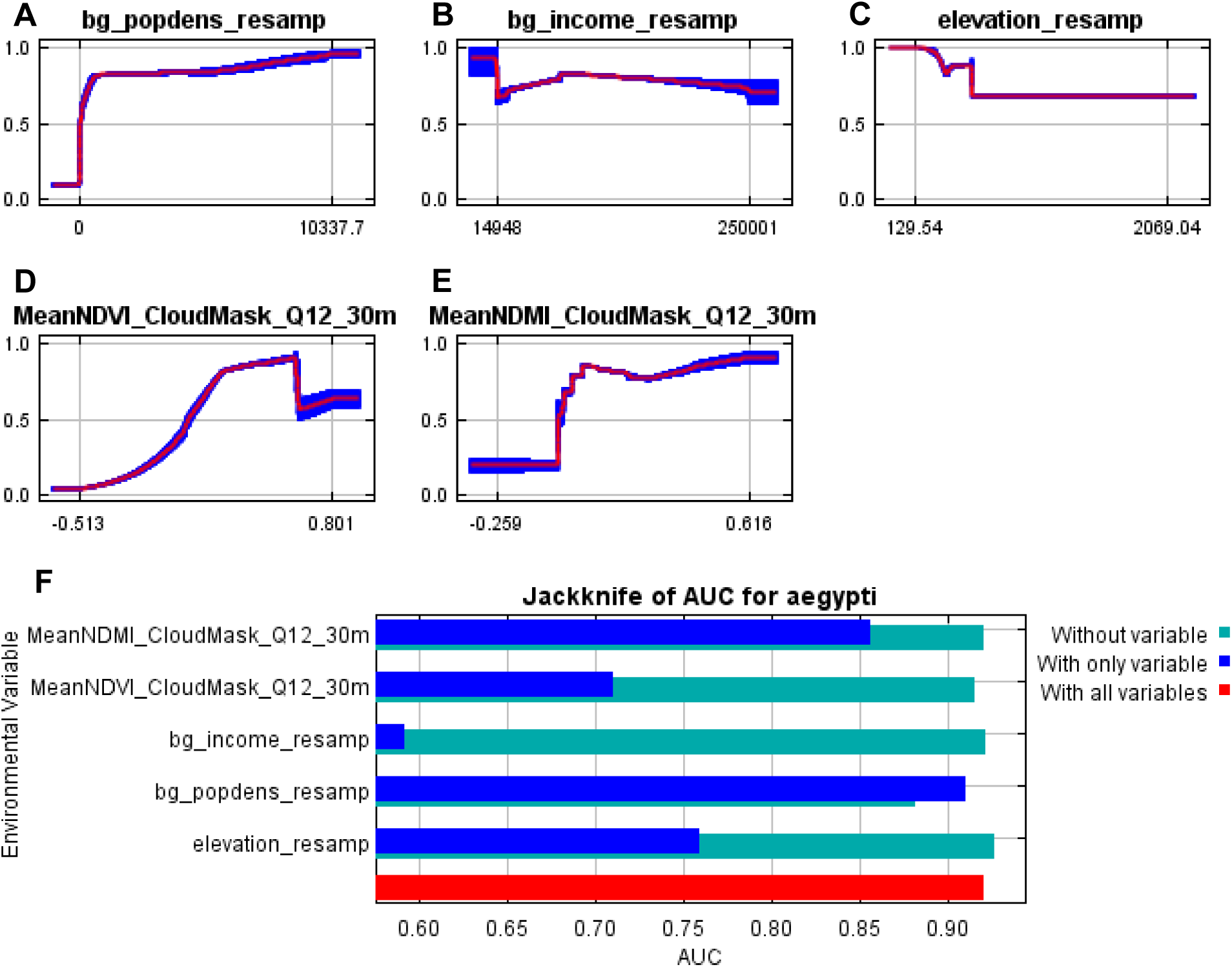
Response curves for Quarter 12 (11/1/16—1/31/17). (**A**) through (**E**) show Maxent response curves for the predictors Population Density, Median Income, Elevation, mean Normalized Difference Vegetation Index (NDVI), and mean Normalized Difference Moisture Index (NDMI). Response curves are shown for each predictor when all other predictors are held constant. Mean response from 10 folds is shown with a red line, and +/- one standard deviation is shown in blue. (**F**) Maxent output showing the average change in the Area Under Curve (AUC) value when only that predictor is included (dark blue), when only that predictor is removed (green), and when all predictors were included (red).

**Figure S49.**
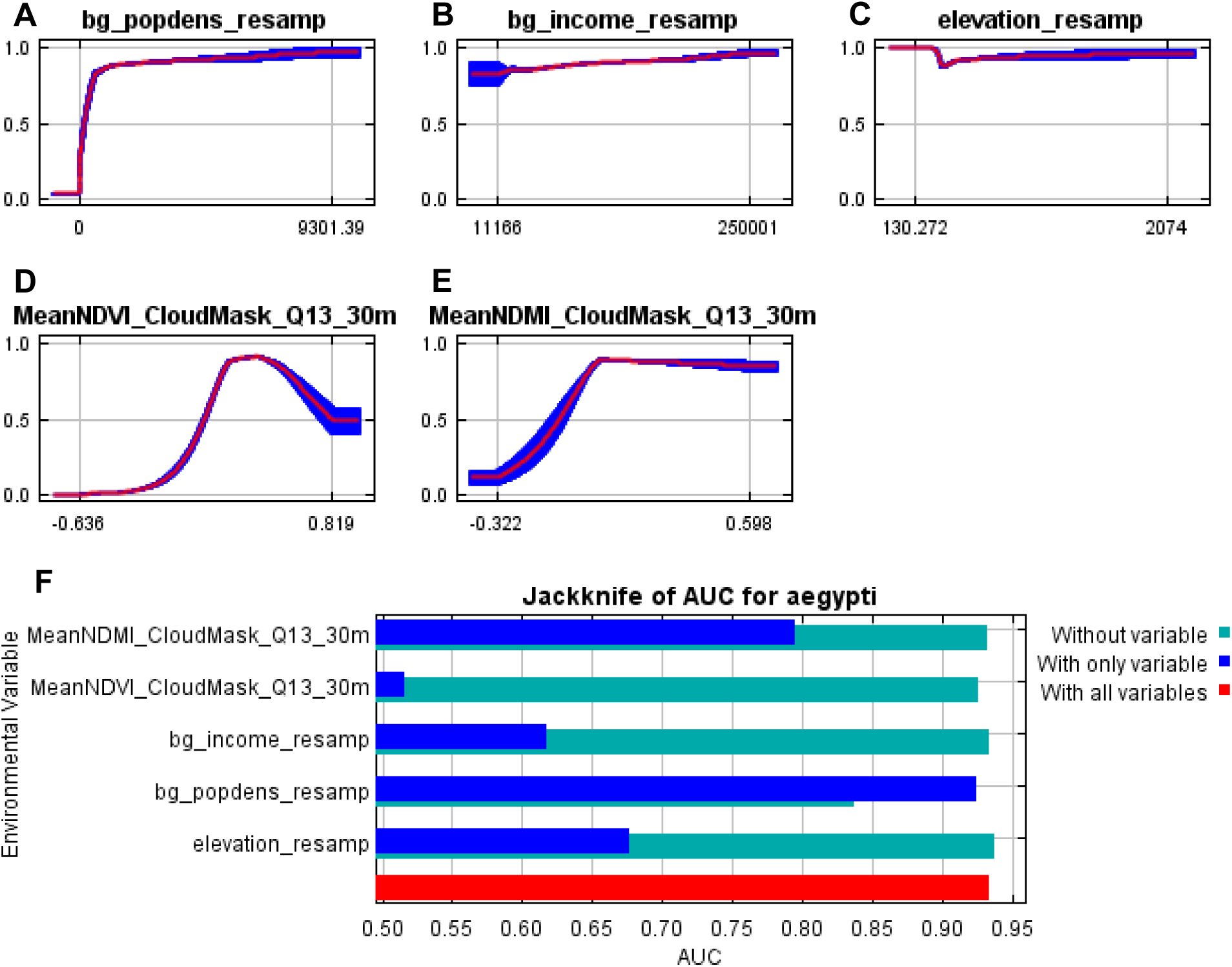
Response curves for Quarter 13 (2/1/17—4/30/17). (**A**) through (**E**) show Maxent response curves for the predictors Population Density, Median Income, Elevation, mean Normalized Difference Vegetation Index (NDVI), and mean Normalized Difference Moisture Index (NDMI). Response curves are shown for each predictor when all other predictors are held constant. Mean response from 10 folds is shown with a red line, and +/- one standard deviation is shown in blue. (**F**) Maxent output showing the average change in the Area Under Curve (AUC) value when only that predictor is included (dark blue), when only that predictor is removed (green), and when all predictors were included (red).

**Figure S50.**
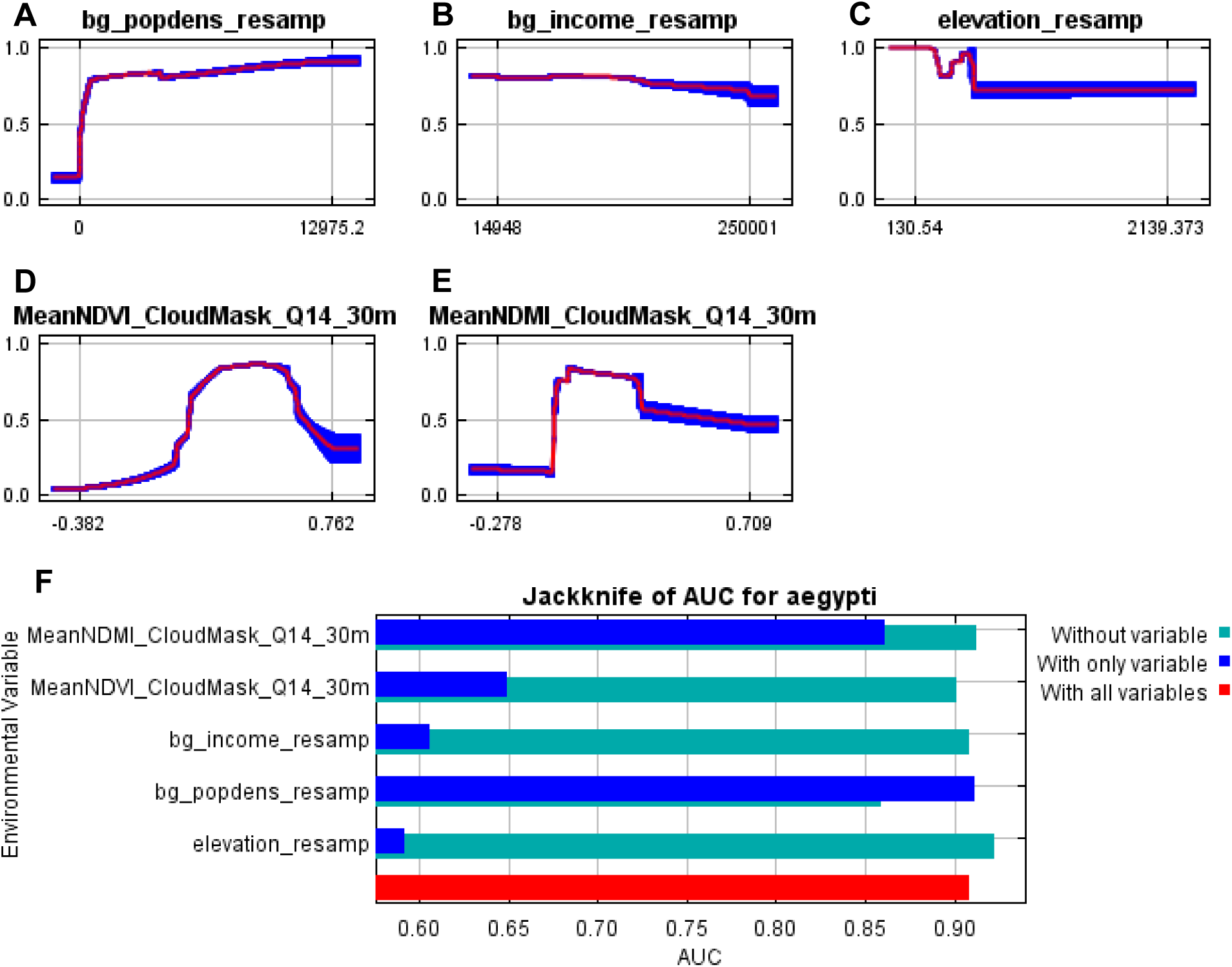
Response curves for Quarter 14 (5/1/17—7/31/17). (**A**) through (**E**) show Maxent response curves for the predictors Population Density, Median Income, Elevation, mean Normalized Difference Vegetation Index (NDVI), and mean Normalized Difference Moisture Index (NDMI). Response curves are shown for each predictor when all other predictors are held constant. Mean response from 10 folds is shown with a red line, and +/- one standard deviation is shown in blue. (**F**) Maxent output showing the average change in the Area Under Curve (AUC) value when only that predictor is included (dark blue), when only that predictor is removed (green), and when all predictors were included (red).

**Figure S51.**
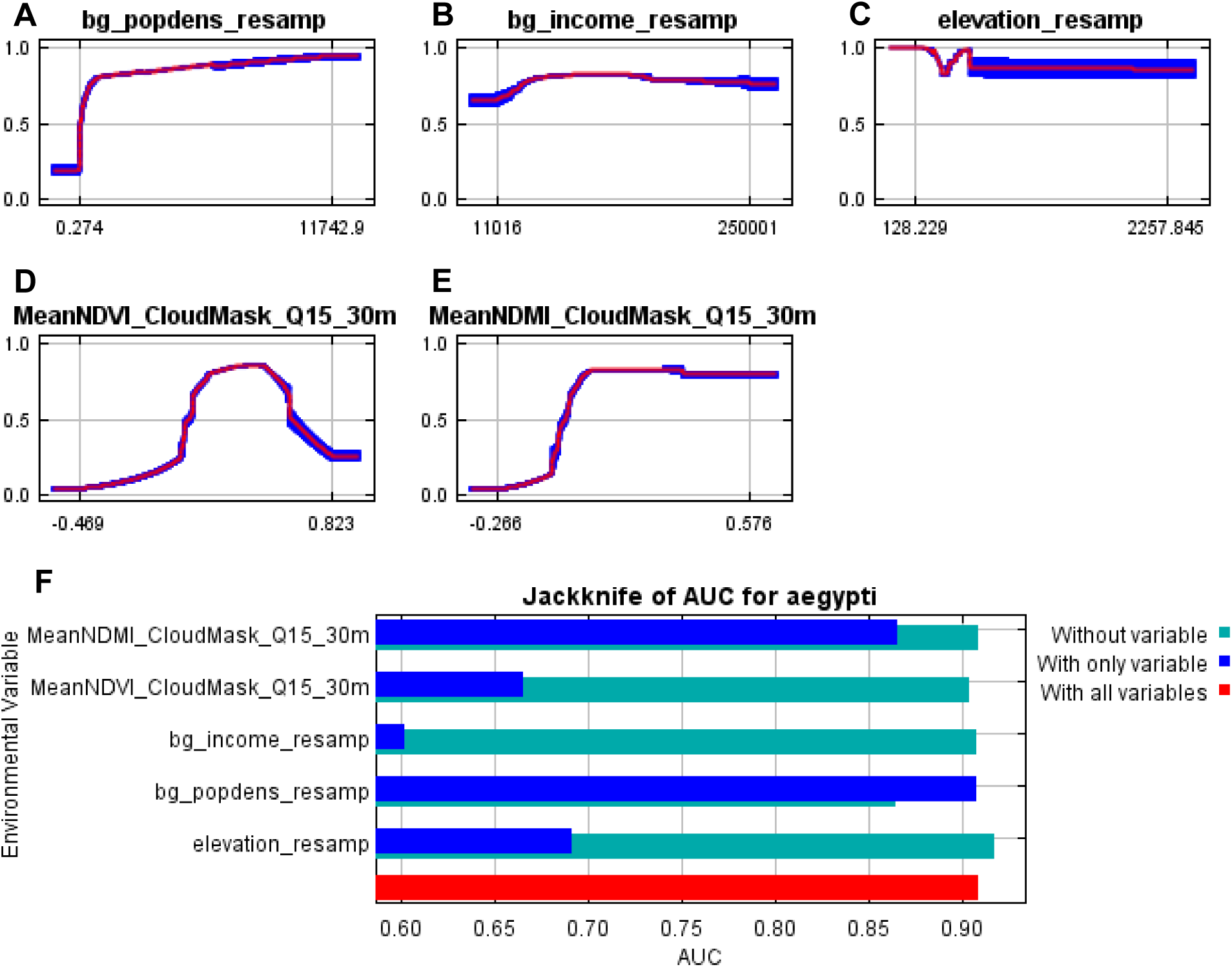
Response curves for Quarter 15 (8/1/17—10/31/17). (**A**) through (**E**) show Maxent response curves for the predictors Population Density, Median Income, Elevation, mean Normalized Difference Vegetation Index (NDVI), and mean Normalized Difference Moisture Index (NDMI). Response curves are shown for each predictor when all other predictors are held constant. Mean response from 10 folds is shown with a red line, and +/- one standard deviation is shown in blue. (**F**) Maxent output showing the average change in the Area Under Curve (AUC) value when only that predictor is included (dark blue), when only that predictor is removed (green), and when all predictors were included (red).

**Figure S52.**
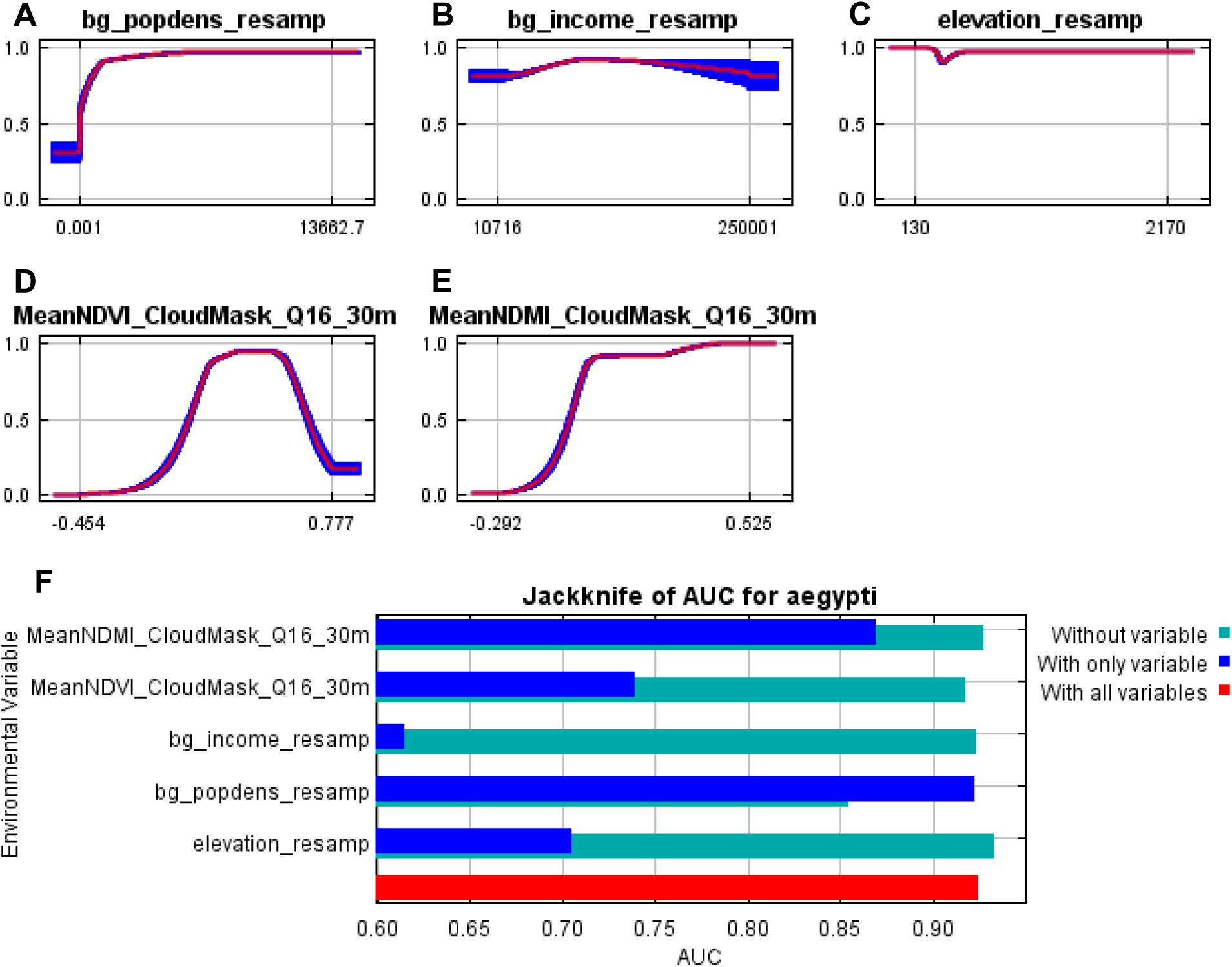
Response curves for Quarter 16 (11/1/17—1/31/18). (**A**) through (**E**) show Maxent response curves for the predictors Population Density, Median Income, Elevation, mean Normalized Difference Vegetation Index (NDVI), and mean Normalized Difference Moisture Index (NDMI). Response curves are shown for each predictor when all other predictors are held constant. Mean response from 10 folds is shown with a red line, and +/- one standard deviation is shown in blue. (**F**) Maxent output showing the average change in the Area Under Curve (AUC) value when only that predictor is included (dark blue), when only that predictor is removed (green), and when all predictors were included (red).

**Figure S53.**
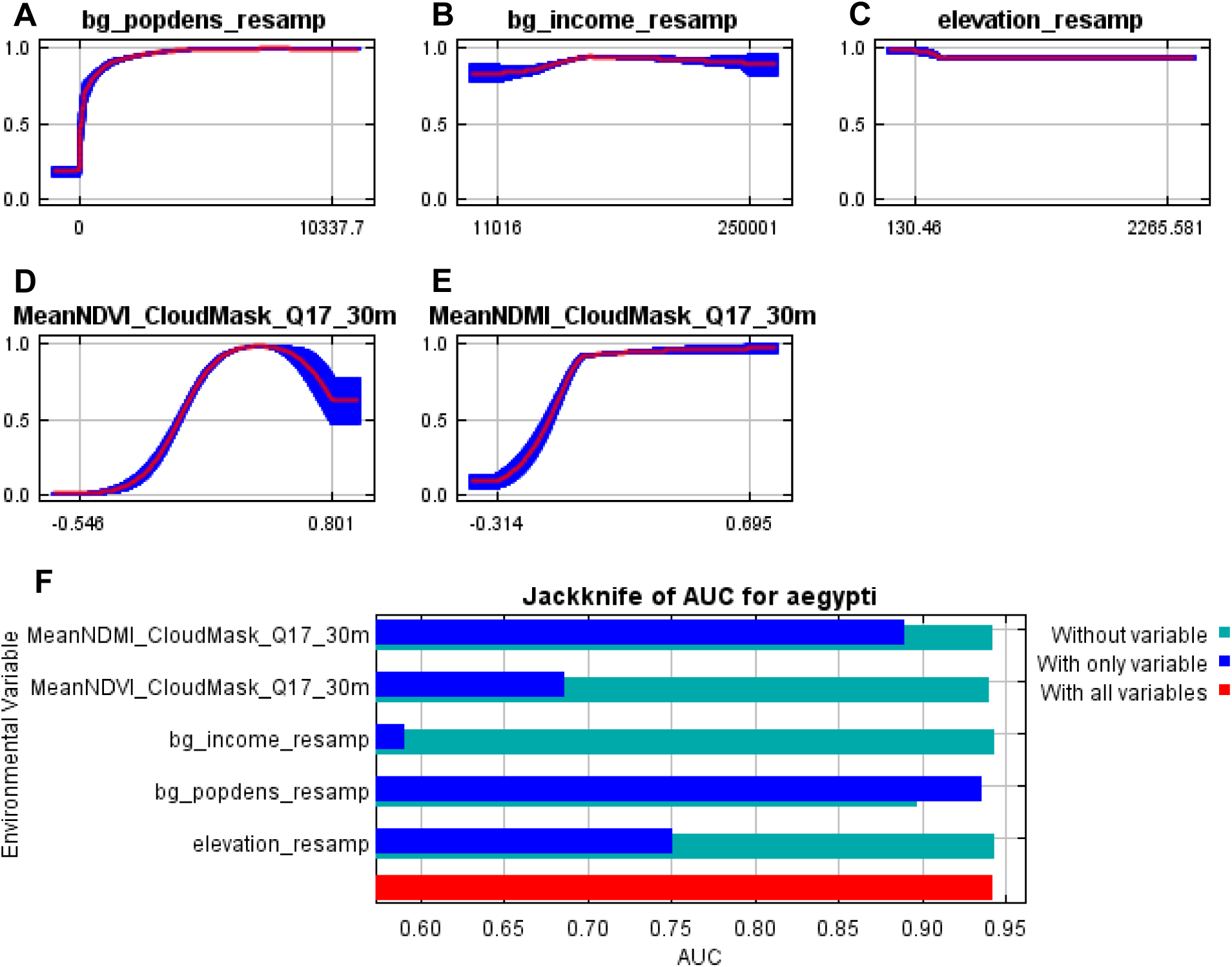
Response curves for Quarter 17 (2/1/18—4/30/18). (**A**) through (**E**) show Maxent response curves for the predictors Population Density, Median Income, Elevation, mean Normalized Difference Vegetation Index (NDVI), and mean Normalized Difference Moisture Index (NDMI). Response curves are shown for each predictor when all other predictors are held constant. Mean response from 10 folds is shown with a red line, and +/- one standard deviation is shown in blue. (**F**) Maxent output showing the average change in the Area Under Curve (AUC) value when only that predictor is included (dark blue), when only that predictor is removed (green), and when all predictors were included (red).

**Figure S54.**
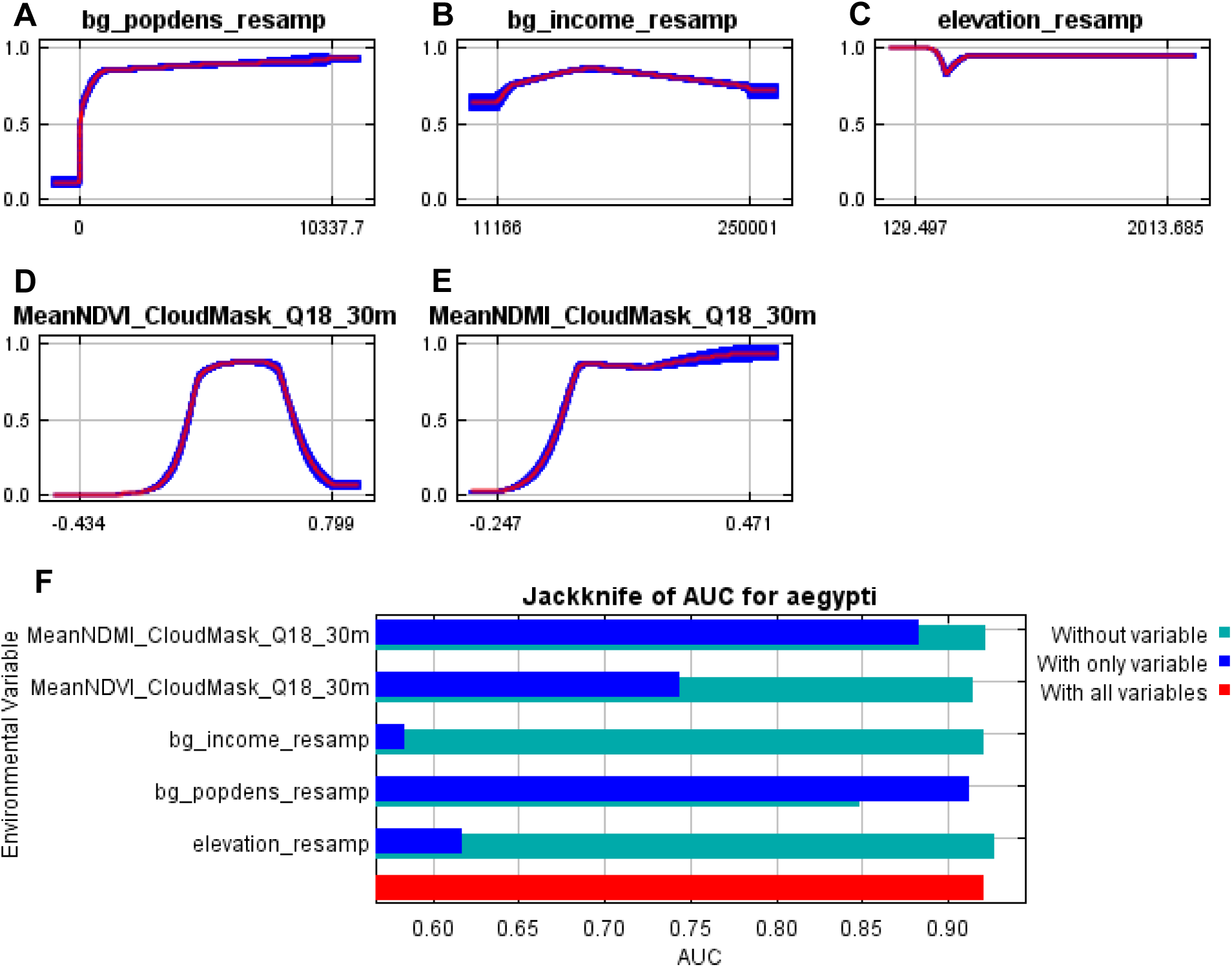
Response curves for Quarter 18 (5/1/18—7/31/18). (**A**) through (**E**) show Maxent response curves for the predictors Population Density, Median Income, Elevation, mean Normalized Difference Vegetation Index (NDVI), and mean Normalized Difference Moisture Index (NDMI). Response curves are shown for each predictor when all other predictors are held constant. Mean response from 10 folds is shown with a red line, and +/- one standard deviation is shown in blue. (**F**) Maxent output showing the average change in the Area Under Curve (AUC) value when only that predictor is included (dark blue), when only that predictor is removed (green), and when all predictors were included (red).

**Figure S55.**
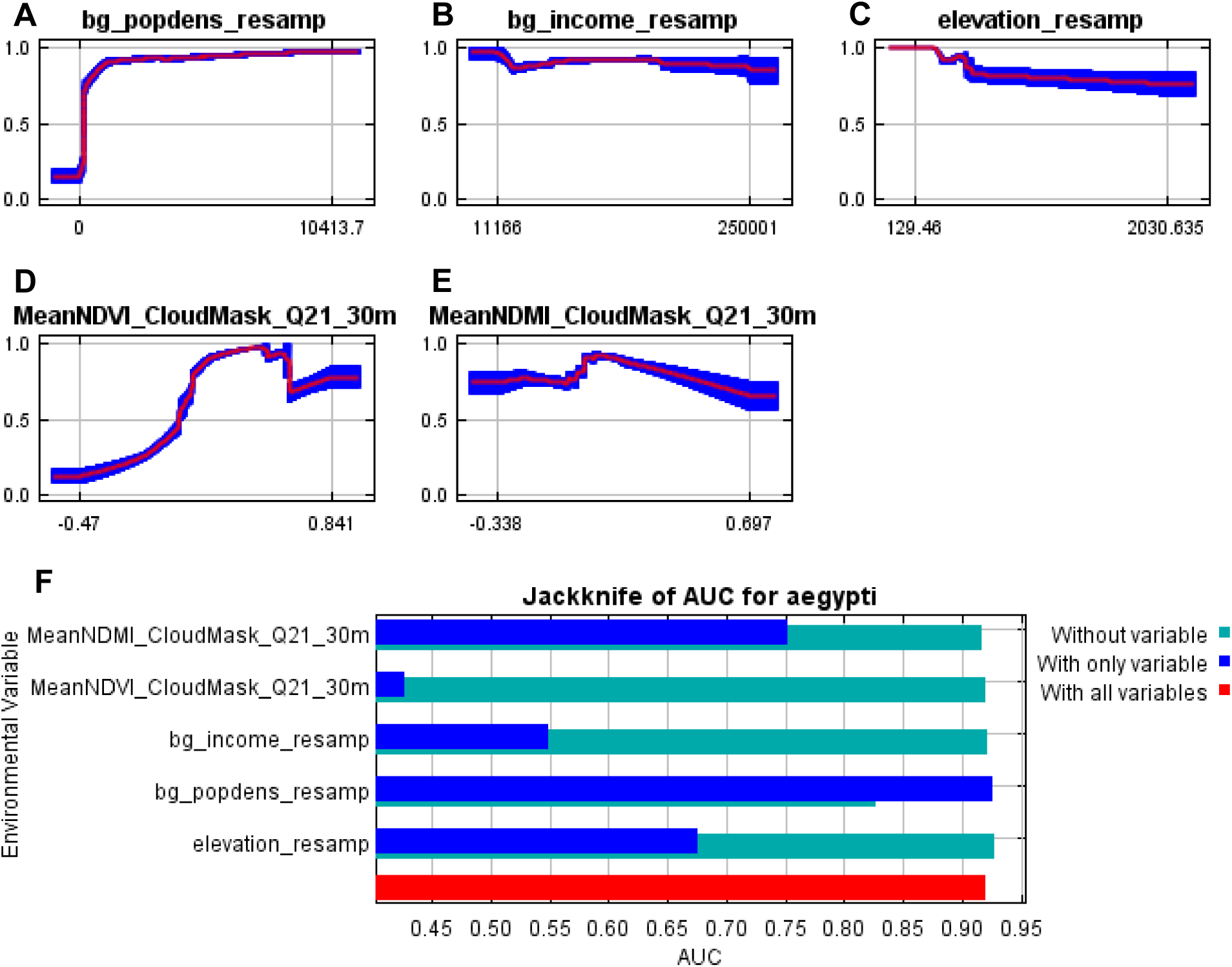
Response curves for Quarter 21 (2/1/19—4/30/19). (**A**) through (**E**) show Maxent response curves for the predictors Population Density, Median Income, Elevation, mean Normalized Difference Vegetation Index (NDVI), and mean Normalized Difference Moisture Index (NDMI). Response curves are shown for each predictor when all other predictors are held constant. Mean response from 10 folds is shown with a red line, and +/- one standard deviation is shown in blue. (**F**) Maxent output showing the average change in the Area Under Curve (AUC) value when only that predictor is included (dark blue), when only that predictor is removed (green), and when all predictors were included (red).

**Figure S56.**
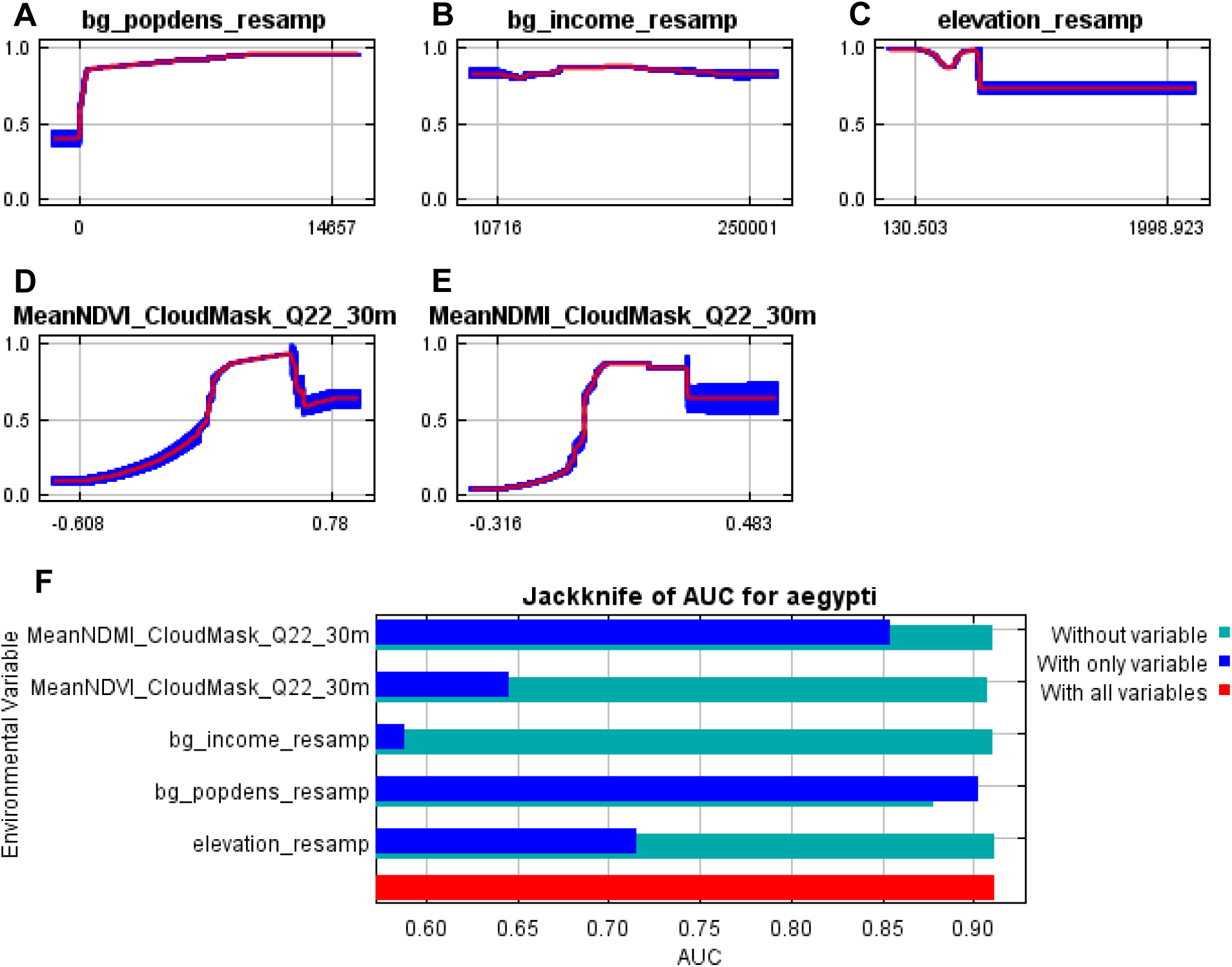
Response curves for Quarter 22 (5/1/19—7/31/19). (**A**) through (**E**) show Maxent response curves for the predictors Population Density, Median Income, Elevation, mean Normalized Difference Vegetation Index (NDVI), and mean Normalized Difference Moisture Index (NDMI). Response curves are shown for each predictor when all other predictors are held constant. Mean response from 10 folds is shown with a red line, and +/- one standard deviation is shown in blue. (**F**) Maxent output showing the average change in the Area Under Curve (AUC) value when only that predictor is included (dark blue), when only that predictor is removed (green), and when all predictors were included (red).

**Figure S57.**
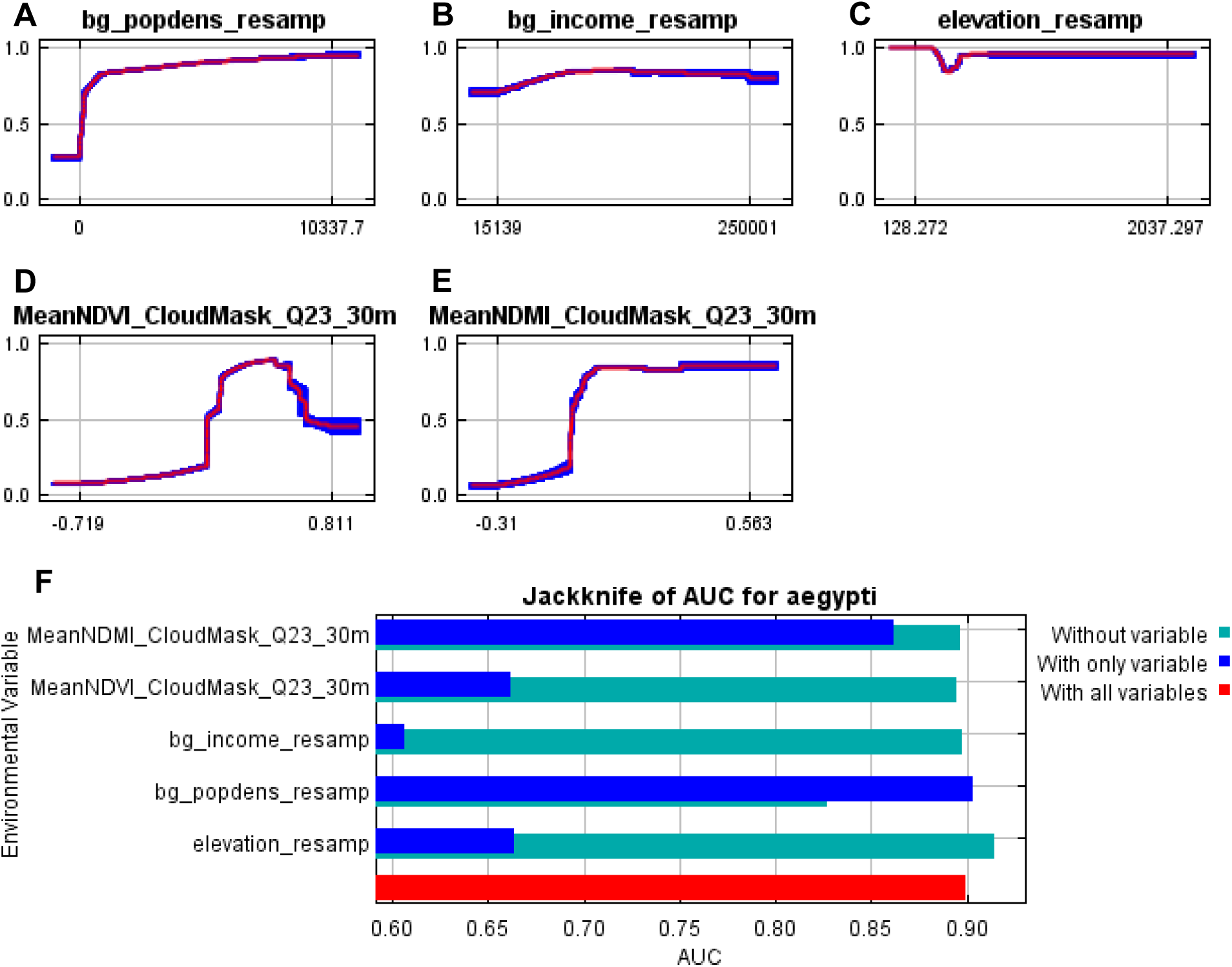
Response curves for Quarter 23 (8/1/19—10/31/19). (**A**) through (**E**) show Maxent response curves for the predictors Population Density, Median Income, Elevation, mean Normalized Difference Vegetation Index (NDVI), and mean Normalized Difference Moisture Index (NDMI). Response curves are shown for each predictor when all other predictors are held constant. Mean response from 10 folds is shown with a red line, and +/- one standard deviation is shown in blue. (**F**) Maxent output showing the average change in the Area Under Curve (AUC) value when only that predictor is included (dark blue), when only that predictor is removed (green), and when all predictors were included (red).

**Figure S58.**
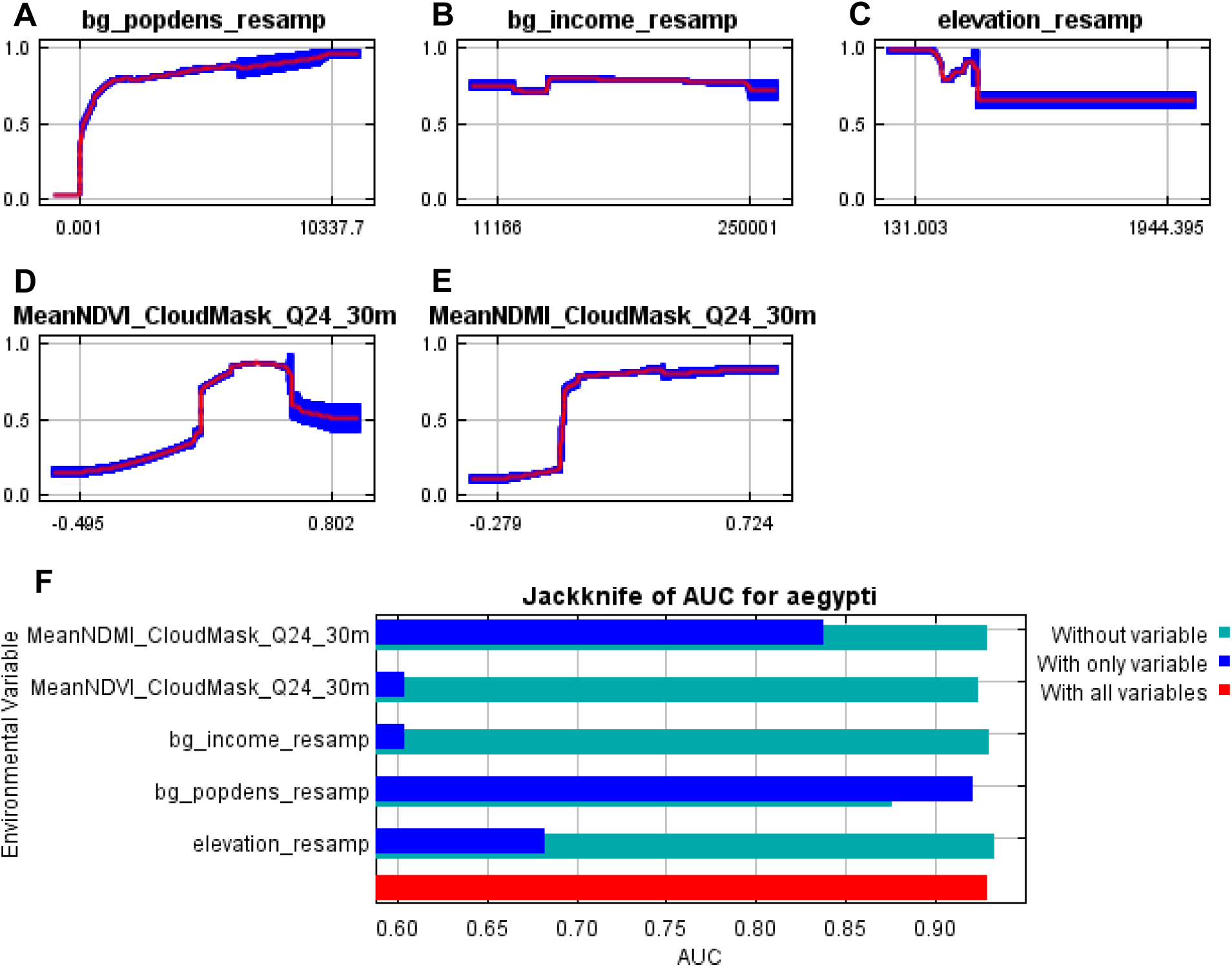
Response curves for Quarter 24 (11/1/19—1/31/20). (**A**) through (**E**) show Maxent response curves for the predictors Population Density, Median Income, Elevation, mean Normalized Difference Vegetation Index (NDVI), and mean Normalized Difference Moisture Index (NDMI). Response curves are shown for each predictor when all other predictors are held constant. Mean response from 10 folds is shown with a red line, and +/- one standard deviation is shown in blue. (**F**) Maxent output showing the average change in the Area Under Curve (AUC) value when only that predictor is included (dark blue), when only that predictor is removed (green), and when all predictors were included (red).

**Figure S59.**
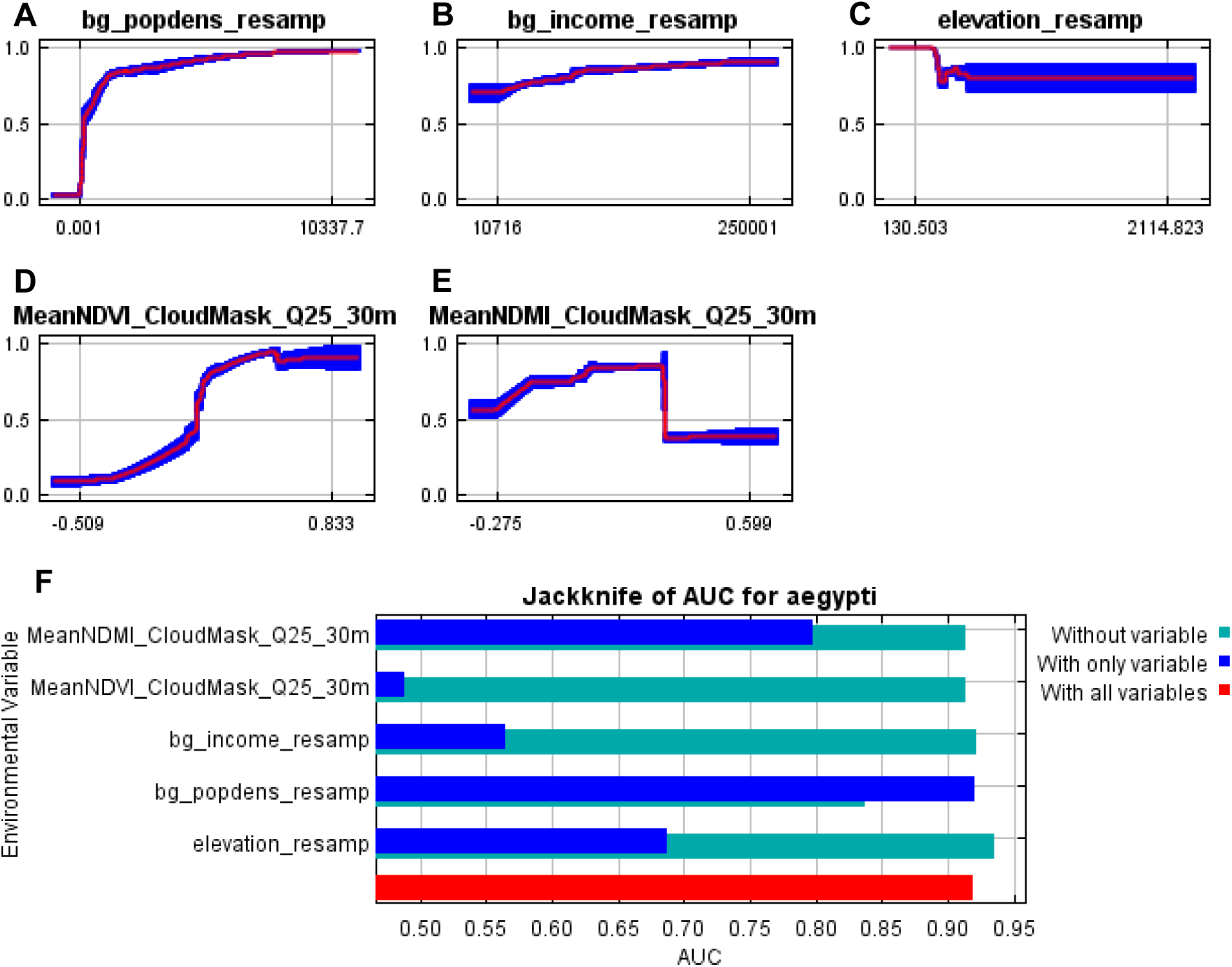
Response curves for Quarter 25 (2/1/20—4/30/20). (**A**) through (**E**) show Maxent response curves for the predictors Population Density, Median Income, Elevation, mean Normalized Difference Vegetation Index (NDVI), and mean Normalized Difference Moisture Index (NDMI). Response curves are shown for each predictor when all other predictors are held constant. Mean response from 10 folds is shown with a red line, and +/- one standard deviation is shown in blue. (**F**) Maxent output showing the average change in the Area Under Curve (AUC) value when only that predictor is included (dark blue), when only that predictor is removed (green), and when all predictors were included (red).

**Figure S60.**
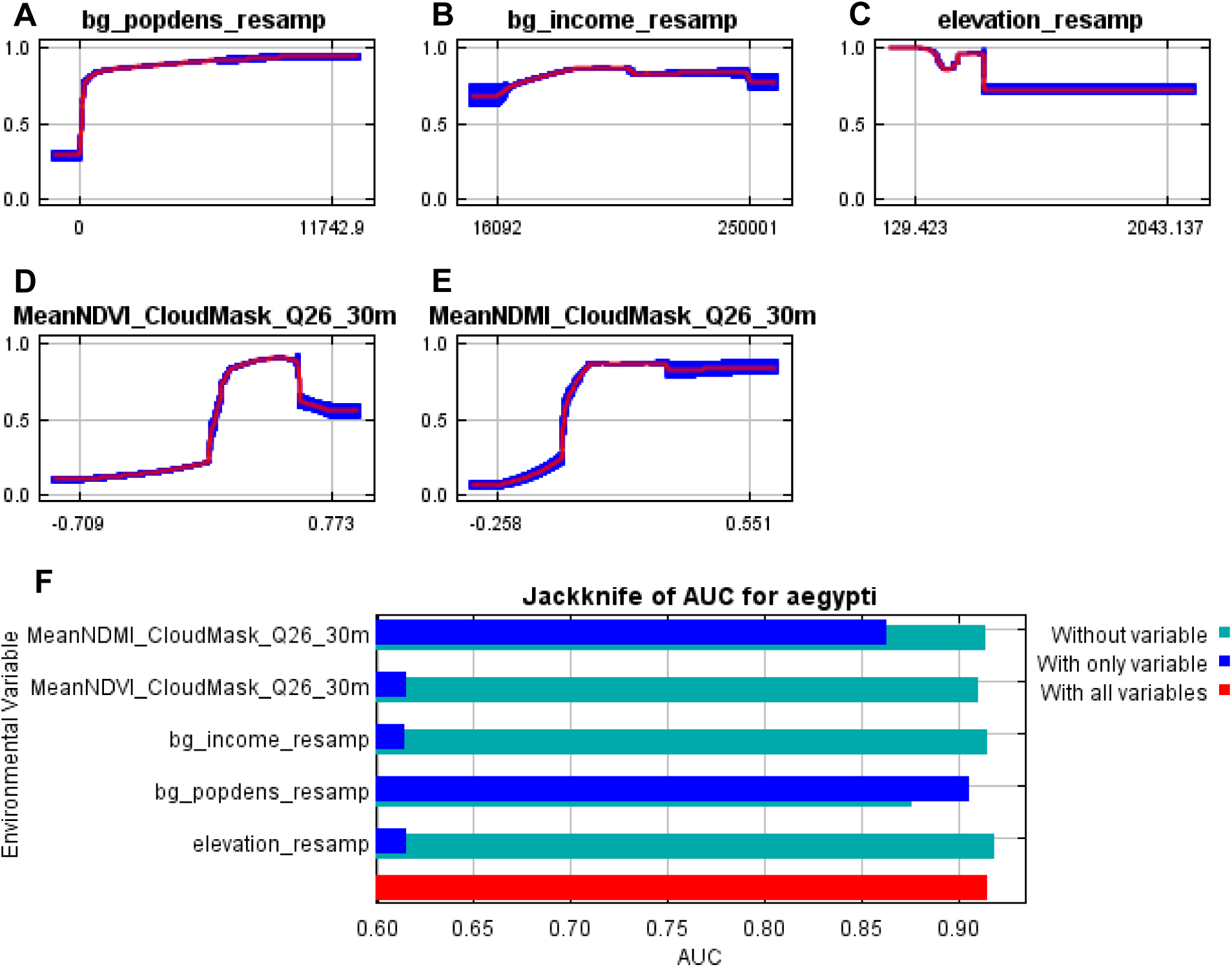
Response curves for Quarter 26 (5/1/20—7/31/20). (**A**) through (**E**) show Maxent response curves for the predictors Population Density, Median Income, Elevation, mean Normalized Difference Vegetation Index (NDVI), and mean Normalized Difference Moisture Index (NDMI). Response curves are shown for each predictor when all other predictors are held constant. Mean response from 10 folds is shown with a red line, and +/- one standard deviation is shown in blue. (**F**) Maxent output showing the average change in the Area Under Curve (AUC) value when only that predictor is included (dark blue), when only that predictor is removed (green), and when all predictors were included (red).

**Figure S61.**
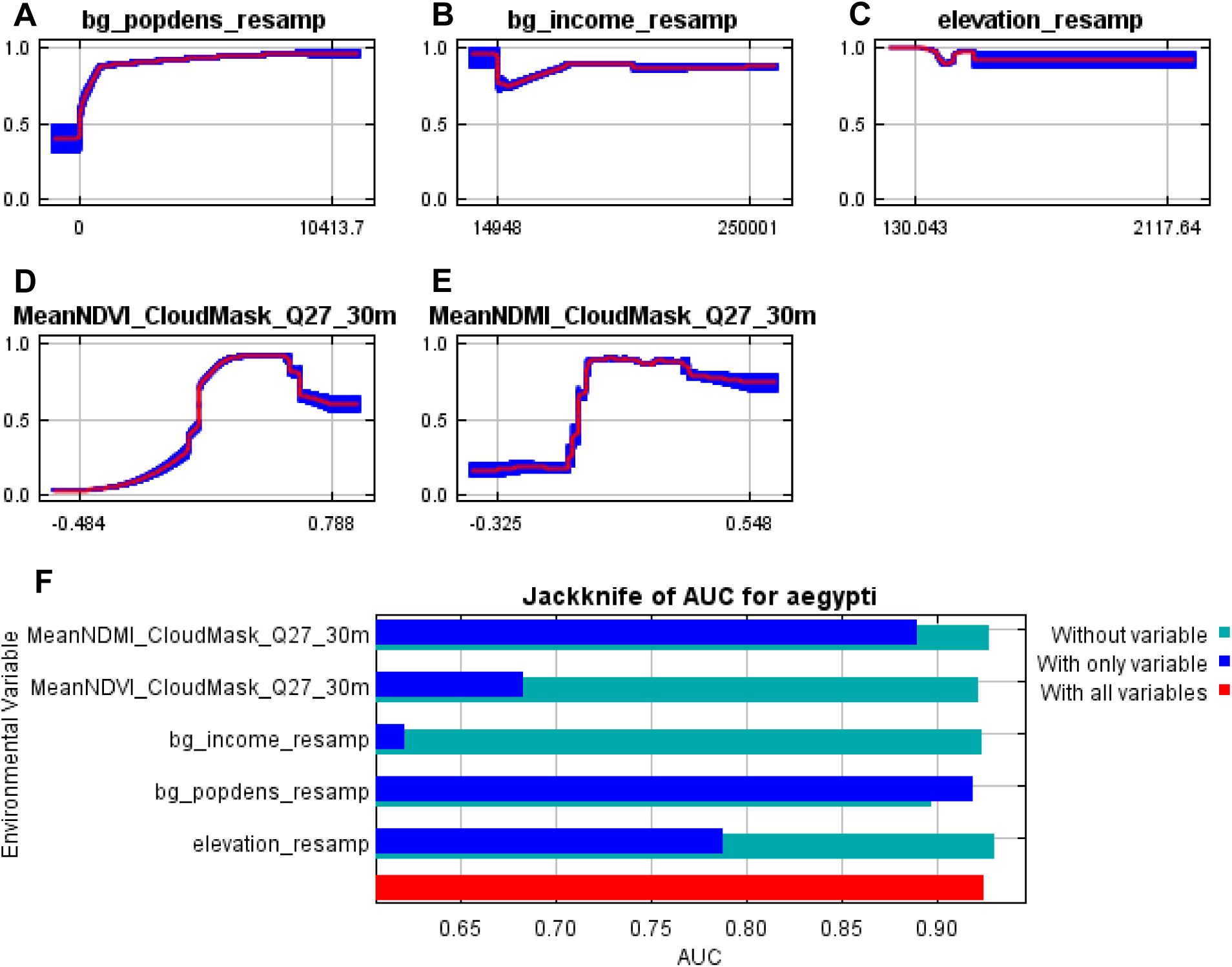
Response curves for Quarter 27 (8/1/20—10/31/20). (**A**) through (**E**) show Maxent response curves for the predictors Population Density, Median Income, Elevation, mean Normalized Difference Vegetation Index (NDVI), and mean Normalized Difference Moisture Index (NDMI). Response curves are shown for each predictor when all other predictors are held constant. Mean response from 10 folds is shown with a red line, and +/- one standard deviation is shown in blue. (**F**) Maxent output showing the average change in the Area Under Curve (AUC) value when only that predictor is included (dark blue), when only that predictor is removed (green), and when all predictors were included (red).

**Table S1.** **Maxent settings used for each quarter.** Abbreviations for feature settings are as follows: L = linear, Q = quadratic, H = hinge, P = product, T = threshold. The feature settings determine the mathematical transformations of the predictors. The regularization multiplier is a penalty that is used to reduce model overfitting.

| Quarter | Feature Settings | Regularization Meter |
| --- | --- | --- |
| 1 | LQHPT | 1 |
| 2 | LQHPT | 1 |
| 3 | LQHPT | 1 |
| 4 | LQHP | 1 |
| 5 | LQHPT | 1 |
| 6 | H | 1 |
| 7 | LQHPT | 1 |
| 8 | LQHPT | 1 |
| 9 | LQHPT | 1 |
| 10 | LQHPT | 1 |
| 11 | LQHPT | 1 |
| 12 | LQHPT | 1 |
| 13 | LQHP | 1 |
| 14 | LQHPT | 1 |
| 15 | LQHPT | 1 |
| 16 | H | 1 |
| 17 | LQH | 1 |
| 18 | LQHP | 1 |
| 19 | n/a | n/a |
| 20 | n/a | n/a |
| 21 | LQHPT | 1 |
| 22 | LQHPT | 1 |
| 23 | LQHPT | 1 |
| 24 | LQHPT | 1 |
| 25 | LQHPT | 1 |
| 26 | LQHPT | 1 |
| 27 | LQHPT | 1 |
| 28 | n/a | n/a |

### Data S1. (separate file)

All data and code used in this analysis is available in a public Google Drive at https://drive.google.com/drive/folders/1Hrqs-Vpv-9w64rtBBXuKq9Po51D3A3-G?usp=sharing. All descriptions of .R code, as well as an intended file structure and workflow can be found in the README folder. Data provided for each quarter are the observations of *Ae. aegypti* in Maricopa County. Latitude and longitude values have been truncated to the hundredths digit and jittered by adding a random number between [-0.01, 0.01] to address fears of trap theft or destruction.

